# Cucurbit[7]uril Enhances Distance Measurements of Spin-Labeled Proteins

**DOI:** 10.1101/2023.08.22.554361

**Authors:** Zhimin Yang, Richard A. Stein, Maren Pink, Peter Madzelan, Thacien Ngendahimana, Suchada Rajca, Mark A. Wilson, Sandra S. Eaton, Gareth R. Eaton, Hassane S. Mchaourab, Andrzej Rajca

**Affiliations:** Department of Chemistry, University of Nebraska, Lincoln, Nebraska 68588-0304, United States; Department of Molecular Physiology and Biophysics, Vanderbilt University, Nashville, Tennessee 37232, United States; IUMSC, Department of Chemistry, Indiana University, Bloomington, Indiana 47405-7102, United States; Department of Biochemistry and Redox Biology Center, University of Nebraska, Lincoln, Nebraska 68588-0304, United States; Department of Chemistry and Biochemistry, University of Denver, Denver, Colorado 80208, United States

**Keywords:** Host-guest systems, Cucurbiturils, EPR spectroscopy, Biophysics, Organic Radicals

## Abstract

We report complex formation between the chloroacetamide 2,6-diazaadamantane nitroxide radical (ClA-DZD) and cucurbit[7]uril (CB-7), for which the association constant in water, *K*_a_ = 1.9 × 10^6^ M^-1^, is at least one order of magnitude higher than the previously studied organic radicals. The radical is highly immobilized by CB-7, as indicated by the increase of the rotational correlation time, *τ*_rot_, by a factor of 36, relative to that in the buffer solution. The X-ray structure of ClA-DZD@CB-7 shows the encapsulated DZD guest inside the undistorted CB-7 host, with the pendant group protruding outside. Upon addition of CB-7 to T4 Lysozyme (T4L) doubly spin-labeled with the iodoacetamide derivative of DZD, we observe the increase in *τ*_rot_ and electron spin coherence time, *T*_m_, along with the narrowing of inter-spin distance distributions. Sensitivity of the DEER measurements at 83 K increases by a factor 4 – 9, compared to the common spin label such as MTSL, which is not affected by CB-7. Inter-spin distances of 3-nm could be reliably measured in water/glycerol up to temperatures near the glass transition/melting temperature of the matrix at 200 K, thus bringing us closer to the goal of supramolecular recognition-enabled long-distance DEER measurements at near physiological temperatures. The X-ray structure of DZD-T4L 65 at 1.12 Å resolution allows for unambiguous modeling of the DZD label (0.88 occupancy), indicating undisturbed structure and conformation of the protein.

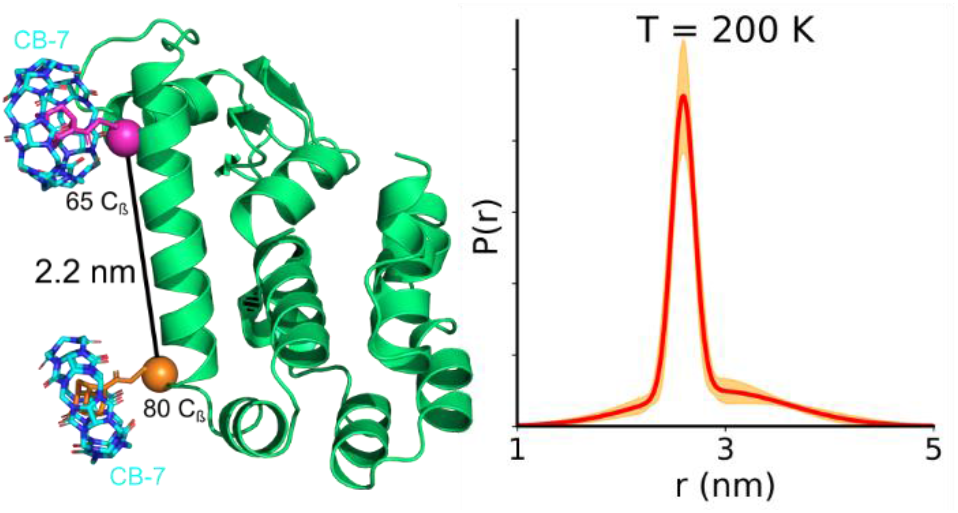

## INTRODUCTION

Tremendous efforts have been made to probe macromolecules in natural biological systems that would provide insight into their structures and functions. Site-directed spin labeling (SDSL) pulsed electron paramagnetic resonance (EPR) dipolar spectroscopy (PDS), which includes double electron–electron resonance (DEER) and double quantum coherence (DQC), is among the best techniques for accurate measurement of the conformations of flexible regions of biomolecules.^1-10^ The advantages include high sensitivity, wide dynamic range, absolute distance distributions, and minimal structural perturbation from relatively small-size nitroxide spin labels.^2,3^ The most widely used spin label for SDSL is methane thiosulfonate pyrolline nitroxide (MTSL),^11^ which selectively binds to cysteine, that can be introduced at selected locations by mutagenesis to provide doubly-labeled proteins.^1,2^ Using nitroxides, these measurements require cryogenic temperatures because of the dynamic averaging effects associated with methyl group rotation, which shorten the electron spin coherence time (*T*_m_) between about 80 and 300 K.^12-15^

Efforts have long been made with limited success to raise the experimental temperature range for the measurements. A promising approach to DQC distance measurements in the physiological temperature range was demonstrated using a spin label based on a carbon-centered trityl radical and immobilized T4 lysozyme (T4L) or nucleic acids.^16,17^ The inherent disadvantage of the trityl label is the lengthy synthesis, physical size, and hydrophobicity, and the distances can only be measured by the less commonly available DQC method,^4,18^ because the spectral width, even at Q-band, is too narrow for DEER, unless the trityl label is ^13^C substituted.^19,20^ Distance measurement up to 4 and 5 nm at physiological temperature was demonstrated by saturation recovery EPR spectroscopy of protein labeled with a nitroxide or a trityl spin label which have long electron spin–lattice relaxation times (*T*_1_) and fast relaxing metal ion, such as Cu^2+^.^21-23^ Notably, the experiment requires insertion of the 5-residue, GGGHG, Cu^2+-^binding loop into the protein sequence.

We have developed a new generation of nitroxide spin labels devoid of methyl groups around the N-O moiety, such as IA-Spiro^24^ and a compact iodoacetamide diazaadamantane (IA-DZD) spin label (Figure 1),^25^ that can provide DEER measurements with improved sensitvity. T4L doubly spin-labeled with IA-DZD possesses sufficiently long *T*_m_ to allow for measurement of distances of up to 4 – 5 nm at 150 K and, with a sensitivity that is superior to MTSL, at 80 K.

**Figure 1.**
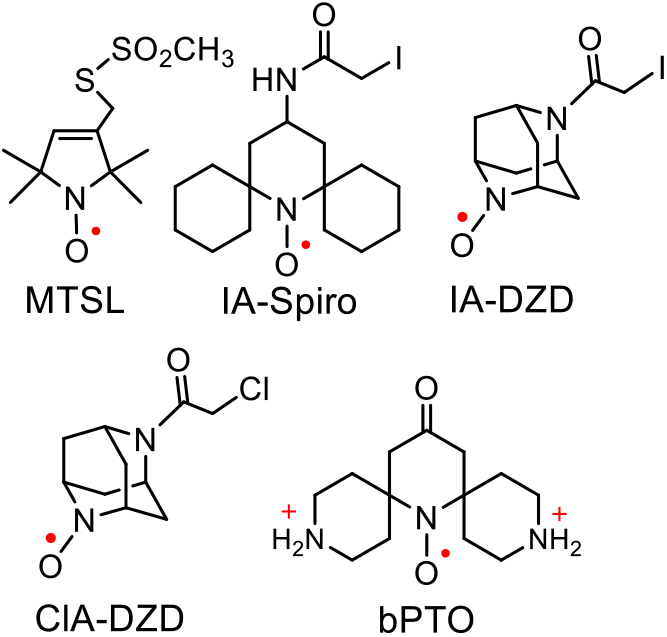
Structures of spin labels, model nitroxide ClA-DZD, and dicationic nitroxide bPTO.

Adamantane and its derivatives are a notable family of molecular guests for supramolecular complexation with macrocyclic cages. We were enticed by the strong affinity of adamantane and intrigued to explore the possibility of employing supramolecular chemistry to facilitate long-distance DEER measurements at near physiological temperatures. We have examined the host-guest supramolecular complexation of IA-DZD and β-cyclodextrin (β-CD), in which we observed the slightly elongated *T*_m_ that further enhanced the sensitivity of DEER distance measurements.^25^ These results inspire us to pursue further extraordinary s pramolecular hosts for superior complexation with adamantane nitroxides.

Cucurbit[n]urils (CB-n) is a family of macrocylic cages with exceptional recognition properties in aqueous medium that have been widely investigated for its applications in drug delivery system and advanced functional materials.^26,27^ Among the CB-n (*n* = 6 – 8) family, CB-7 is the most water soluble host and has the strongest association constants, *K*_a_ = 7.2 × 10^17^ M^−1^ in D_2_O, with selected diamagnetic dicationic guests.^28-31^ However, nitroxides as guests are far behind,^32-34^ with the reports of strongest *K*_a_ ≈ 2 × 10^5^ M^-1^ for dicationic nitroxide radical **bPTO** (Figure 1)^35^ and *K*_a_ = 2.8 × 10^6^ M^−1^ for a TEMPO-cobaltocenium derivative, based on inclusion of cobaltocenium moiety within CB-7.^36^ Nevertheless, the typical association constants for neutral adamantane derivatives with CB-7 are several orders magnitude higher than those with β-CD,^26,37^ thus much tighter complexes with superb association constant may be expected for the DZD nitroxide and CB-7 than for β-CD. Thus, we envisage narrow distance distributions and improved sensitivity of DEER distance measurement for the spin labeled DZD-T4L in the presence of CB-7, when the spin labels are at solvent exposed sites.

Herein we report the complex formation between the diazaadamantane nitroxide radical (DZD) and CB-7, for which the association constant in water is at least one order of magnitude higher than for previously studied organic radicals. The complexed DZD is highly immobilized by CB-7, as indicated by the increase of the rotational correlation time (*τ*_rot_) by a factor of 36, compared to that in buffer solution. The X-ray structure of the ClA-DZD@CB-7 complex clearly shows the encapsulated DZD guest inside the undistorted CB-7 host, with the pendant group protruding outside. Addition of CB-7 to T4L doubly spin-labeled with IA-DZD (DZD-T4L) increases both the *τ*_rot_ of the DZD spin label and its electron spin coherence time, *T*_m_, and importantly, narrows the inter-spin distance distributions. Sensitivity of the DEER measurements at 83 K increases by a factor 4 – 9, compared to the common spin label such as MTSL, which is not affected by CB-7. In addition, inter-spin distances of 3-nm could be reliably measured in glycerol/water up to a temperature of 200 K, which is near the glass transition/melting temperature of the matrix, thus bringing us closer to the goal of supramolecular recognition-enabled long-distance DEER measurements at near physiological temperatures. We also report the crystallographic characterization of DZD-T4L 65 at 1.12 Å resolution that allows for unambiguous modeling of the DZD label (0.88 occupancy) to illustrate that the DZD label leaves intact both structure and conformation of the protein. These results are advances in the supramolecular approach to the DEER distance measurements and lay a foundation for potential applications to probe macromolecules in natural biological systems at physiological temperatures.

## RESULTS AND DISCUSSION

### 1. Complexation of ClA-DZD with CB-7

Our objective is to explore the extent of CB-7 complexation with DZD spin labeled protein to gain insight into the potential benefit of supramolecular recognition in DEER distance measurements. ClA-DZD, which is a synthetic precursor to IA-DZD,^25^ is less hydrophobic than IA-DZD, and thus it is expected to resemble the DZD-linked-to-Cys on protein.

#### 1.1. Isothermal calorimetry (ITC)

Nitroxide radical ClA-DZD shows the highest association constant, *K*_a_ = 1.9 × 10^6^ M^-1^, in water among supramolecular complexes of neutral and charged nitroxide radicals with CB-7 (Table 1).^35,38^ Notably, this value of *K*_a_ is larger by a factor of 2500, compared to that previously reported for the ClA-DZD@β-CD complex,^25^ affirming the superior binding between DZD and CB-7.

**Table 1.** Isothermal Calorimetry: Association Constants (*K*_a_) of ClA-DZD with CB-7 and β-CD.

| Host | Medium | $\text{Na}^+$ (mM) | pH | $K_a \times 10^5 (\text{M}^{-1})$ |
| --- | --- | --- | --- | --- |
| CB-7 | water | 0 | ~7 | $19.3 \pm 0.8$ |
| | phosphate<sup>a</sup> | 69 | 7.48 | $33.7 \pm 1.1$ |
| | Tris-MOPS-1<sup>b</sup> | 53 | 7.24 | $3.74 \pm 0.07$ |
| | Tris-MOPS-2<sup>b</sup> | 3 | ~7 | $0.432 \pm 0.018$ |
| | Tris<sup>b,c</sup> | 0 | 7.26 | $1.03 \pm 0.14$ |
| β-CD | water | 0 | ~7 | $(7.55 \pm 0.07) \times 10^{-3}$ |

<sup>a</sup> 50 mM Phosphate. <sup>b</sup> All 6 mM Tris buffers contain 0.01 mM EDTA; in addition, tris-MOPS-1 and -2 contain 9 mM MOPS, 3-(*N*-morpholino)propane-1-sulfonic acid and 0.02% (by weight) $\text{NaN}_3$ ; tris-MOPS-1 has 50 mM $\text{NaCl}$ . <sup>c</sup> pH is adjusted using 2.73 mM Tris and 3.27 mM Tris hydrochloride. Further details may be found in the SI: Table S4 and Figs. S7–S13.

In phosphate buffer containing 69 mM Na^+^, we observe *K*_a_ = 3.4 × 10^6^ M^-1^, which is 75% higher than that in pure water. The increase in *K*_a_ might suggest *cooperative binding* of sodium cations to the NO moiety of DZD in the complex.^39^ Such binding may be facilitated by the relatively large partial negative charge on the oxygen atom of the highly pyramidal NO moiety of DZD nitroxide.^25^ This observation is contrary to the reports of *competitive binding* of alkali metal cations with supramolecular guests,^40,41^ including nitroxide radicals.^38,42^ In buffer containing tris(hydroxymethyl)amino-methane (Tris) and 3-(*N*-morpholino)propane sulfonic acid (MOPS), *K*_a_ is lower than in water or phosphate buffer. For example, ITC measurements give *K*_a_ = 3.7 × 10^5^ M^-1^ in Tris-MOPS 1, containing 53 mM Na^+^ and 9 mM MOPS, and *K*^a^ = 1.0 × 10^5^ M^-1^ in Tris buffer, without MOPS and Na^+^ (Table 1).

#### 1.2. X-ray crystallography

CB-7 has a propensity to form supramolecular gels in aqueous solution,^43,44^ and thus it is difficult to crystallize. However, there are numerous examples of crystalline inclusion complexes of CB-7 with neutral and cationic guests.^45-47^ Notably, to our knowledge, only a few single crystal X-ray structures of nitroxide (mostly TEMPO-like) complexes with CB-8 have been reported^35,36,48-50^ and there are only two reports on nitroxide (TEMPO-like and nitronyl nitroxide) complexes with CB-7.^50,51^

Crystals of ClA-DZD@CB-7 complex are obtained by slow solvent evaporation from a 1:1 molar mixture of ClA-DZD and CB-7 in water. A complete set of X-ray diffraction data to a resolution of 0.84 Å is collected. The structure reveals the formation of a 1:1 complex of ClA-DZD with CB-7. The presence of nitroxide radicals inside the CB-7 host is evidenced by the average N-O bond length of 1.296 Å (Table S2, SI), which is comparable to 1.2843(19) Å found in IA-DZD spin label.^25^ The asymmetric unit consists of four independent ClA-DZD@CB-7 supramolecules and at least 41 molecules of co-crystallized water molecules. All the water molecules are positioned outside the CB-7 cavity, which is consistent with large value of *K*_a_ observed by ITC (Table 1). In one of the supramolecules, the ClA-DZD guest is not disordered; however, in the other three supramolecules, the host is disordered over two or three positions (Figs. S2 and S3, SI). As illustrated in Figure 2A&B, the ClA-DZD molecule is approximately placed at the center of the near-perfect 7-fold symmetric host, suggesting a good size match between the guest and host. In contrast, in the X-ray structure of a TEMPO derivative, the CB-7 cavity is significantly distorted from 7-fold symmetry.^50^ We find 10 close contacts ≤ sum of vdW radii, predominantly non-classical H-bonds, between the guest and the host (Fig. S2, SI); this is consistent with a near perfect fit of the DZD nitroxide with the CB-7 cavity. The pendant ClA-unit is exposed outside the portals of CB-7.

**Figure 2.**
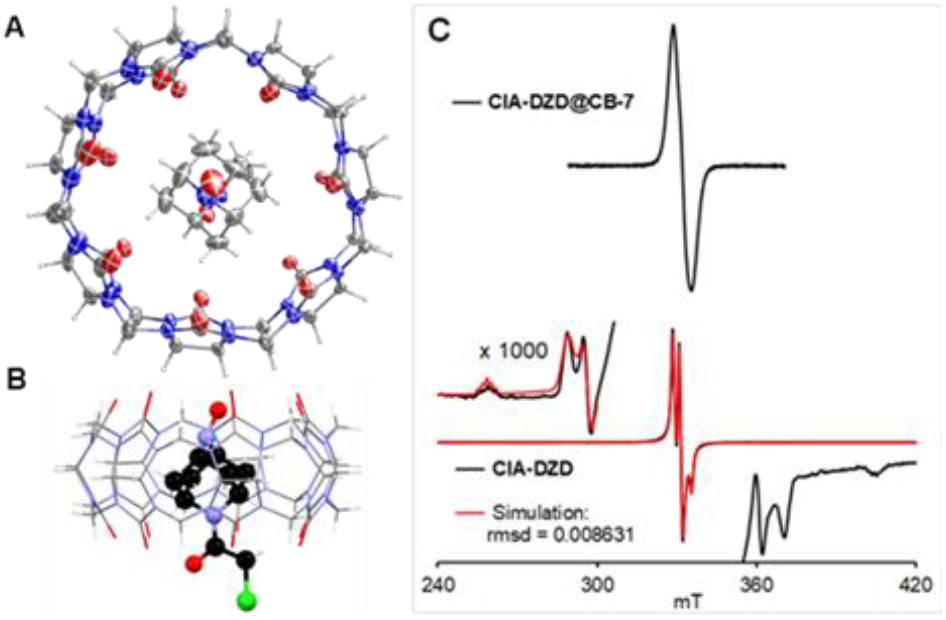
X-ray crystallography of ClA-DZD@CB-7 at 123 K: (A) top view, Ortep plot with carbon, nitrogen, oxygen, and chlorine atoms depicted with thermal ellipsoids set at the 50% probability level and (B) side view, the guest and host are shown in ball-and-stick and in wire plots. Molecules of water are omitted for clarity. (C) CW EPR spectra of polycrystalline ClA-DZD@CB-7 and ClA-DZD at room temperature. For more details, see: SI, Tables S2 and S3, and Figs. S2–S6.

We confirm the presence of nitroxide radicals in the crystals of ClA-DZD@CB-7 by EPR spectroscopy. The spectrum consists of a single peak, which is broadened by unresolved magnetic dipole-dipole interactions, corresponding to inter-spin distances of about 13 Å (Fig. S6, SI). This result agrees with the nearest neighbor nitroxide-nitroxide distances in the X-ray structure, as illustrated by the O∙O contact range of 10–21 Å within the asymmetric unit. Notably, this spectrum is significantly different compared to that for polycrystalline ClA-DZD, which consists of well resolved center peaks for an *S* = ½ crystal defects and the broader side-peaks; the side-peaks correspond to *S* = 1 thermally excited state for the dimer of nitroxides with nitroxide-nitroxide contacts of <2.6 Å.^25^

#### 1.3. CW EPR spectroscopy

We investigate complexation of ClA-DZD with CB-7 in Tris-MOPS-1 buffer (Table 1) by CW EPR spectroscopy (Figure 3).

**Figure 3.**
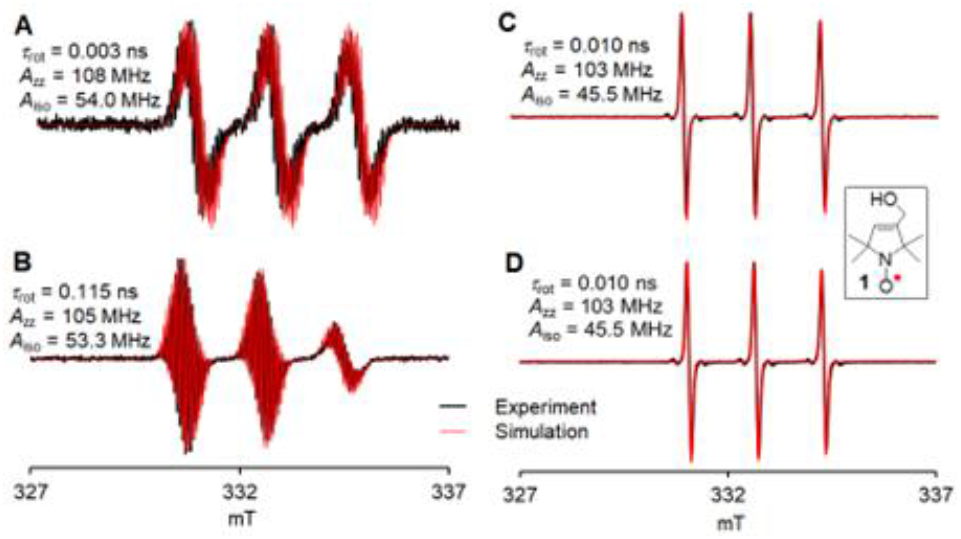
EPR spectroscopy in fluid Tris-MOPS-1 buffer (Table 1) at 295 K: rotational correlation times, *τ*_cor_, and ^14^N hyperfine tensor components, *A*_zz_, and isotropic ^14^N hyperfine couplings, *A*_iso_.^52^ (A) 0.28 mM ClA-DZD (B) 0.28 mM ClA-DZD with 2 equiv of CB-7; (C) 0.25 mM model pyrroline nitroxide **1**; (D) 0.25 mM **1** with 2 equiv of CB-7.

We first acquire the EPR spectrum of 0.28 mM ClA-DZD to determine the ^14^N hyperfine coupling, *A*^iso^, and *τ*_rot_.^52^ Upon addition of 2 equiv of CB-7, the spectrum undergoes a dramatic change, with a small decrease of *A*_iso_, from 54 to 53.3 MHz, and a large increase of *τ*_rot_, by a factor of 36, from 0.003 to 0.115 ns. For comparison, we carry out the analogous experiment using pyrroline nitroxide **1** (a model compound for MTSL), from which we find that *A*_iso_ = 45.5 MHz and *τ*_rot_ = 10 ps remain unchanged by addition of CB-7, indicating the absence of complexation by CB-7. The decrease in *A*_iso_ is expected upon inclusion of nitroxide radical into the hydrophobic cavity of CB-7.^35,38,42,53,54^ However, the formation of the ternary complex (see above, ITC) presumably impedes the decrease of *A*_iso_.^35,38,39,42,50,53,54^ Unlike all nitroxides studied for CB-7 or CB-8 complexation to date,^35,38,42,50,53,54^ ClA-DZD is a conformationally-fixed, relatively rigid radical, with much greater charge separation within the NO moiety as suggested by the greater magnitude of its *A*_iso_,^55^ compared to typical nitroxides, including **1** (Figure 3). The rigidity and the increased charge separation in ClA-DZD might also contribute to the cooperativity in binding of Na^+^ to nitroxide@CB-7 complex, as observed in the ITC studies. The increase in *τ*_rot_ and decrease in *A*_zz_ are analogous to what was observed when IA-DZD was complexed with β-CD.^56^

### 2. Complexation of DZD-T4L with CB-7

Spin labeling of the T4L mutants is carried out using established methods as described in the Experimental section, to obtain DZD-T4L 135, DZD-T4L 65, and doubly labeled DZD-T4L 65/80 and DZD-T4L 65/135.

#### 2.1. X-ray Crystallography of DZD-T4L 65

The crystals of *DZD-T4L 65* were grown using established conditions as described in the Experimental section. The DZD label is attached through a chemically robust sulfide (S-C) linkage to the sole cysteine residue of the triple C54T/C97A/K65C T4L mutant, placing the label near the N-terminal portion of helix C, which connects the two domains of the enzyme (Figure 4A). Notably, DZD-T4L is structurally nearly identical to the unmodified protein, with a C_α_ RMSD of 0.34 Å to PDB 2LZM.^57^

**Figure 4.**
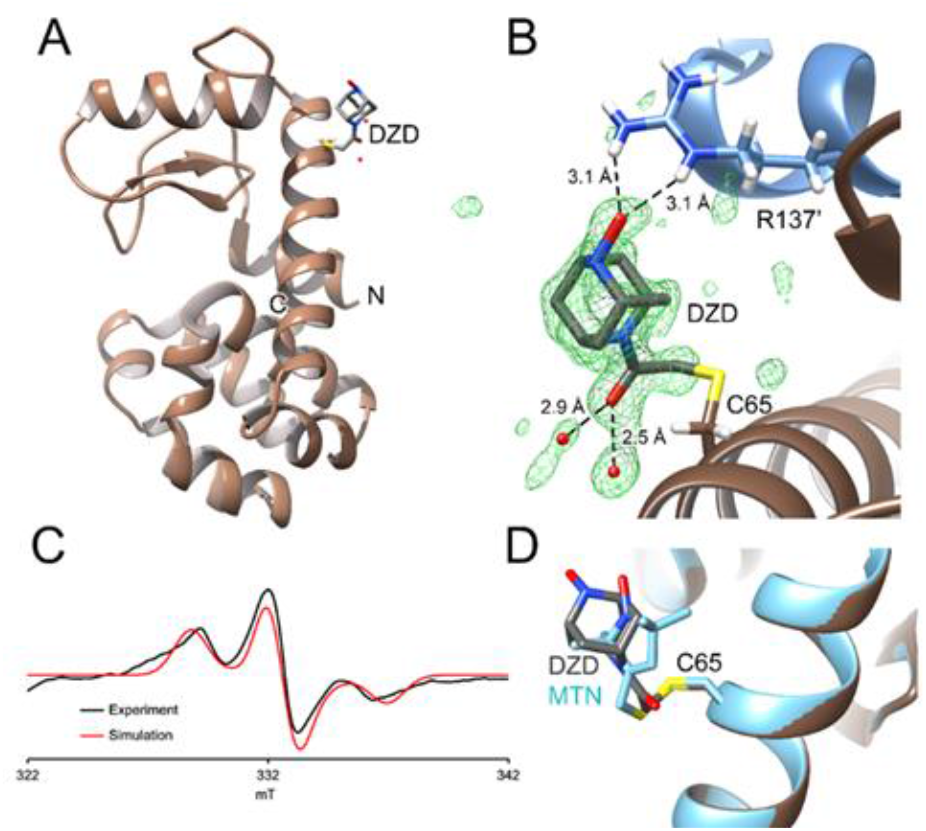
DZD-T4L 65: (A) The ribbon diagram of T4L showing the DZD label at C65 near the N-terminal region of helix C. (B) Minimally biased omit mF_o_-DF_c_ electron density contoured at 3.0 σ (green) for the DZD label, which was calculated prior to the introduction of DZD into the structural model. Hydrogen bonds are shown as dotted lines with distances. R137 is from a neighboring molecule in the crystal lattice (blue, primed label) and makes a bifurcated hydrogen bond with the oxygen atom of the nitroxide label. (C) The CW EPR spectrum of polycrystalline DZD-T4L in reservoir solution prior to X-ray data collection confirms the presence of the radical on the spin label. (D) A comparison of the DZD label (this work, grey) and the MTN spin label from PDB 6PGY, showing similar overall orientations in the crystal.

X-ray diffraction data were recorded to 1.12 Å resolution, allowing the DZD label to be modeled into unambiguous mF_o_-DF_c_ difference electron density extending from Cys65 (Figure 4B). The label occupancy is 0.88 as refined in PHENIX and its ADPs are elevated (DZD, <ADP>=25.38 Å^2^), compared to neighboring atoms (Cys65 <ADP>=13.13 Å^2^), indicating some disorder of the label. Additional evidence for local disorder is the presence of some minor unmodeled difference density in the vicinity of the DZD moiety. Apart from the covalent attachment to Cys65, the label makes only one other direct contact with the protein: a bifurcated hydrogen bond^58^ between the O1 atom of the label to the guanidinium group of Arg137 from a neighboring molecule in the lattice (Figure 4B). However, there are two hydrogen bonds between the carbonyl oxygen atom (O2) of the DZD to nearby water molecules that provide indirect, solvent-mediated contacts between the label and the protein (Figure 4B). Because these contacts are predominantly with neighboring molecules in the lattice, it is likely that the DZD label has more orientational freedom when T4 lysozyme is in solution.

We attempted to both co-crystallize and soak CB-7 into DZD-T4L crystals without success. The crystal structure of DZD-T4L indicates that the CB-7 is too large to be accommodated in the crystal lattice at the Cys65 label site. These constraints are specific to this crystal lattice and are not expected to play a role in the solution behavior of CB-7-DZD-T4L complexes.

We determined the chemical species present at the N-O bond of the DZD label by unrestrained refinement and bond length analysis (see: Experimental). The N1-O1 bond length is 1.5806(611) Å after removing the distance restraint on this bond and inverting the least squares matrix in SHELXL. The PHENIX-refined model, which is the one deposited in the PDB, has a restrained N-O bond length of 1.54 Å. In both cases, this bond length is markedly longer than the expected N-O bond length for a nitroxide radical, which is 1.2843(19) Å in IA-DZD spin label.^25^ Therefore, we conclude that the crystallized DZD label on T4L is likely the corresponding hydroxylamine. The DZD-T4L crystals initially contain some radical with the expected EPR features of DZD, indicating that the radical species survives crystallization (Figure 4C). One possible reason for conversion of the radical to the hydroxylamine in the crystal structure is X-ray radiation damage from the intense 17 keV synchrotron X-ray beam, although we did not test this.

T4L is an established protein system for characterizing spin labels and multiple crystal structures have been determined of different spin labeled forms of the enzyme. ^59-64^ Because the motivation for deploying spin labels in T4L is often to characterize the site-resolved conformational dynamics of the protein, the sites of attachment differ in these studies. Hubbell and coworkers used the same K65C mutation in helix C to attach an MTSL nitroxide spin label to T4L.^59^ Two crystal structures containing closely related *S*-(1-oxyl-2,2,5,5-tetramethyl-2,5-dihydro-1H-imidazol-4-yl) methanesulfonothioate (PDB 3K2R) and *S*-[(1-oxyl-2,2,5,5-tetramethyl-2,5-dihydro-1H-pyrrol-3-yl)methyl] methanesulfonothioate (PDB 6PGY) attached to Cys65 via a disulfide are available, although neither have associated publications yet. Overall, the orientation of these spin labels is similar to DZD (Figure 4D), although neither makes the hydrogen bond between the nitroxide oxygen atom to the symmetry-related Arg137 residue that is observed in the present structure. Because of the absence of a stabilizing contact, the spin label-T4L tends to have more crystallographically disordered labels, with correspondingly weaker supporting electron density.

#### 2.1. CW EPR spectroscopy

Analysis of the room temperature CW spectra of DZD-T4L 135, DZD-T4L 65, and doubly labeled DZD-T4L 65/80 in 2:1 buffer/glycerol (w/g) matrix, show that binding of the label to the protein increases *τ*_rot_ to 1.33, 1.41, and 1.76, respectively (Table S5, Figs. S14-S16, SI). The position dependence of *τ*_rot_ is attributed to different interactions with neighboring sidechains. Upon addition of 2 – 4 equiv of solid CB-7 changes in EPR spectra indicate increases in *τ*_rot_ to 1.62, 1,62, and 2.00 ns, respectively and smaller values of *A*_zz_ and *A*_iso_ (Table S5, Figs. S14–S17, SI). For DZD-T4L 65/80, *A*_iso_ = 55.7 MHz decreases to 53.3 MHz upon addition of CB-7 (Fig. S18, SI). The increases in *τ*_rot_ suggest greater steric interactions with neighboring protein sidechains for the bulkier CB-7 bound spin label than for the label alone. The significant decreases in *A*_iso_ suggest a more hydrophobic environment for nitroxide radical inside the CB-7 cavity, similar to what was observed for ClA-DZD binding to CB-7 (Figure 3, Table S5) or IA-DZD to β-CD.^25^ In summary, these results reveal the strong host-guest molecular recognition of nitroxide in DZD-T4L by CB-7 in 2:1 w/g matrix.

#### 2.3 Electron spin relaxation times

Binding to CB-7 causes a significant increase of the electron spin coherence time, *T*_m_, at 80 K and 150 K; e.g., for DZD-T4L/CB-7 at 80 K, *T*_m_ = 4.9 – 5.0 μs, compared to *T*_m_ = 4.4 – 4.5 μs for DZD-T4L/β-CD, *T*_m_ = 4.2 μs for DZD-T4L or IA-DZD, and *T*_m_ = 2.1 – 2.7 μs for MTSL (Table 2).

**Table 2.** Electron spin relaxation times: *T*_m_ and *T*_1_.^a^.

| Protein mutant | spin label | $T$ (K) | $T_m$ (μs) | $\beta$ | $T_1$ (μs) | $\beta$ |
| --- | --- | --- | --- | --- | --- | --- |
| / | MTSL | 80 | 2.06 | 0.96 | - | - |
|  |  | 140 | 0.31 | 0.71 | - | - |
|  | IA-DZD | 80 | 4.22 | 2.24 | 640 | - |
|  |  | 150 | 3.38 | 1.82 | 108 | - |
|  | IA-DZD/CB-7 | 80 | 4.67 | 2.34 | 455 | - |
|  |  | 150 | 3.72 | 1.70 | 100 | - |
| T4L 65/80 | DZD | 80 | 4.16 | 2.12 | 261 | 0.72 |
|  |  | 80 | 4.46 | 2.14 | 320 | 0.71 |
|  |  | 150 | 3.20 | 1.59 | 64 | 0.74 |
|  |  | 150 | 3.83 | 1.58 | 70 | 0.73 |
|  | DZD/CB-7 | 80 | 4.97 | 2.33 | 481 | 0.74 |
|  |  | 150 | 4.62 | 2.08 | 124 | 0.77 |
| | DZD/β-CD | 80 | 4.45 | 2.10 | 208 | 0.68 |
|  |  | 150 | 3.82 | 1.77 | 50 | 0.70 |
| T4L 65/135 | DZD | 80 | 4.20 | 2.03 | 218 | 0.70 |
|  |  | 80 | 4.20 | 2.01 | 224 | 0.69 |
|  |  | 130 | 3.63 | 1.63 | 85 | 0.72 |
|  |  | 150 | 3.24 | 1.50 | 76 | 0.70 |
|  | DZD/CB-7 | 80 | 4.94 | 2.38 | 510 | 0.70 |
|  |  | 150 | 4.27 | 1.89 | 365 | 0.73 |
| | DZD/β-CD | 80 | 4.40 | 2.03 | 229 | 0.70 |
|  |  | 150 | 3.60 | 1.64 | 54 | 0.72 |

<sup>a</sup> $\beta$ , exponential stretch parameter; data for β-CD<sup>25</sup> were re-fit with stretch exponential. Relaxation times are measured at the maximum amplitude peak in the echo-detected field-sweep spectra (Figs. S19 and S21, SI). T4L solutions contained 1 mM CB-7. MTSL data are obtained in trehalose/sucrose; in glycerol/water at 76 K, $T_m = 2.7$ μs is obtained. Uncertainties are >5% for $T_m$ and about 10% for $T_1$ .

The exponential stretch parameter, *β*, increases upon complexation with CB-7, indicating enhanced immobilization of nitroxide. We note that *T*_1_ at 80 K, increases from 200 – 300 μs for DZD-T4L to 500 μs for DZD-T4L/CB-7. The increase of *T*_1_ limits the shot repetition time, >1.5\**T*_1_, for DEER experiments, resulting in slower data acquisition.

### 3. DEER Distance Measurements

The DEER distance measurements are carried out on the doubly labeled DZD-T4L and MTSL-T4L mutants 65/80 and 65/135 in a 2:1 w/g matrix without and with 1 mM CB-7 at 83 K. These two T4L mutants contain cysteines at solvent accessible sites that span C_β_–C_β_ distances of 2.2 and 3.6 nm. Upon addition of CB-7 to the DZD-T4L mutants, significantly narrower main components of inter-spin distance distribution are observed, while the MTSL-T4L mutants are not affected by addition of CB-7 (Figure 5, Table 3).

**Table 3.** DEER distance distributions and sensitivity as limiting concentration of spin label [LC].^a^.

| $d_2^b$<br>(μs) | T4L | spin<br>label | w/g<br>matrix | $T$<br>(K) | $r$ ($\sigma$)<br>[nm] | [LC]<br>(μM) |
| --- | --- | --- | --- | --- | --- | --- |
| 3.5 | 65/80 | DZD |  | 83 | 2.44 (0.29) | 6.48 |
|  |  |  | + CB-7 |  | 2.56 (0.19) | 3.28 |
|  |  | MTSL |  |  | 2.60 (0.34) | 14.5 |
|  |  |  | + CB-7 |  | 2.60 (0.35) | 12.9 |
|  | 65/135 | DZD |  |  | 4.56 (0.47) | 3.86 |
|  |  |  | + CB-7 |  | 4.90 (0.19) | 1.72 |
|  |  | MTSL |  |  | 4.45 (0.27) | 13.0 |
|  |  |  | + CB-7 |  | 4.51 (0.26) | 16.2 |
| 2.0 | 65/80 | DZD | + CB-7 | 83 | 2.62 (0.18) | 0.57 |
|  |  |  |  | 150 | 2.63 (0.26) | 1.15 |
|  |  |  |  | 170 | 2.58 (0.23) | 1.78 |
|  |  |  |  | 180 | 2.68 (0.69) | 3.40 |
|  |  |  |  | 190 | 2.61 (0.23) | 12.2 |
| 1.2 | 65/80 | DZD | + CB-7 | 200 | 2.73 (0.49) | 38.1 |

<sup>a</sup> [LC] = $\{10 \cdot [\text{SL}] \cdot (\# \text{scans})^{1/2}\} / \text{SNR}$ , where [SL] is concentration of spin label and SNR is signal-to-noise ratio. <sup>b</sup> $d_2$ = echo acquisition time. More details, including data for [LC] and β-CD,<sup>25</sup> may be found in the SI: Tables S8 and S9.

**Figure 5.**
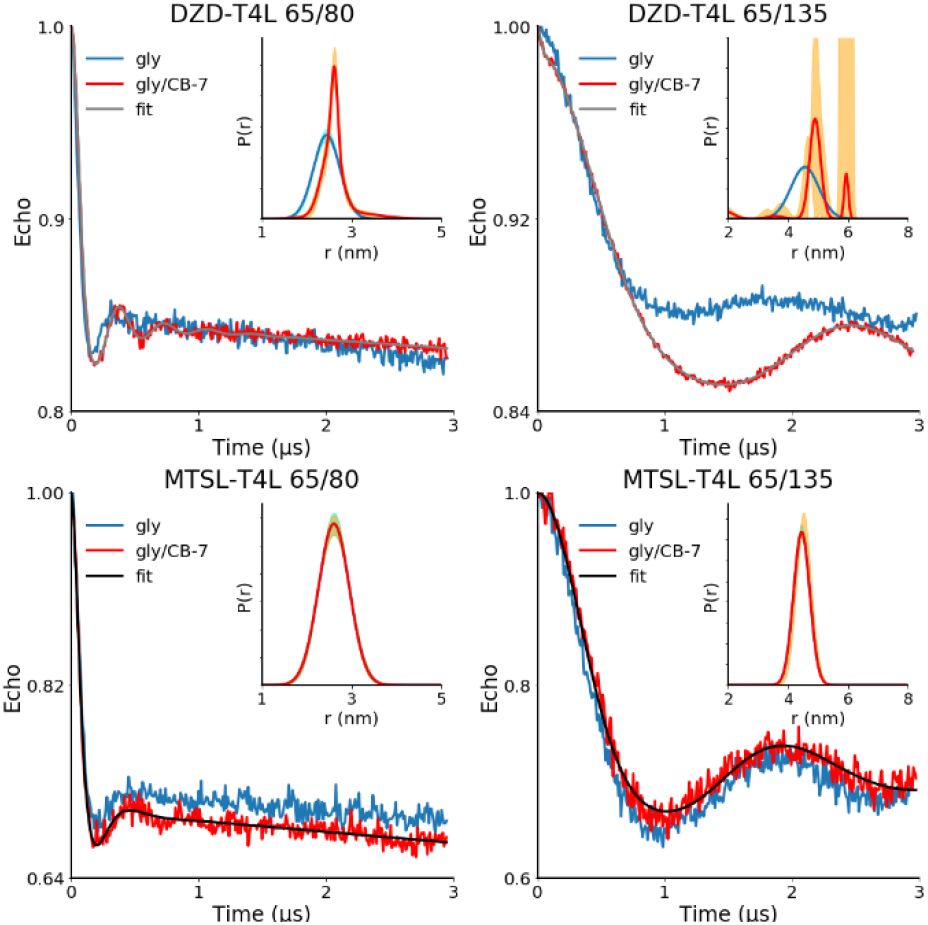
DEER distance measurements on doubly spin labeled DZD-T4L and MTSL-T4L in 2:1 w/g matrix without CB-7 (gly) and with 1 mM CB-7 (gly/CB-7) at 83 K. The water phase is Tris-MOPS-1 buffer (Table 1), i.e., 6 mM Tris, 9 mM MOPS, 0.01 mM EDTA, and 50 mM NaCl (pH 7.2). Main plots: normalized echo vs. time with fits, using a sum of Gaussians as the distance distribution. Inset plots: distance distributions with 95% confidence bands (orange lines).^65^ For additional details, including alternative distance distributions for DZD-T4L 65/135, see: SI, Tables S7 and S8, and Fig. S29.

As indicated by the limiting concentration (LC), defined as minimum concentration of spin label in the sample providing signal-to-noise ratio of 10 in one scan, sensitivity of the DEER measurements at 83 K in the presence of CB-7 increases by a factor 4 – 9 for DZD-T4L, compared to the MTSL-T4L. In addition, values of LC suggest that sensitivity of the measurements at *T* = 83 K increases by a factor of 2.0 – 2.2 upon addition of CB-7 to the DZD-T4L mutants, while it remains approximately unchanged for the MTSL-T4L mutants (Table 3). This two-fold increase in sensitivity is attributed to the longer *T*_m_ observed in DZD-T4L mutants after addition of CB-7 (Table 2, Fig. S25-S26).

We explore further the highest possible temperature for practical DEER distance measurement (Figure 6). We use doubly spin labeled DZD-T4L 65/80 in a 2:1 w/g matrix with 1 mM CB-7, with aqueous phase corresponding to the 6 mM Tris buffer.

**Figure 6.**
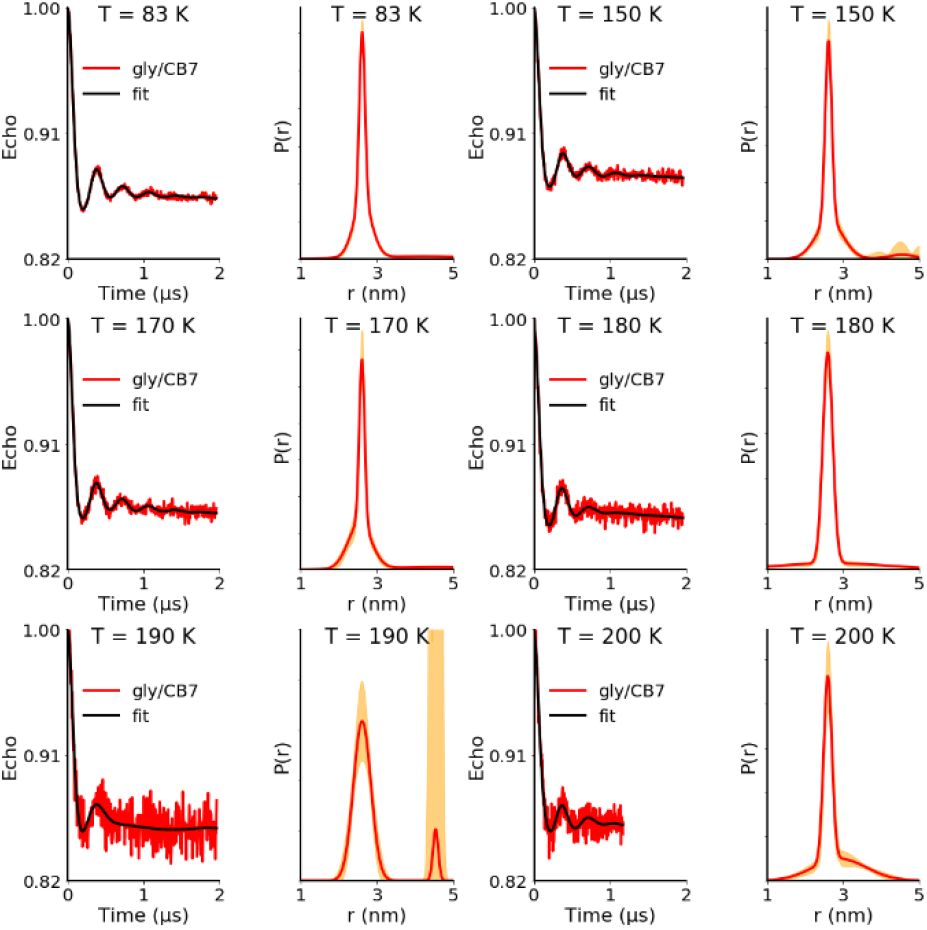
DEER distance measurements on doubly spin labeled DZD-T4L 65/80 in 2:1 w/g matrix with 1 mM CB-7 at *T* = 83– 200 K: normalized echo vs. time with fits, using a sum of Gaussians, and distance distributions, *P*(*r*), with 95% confidence bands (orange lines).^65^ The water phase is Tris-MOPS-1 buffer (Table 1). Additional information may be found in Table 3 and the SI: Table S9 and Figs. S30 and S31.

The 2:1 water/glycerol matrix (v/v), corresponding to approximately 10 mol% glycerol in water, has been postulated to consist of mesoscopic domains of glycerol-rich (40 mol%) glass/liquid and water-rich ice cores in the temperature range 80–270 K. The glycerol rich domains possess a glass transition temperature, *T*_g_ ≈ 170 K, while the ice cores undergo a melting process in a broad range of temperatures, ca. 200 – 270 K.^66,67^

Notably, we can observe DEER signal up to 200 K (Figure 6), which corresponds to the apparent onset of melting of water-rich ice cores in the 2:1 w/g matrix. To obtain a reasonable signal-to-noise ratio (SNR) in a reasonable amount of time at 200 K, we use a relatively short echo acquisition time, *d*_2_ = 1.2 μs and a short repetition time of 150 μs, to reflect a shortened *T*_1_ at this temperature. No discernible signal was seen at higher temperatures. We note that such measurements at *T* = 140 – 200 K would not be feasible on MTSL-T4L proteins, because of very small *T*_m_ = 0.1 – 0.5 μs, as determined for MTSL (Fig. S20) or simple pyrroline nitroxide radical, such as 3-carbamoyl-2,2,5,5-tetramethyl-3-pyrrolin-1-oxyl.^68^ We also note that MTSL-T4L 96 in 2:1 w/g matrix showed a discontinuity in the slope of the microwave power (*P*) saturation plots, ln *P*_1/2_ vs 1/*T*, at *T* ≈ 200 K;^68^ this would suggest a significant change in electron spin relaxation times at that temperature. Further studies to understand changes in the microenvironment of nitroxide radicals, corresponding to the abrupt disappearance of DEER signal at 210 K will be needed.

## CONCLUSION

The complex of neutral 2,6-diazaadamantane-based ClA-DZD nitroxide radical with CB-7 possesses the largest association constant to date among all studied nitroxides. The radical inside the host is significantly immobilized, as indicated by the increase of rotational correlation time by a factor of 36 upon complexation. The X-ray structure of the ClA-DZD@CB-7 complex shows the guest nitroxide radical encapsulated inside of the CB-7 host, with the ClA-side chain protruding outside of the host. This structure suggests that strong complexation of DZD nitroxide attached to protein by CB-7 is feasible. A strong host-guest molecular recognition, between CB-7 and DZD-based nitroxide radical, leads to substantial increase of the electron spin coherence time, *T*_m_, for DZD-T4L/CB-7 which is 4.9 – 5.0 μs at 80 K and 4.3 – 4.6 μs at 150 K. These relatively long *T*_m_ improve sensitivity of the DEER measurements at 83 K by a factor 4 – 9, compared to a common spin label such as MTSL, which is not affected by CB-7. In addition, a 3-nm distance may be reliably measured in aqueous buffer/glycerol up to a temperature of 200 K, which is near the glass transition/melting temperature of the matrix, thus bringing us closer to the goal of supramolecular recognition-enabled long-distance DEER measurements at near physiological temperatures. The X-ray structure of DZD-T4L 65 was obtained at 1.12 Å resolution, thus allowing for unambiguous modelling of the DZD label (0.88 occupancy). Most importantly, the presence of DZD label does not affect the structure and conformation of the protein.

## EXPERIMENTAL SECTION

### X-ray Crystallography

#### ClA-DZD@CB-7

Crystals of ClA-DZD@CB-7 were obtained by slow evaporation of solvent form 1:1 molar solution of ClA-DZD and CB-7 in water. Data were collected at 123 K using *Mo Kα* radiation and integrated (SAINT).^69^ Intensity data were corrected for absorption using the Multi-Scan method (TWINABS).^70^ Space group *P*-1 was determined based on intensity statistics and the lack of systematic absences. The structure was solved with intrinsic methods and refined on F^2^ using SHELX suite of programs.^71,72^ Crystal data for ClA-DZD@CB-7: C_42_H_42_N_28_O_14_, C_10_H_14_N_28_ClN_2_O_2_, 10.25 H_2_O, *M*_r_ = 1577.38 g mol^−1^, colorless block, 0.355 × 0.187 × 0.096 mm^3^, triclinic, *P*-1, *a* = 13.1522(12) Å, *b* = 24.929(2) Å, *c* = 40.858(4) Å, *V* = 13282(2) Å^3^, *Z* = 8, *μ*(*Mo Kα*) = 0.167 mm^−1^, *θ*_max_ = 25.24°, 292,836 reflections measured, 54,462 independent (*R*_int_ = 0.0967), *R*1 = 0.1239 [*I* > 2*σ*(*I*)], *wR*2 = 0.3413 (all data), residual density peaks: 1.420 and –1.062 e Å^−3^. Additional crystal and structure refinement data for ClA-DZD@CB-7 are in the Supporting Information and the file in CIF format, deposited to CCDC.

#### DZD-T4L 65

DZD-T4L 65 at 6.9 mg/mL in storage buffer (9 mM MOPS, 6 mM Tris pH 7.2, 0.1 mM EDTA, 50 mM NaCl, 0.02% sodium azide) was crystallized using sitting drop vapor equilibration by mixing 1 μl of protein and 1 μl of the reservoir (1.6 M sodium/potassium phosphate pH 7.4, 100 mM NaCl, 40 mM β-mercaptoethanol) and incubating at room temperature for 3-10 days. This condition is commonly used for crystallization of T4 lysozyme.^57^ Crystals measuring approximately 200×200×100 μm^3^ in space group *P*3_2_21 were cryoprotected by transfer through light mineral oil (Fisher Scientific) until all surface mother liquor was removed and then cryocooled by immersion in liquid nitrogen.

X-ray diffraction data were collected at the Stanford Synchrotron Radiation Laboratory beamline 12-2 using 17 keV radiation, a Pilatus 6M pixel-array detector (PAD), and shutterless collection with 0.2 sec exposure per 0.15° oscillation at 85% beam attenuation. Data were processed with XDS,^73^ Pointless,^74^ and Aimless^75^ with final statistics provided in Table S1, SI. The initial model was obtained by molecular replacement in PHASER,^76^ using PDB 7LXA as a search molecule,^77^ and then refined in PHENIX^78^ using riding hydrogen atoms and anisotropic atomic displacement parameters (ADPs). Relative weights for both geometry and ADPs were optimized. Initial inspection of mF_o_-DF_c_ electron density allowed unambiguous placement of the DZD label, which was subsequently refined using restraints generated in PRODRG.^79^ The model was manually improved in COOT^80^ between cycles of refinement and validated using tools in COOT,^80^ PHENIX,^78^ and MolProbity.^81^ Potassium and chloride ions were modeled in electron density where anomalous difference map peaks (>4σ) and the local environment indicated probable ion identity. Anisotropic ADPs were validated using the PARVATI server.^82^ Crystallographic data and model statistics for DZD-T4L 65 are presented in Table S1. The unrestrained length and estimated standard uncertainty (ESU) of the N-O bond of the DZD label was determined using SHELXL (version 2018/3),^72^ as distributed in the CCP4 suite.^83^ The converged PHENIX model was refined in SHELXL using a conjugate gradient least squares target function based on intensities (keyword HKLF 4). Riding hydrogen atoms and anisotropic ADPs were refined for the whole model and the SHELX version of the PRODRG restraints were used for the DZD label. Upon reaching convergence, the distance restraint (DFIX) on the N1-O1 bond was removed and the model was refined for an additional 20 cycles. The full least squares matrix on coordinates alone (i.e., the Hessian matrix excluding ADP elements) was inverted and ESUs determined by a final cycle of unrestrained refinement with full least squares (keywords L.S. and BLOC1) in SHELXL. Several crystals were transferred in the reservoir buffer into 3 quartz capillaries, sealed with capillary wax (Hampton Research cat # HR4-328) and used for EPR spectroscopy.

#### Complexation of ClA-DZD with CB7

##### Isothermal titration calorimetry (ITC)

association constants, *K*_a_, and thermodynamic parameters for the formation of inclusion complexes of ClA-DZD were obtained from ITC (VP-ITC Microcal Inc., Northampton, MA). The experiments were carried out at 25 °C in a buffer or purified water as specified in Table 1. In a typical procedure, a degassed aliquot (1.42 mL) of 0.2 mM CB-7 solution was added to the reaction cell and an identical volume of solvent (buffer or water) without CB-7 was placed in the reference cell. A degassed 2.1 mM ClA-DZD solution (300 μL) was loaded in the titration syringe, and then added in 10 μL aliquots to the reaction cell under continuous stirring at 260 rpm. The titrant was injected over 12 s with an interval of 300 s between the injections. Typical ITC thermograms are provided in the SI, Figs. S7–S13.

##### EPR spectroscopy

CW EPR spectra were obtained on a Bruker EMX+ equipped with Bruker high-sensitivity cavity (Figures 2–4). (Field-swept echo detected spectra were obtained as described below.) The CW spectra were simulated with EasySpin, using either *pepper* or *chili* modules.^52^ More details may be found in the SI.

### Spin labeling of T4L

Sample labeling was carried out as previously described.^25,84,85^ Briefly, T4L mutants were expressed in K38 cells in Luria Broth (LB) and purified using cation exchange chromatography. Labeling of the mutants was initiated by adding an excess of spin-label and incubating for 2 hours at room temperature, a second addition of spin-label was made and the samples incubated overnight at 4 °C. The samples were then desalted and concentrated.

### Electron spin relaxation studies

Field-swept echo detected spectra and relaxation times were recorded at X-band and Q-band on a Bruker E580 equipped with an Oxford ESR935 cryostat and a Bruker X-band ER4118X-MS5 split ring resonator or Q-band ER5107D2 resonator. The length of a π/2 pulse was 20 ns at X-band and 40 ns at Q-band. Field-swept echo-detected spectra were obtained with a two-pulse π/2-*τ*-π-*τ*-echo sequence and 2-pulse phase cycling. The constant *τ* was 180 ns at X-band and 200 ns at Q-band. *T*_m_ was measured by two-pulse echo-decay with a π/2-*τ*-π-*τ*-echo sequence and two-step phase cycling. *T*_1_ was measured by inversion recovery with a π-T-π/2-*τ*-π-*τ*-echo sequence with variable T that is stepped to define the relaxation process. and two-step phase cycling. The constant *τ* in the inversion recovery pulse sequence was 200 ns at X-band and 360 ns at Q-band. Echo decays and inversion recovery curves were fitted with stretched exponentials (Table S6, Figs. S23, S24, S28).

### DEER measurements

The dipolar time evolution data was obtained at 83–200 K using a standard DEER four-pulse protocol, (π/2)mw1–τ1– (π)mw1–τ1–(π)mw2–τ2–(π)mw1–τ2–echo,^86^ on a Bruker 580 pulsed EPR spectrometer operating at Q-band frequency (33.9 GHz). All pulses were square pulses with lengths of 12, 24, and 40 ns, for (π/2)mw1, (π)mw1, and (π)mw2, respectively. Temperature control was provided by an Oxford Instruments continuous flow cryostat controlled by an ITC503 temperature controller unit. Distance distributions were obtained from the time evolution data by assuming that the distance distribution can be approximated by a sum of Gaussians,^87,88^ including Gaussian distance distributions with 95% confidence bands.^65^

## Supporting information

SI_Cucurbit 7 uril enhances distance measurements of spin-labeled proteins

## ASSOCIATED CONTENT

(Word Style “TE_Supporting_Informati**on”). Supporting Information**. A brief statement in nonsentence format listing the contents of material supplied as Supporting Information should be included, ending with “This material is available free of charge via the Internet at http://pubs.acs.org.“ For instructions on what should be included in the Supporting Information as well as how to prepare this material for publication, refer to the journal’s Instructions for Authors

### Accession Codes

Structure factors and coordinates for DZD-T4 lysozyme are available from the Protein Data Bank with accession code 8F11 (PDB id).

CCDC 2289150 contains the supplementary crystallographic data for this paper. These data can be obtained free of charge via www.ccdc.cam.ac.uk/data_request/cif, or by emailing, or by contacting The Cambridge Crystallographic Data Centre, 12 Union Road, 495 Cambridge CB2 1EZ, UK; fax: +44 1223 336033..

## AUTHOR INFORMATION

### Author Contributions

The manuscript was written through contributions of all authors. All authors have given approval to the final version of the manuscript.

### Notes

Any additional relevant notes should be placed here.

## ACKNOWLEDGMENT

We thank the National Institutes of Health (R01GM124310-01 to AR, SR, SSE, and HM) and National Science Foundation (CHE-1955349 and CHE-2247170 to AR) for support of this research. M.A.W. is supported by NIH R01GM139978. Support for the acquisition of the Bruker Venture D8 diffractometer through the Major Scientific Research Equipment Fund from the President of Indiana University and the Office of the Vice President for Research is gratefully acknowledged. We thank Dr. Chan Shu (Nebraska) for obtaining EPR spectrum of DZD-T4L 65 crystals (Figure 4c) and reproducing selected ITC experiments.

SYNOPSIS TOC (Word Style “SN_Synopsis_TOC”). If you are submitting your paper to a journal that requires a synopsis graphic and/or synopsis paragraph, see the Instructions for Authors on the journal’s homepage for a description of what needs to be provided and for the size requirements of the artwork.

