## Supplementary material for "Cucurbit[7]uril Enhances Distance Measurements of Spin-Labeled Proteins": SI_Cucurbit 7 uril enhances distance measurements of spin-labeled proteins

### Table of Contents

1. General procedures and materials ------------------------------------------------------------ p. S2

2. X-ray crystallography and EPR spectrum of DZD-T4L-65

   (Table S1 and Fig. S1) ------------------------------------------------------------------------ pp. S3–S4

3. X-ray crystallography and

EPR spectrum of DZD@CB-7

   (Tables S2 and S3, Figs. S2 – S6) ------------------------------------------------------------ pp. S5–S12

4. Complexation of ClA-DZD with CB-7: ITC data (Table S4, Figs. S7–S13) ------------- pp. S12–S19

5. Complexation of ClA-DZD and DZD-T4L mutants with CB-7: CW EPR -------------- pp. S19–S24

   spectroscopy at UNL, DU, and Vanderbilt (Table S5 and Figs. S14–S17).

6. Field-swept echo-detected spectra and simulations (Fig. S18) ------------------------------ p. S25

7. Electron spin relaxation studies on spin labels and doubly spin labeled ------------------ pp. S26–S35

   DZD-T4L (Table S6, Figs. S19 – S28).

8. Spin labeling and DEER measurements (Tables S7 and S8, Figs. S29–S31) ------------ pp. S36–S41

9. Addition of 1 mM CB-7 does not affect the T4L structure ----------------------------------- p. S42

10. References for Supporting Information ------------------------------------------------------ pp. S42–S44

#### 1. General procedures and materials.

Throughout the following paragraphs labels "ZY241" and alike correspond to sample or experiment codes directly traceable to the laboratory notebooks or raw data.

Per-deuterated solvents for NMR spectroscopy were obtained from Cambridge Isotope Laboratories. All other commercially available chemicals were obtained from either Sigma-Aldrich or Acros, unless indicated otherwise; e.g., cucurbit[7]uril (CB-7) was mainly obtained from Strem. Syntheses of ClA-DZD and IA-DZD were described previously.[S1] Standard techniques under inert atmosphere, using vacuum lines and Schlenk glassware, were employed.

### 2. X-ray crystallography of DZD-T4L-65: Table S1.

| **Data Statistics** | |
|---|---|
| Sample | T4 lysozyme-DZD |
| Diffraction source | SSRL 12-2 |
| Wavelength (Å) | 0.729 |
| Temperature (K) | 100 |
| Detector | Pilatus 6M |
| Space group | $P3_221$ |
| a, b, c (Å) | 60.05 60.05 95.62 |
| α, β, γ (°) | 90, 90, 120 |
| Mosaicity (°) | 0.14 |
| Resolution range (Å)[1] | 35.20-1.12 (1.14-1.12) |
| Total no. of observations | 1063890 (42687) |
| No. of unique observations | 77616 (3537) |
| Completeness (%) | 99.6 (92.7) |
| Multiplicity | 13.7 (12.1) |
| $\langle I/\sigma(I)\rangle$ | 13.0 (0.7) |
| $CC_{1/2}$[2] | 1.000 (0.246) |
| $R_{meas}$[3] | 0.083 (3.729) |
| **Model Statistics** | |
| PDB accession code | 8F11 |
| Refinement program | PHENIX 1.19.2_4158 |
| Resolution range (Å) | 35.20-1.12 (1.13-1.12) |
| Completeness (%) | 99.22 (80.00) |
| No. of reflections | 77162 (2203) |
| No. of reflections, test set | 3874 (121) |
| $R_{work}$ | 0.1357 (0.3105) |
| $R_{free}$[4] | 0.1585 (0.3307) |
| No. of protein, water, heteroatoms (including DZD) | 1483, 299, 31 |
| Average root mean square deviations; bonds (Å), angles (°) | 0.007, 0.972 |
| Average ADPs; protein, water, heteroatoms (Å$^2$) | 17.37, 32.56, 22.90 |
| Average ADP anisotropy[5]; protein, water, heteroatoms | 0.427, 0.478, 0.454 |
| MolProbity clashscore[6] | 0.98 |
| Ramachandran plot; outliers, allowed, favored (%) | 0.00, 1.85, 98.15 |

[1]Values in parenthesis are for the highest resolution shell
[2]$CC_{1/2}$ was used to determine the high-resolution cutoff.[S2]
[3]Multiplicity-corrected R value for data as defined in this ref.[S3]
[4]R value for model calculated using only test set reflections as defined in this ref.[S4]
[5]Anisotropy is defined as the ratio of the smallest to largest eigenvalue of the ADP tensor and was calculated using PARVATI.[S5]
[6]Defined as the number of van der Waals overlaps of 0.4 Å or greater per 1000 atoms.[S6]

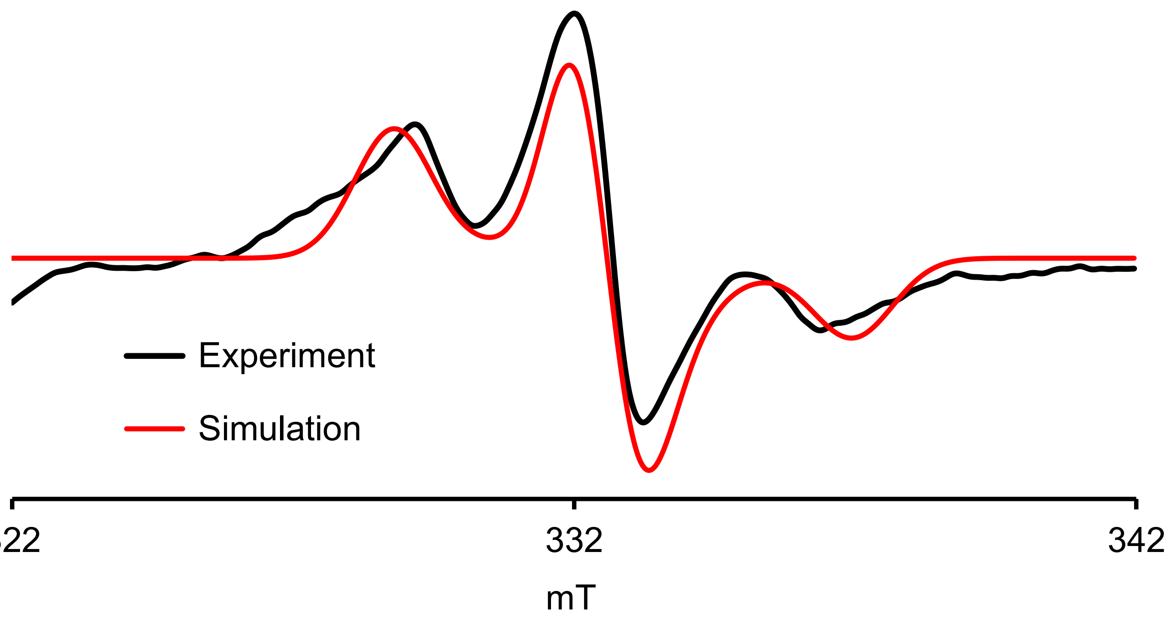

**Fig. S1.** EPR ($\nu$ = 9.3329 GHz, label: CS853R28) spectrum of polycrystalline DZD-T4L 65 suspended in buffer at 294 K; same data as in Figure 2C (main text). Simulation (*Pepper*)[S7] parameters: $S$ = 1/2, $A_{xx}$ = 21.0 MHz, $A_{yy}$ = 29.0 MHz, $A_{zz}$ = 114.3, $g_{xx}$ = 2.0076, $g_{yy}$ = 2.0050, $g_{zz}$ = 2.0029; lwpp (Gaussian) = 1.417 mT.

### 3. X-ray crystallography and EPR spectrum of ClA-DZD@CB7.

Crystallographic data were deposited in the Cambridge Crystallographic Data Centre (**CCDC 2289150**). The data can be obtained free of charge from the Cambridge Crystallographic Data Centre via www.ccdc.cam.ac.uk/data_request/cif.

The single crystals of **ClA-DZD@CB7** were obtained by slow solvent evaporation from a 1:1 molar mixture of ClA-DZD and CB-7 in water (sample label: ZY241), according to the following procedure. CB-7 was heated at 130 ºC under high vacuum line for 22 hours, and water content was determined by $^1$H NMR in DMSO-$d_6$ (Fig. S5). This treatment of CB-7 allowed for almost all of HCl to be removed as determined by pH measurements, e.g. CB-7 (0.40 mg) in nanowater (1.5 mL) gave pH 6.26 – 6.32, compared to pure nanowater pH 6.25. ClA-DZD was recrystallized from $CHCl_3$/pentane, collected and dried under high vacuum line overnight before using. Solvent was nanowater (17.6 mΩ · $cm^{-1}$), which was bubbled by $N_2$ gas flow at 1-2 bubbles/sec overnight. Vials were cleaned by ethanol, acetone and hexane, dried in oven at least overnight. 4 mM ClADZD was added dropwise into 4 mM CB7, and then the solution was gently swirled. The solution was kept in a cabinet ta room temperature in dark. After 3 – 15 d, pink prismatic or very fine crystals were obtained. It should be noted that it was important for the solution to be gently mixed and in particular, sonication may not be a good choice, as it sometimes gave cloudy CB7 stock/mixed solution, failing in providing single crystals.

A colorless crystal (approximate dimensions 0.355 × 0.187 × 0.096 $mm^3$) was placed onto the tip of a MiTeGen loop and mounted on a Bruker Venture D8 diffractometer equipped with a PhotonIII detector at 123(2) K. The data collections for crystal of **ClA-DZD@CB7** were carried out using Mo Kα radiation (graphite monochromator, $\lambda$ = 0.71073 Å) with a frame time of 2.5 and 28 seconds and a detector distance of 140 mm. A collection strategy was calculated and complete data to a resolution of 0.84 Å were collected. Twenty-four major sections of frames were collected with 0.5º ω and ϕ scans. The total exposure time was 56.86 hours. The frames were integrated [S8] considering twinning (twin law by the rows -1 0 0 0 -1 0 0.91 0 1, 180° rotation about reciprocal axis 0 0 1) with the Bruker SAINT software package[S8] using a narrow-frame algorithm. The integration of the data using a triclinic unit cell yielded a total of 292836 reflections to a maximum θ angle of 26.52° (0.80 Å resolution), of which 54462 were independent (average redundancy 5, completeness = 89.0%, $R_{int}$ = 8.84%, $R_{sig}$ = 7.53%) and 42230 (77.54%) were greater than 2σ($F^2$). The final cell constants of a = 13.1522(12) Å, b = 24.929(2) Å, c = 40.858(4) Å, α = 83.697(2)°, β = 86.761(2)°, γ = 87.192(2)°, volume = 13282.(2) Å$^3$, are based upon the refinement of the XYZ-centroids of 9018 reflections above 20 σ(I) with 5.055° < 2θ < 47.93°. Data were corrected for absorption effects using the Multi-Scan method (TWINABS).[S9] The ratio of minimum to maximum apparent transmission was 0.765. The calculated minimum and maximum transmission coefficients (based on crystal size) are 0.9430 and 0.9840. Additional crystal and refinement information may be found in Table S3.

The space group P-1 was determined based on intensity statistics and the lack of systematic absences. The structure was solved and refined using the SHELX suite of programs.[S10,S11] An intrinsic-methods solution was calculated, which provided most non-hydrogen atoms from the E-map. Full-matrix least squares / difference Fourier cycles were performed, which located the remaining non-hydrogen atoms. All non-hydrogen atoms were refined with anisotropic displacement parameters with exception of the co-crystallized $H_2O$ molecules. Hydrogen atoms were placed in ideal positions and refined as riding atoms with relative isotropic displacement parameters. Hydrogen atoms for the co-crystallized solvent $H_2O$ were not placed. The final anisotropic full-matrix least-squares refinement on $F^2$ with 3899 variables converged at R1 = 12.39%, for the observed data and wR2 = 34.13% for all data. The goodness-of-fit was 1.043. The largest peak in the final difference electron density synthesis was 1.420 $e^-/Å^3$ and the largest hole was -1.062 $e^-/Å^3$ with an RMS deviation of 0.131 $e^-/Å^3$. On the basis of the final model, the calculated density was 1.578 $g/cm^3$ and F(000), 6604 $e^-$.

Four formula units are in the asymmetric unit. The ClA-DZD guest is not disordered in one CB-7 host (Pair A) (Fig. S2). The ClA-DZD guests are disordered in two CB-7 hosts over two positions (pairs B and C) and in one CB-7 host over three positions (Pair D) (Figs. S3 and S4). Restraints and constraints were applied to model the disorder, some of which were also correlated to solvent $H_2O$ disorder. Solvent $H_2O$ positions were either refined at 100% sites, correlated (with ClA-DZD disorder) partial sites, or at fixed 50% sites. Representative bond lengths for N-O bond in nitroxides are presented in Table S2.

In alternative refinements (not the refinement presented here) Squeeze[S12] (PLATON) was used to assess void space and solvent content by electron count in the solvent accessible space. These methods provided information about the total potential solvent accessible void space per unit cell (~2200 $Å^3$ or 1/6 of the unit cell) and an electron count of ~966 per unit cell (~12 $H_2O$ per formula unit). However, a refinement with modified data, where solvent contribution to the structure factors was assessed by back Fourier transformation did not lead to a satisfactory model and residuals and difference e-density remained high.

**Table S2.** Summary of representative N-O bond lengths for ClA-DZD@CB-7 in the asymmetric unit.

| Nitroxide N-O bond | Bond length (Å) |
|---|---|
| O15A-N29A | 1.290(7) |
| O15B-N30B | 1.29(2) |
| O15C-N29C | 1.295(12) |
| O15D-N29D | 1.289(7) |
| O15E-N29E | 1.29(3) |
| O15F-N29F | 1.307(17) |
| O15G-N29G | 1.314(17) |
| average | 1.296 |

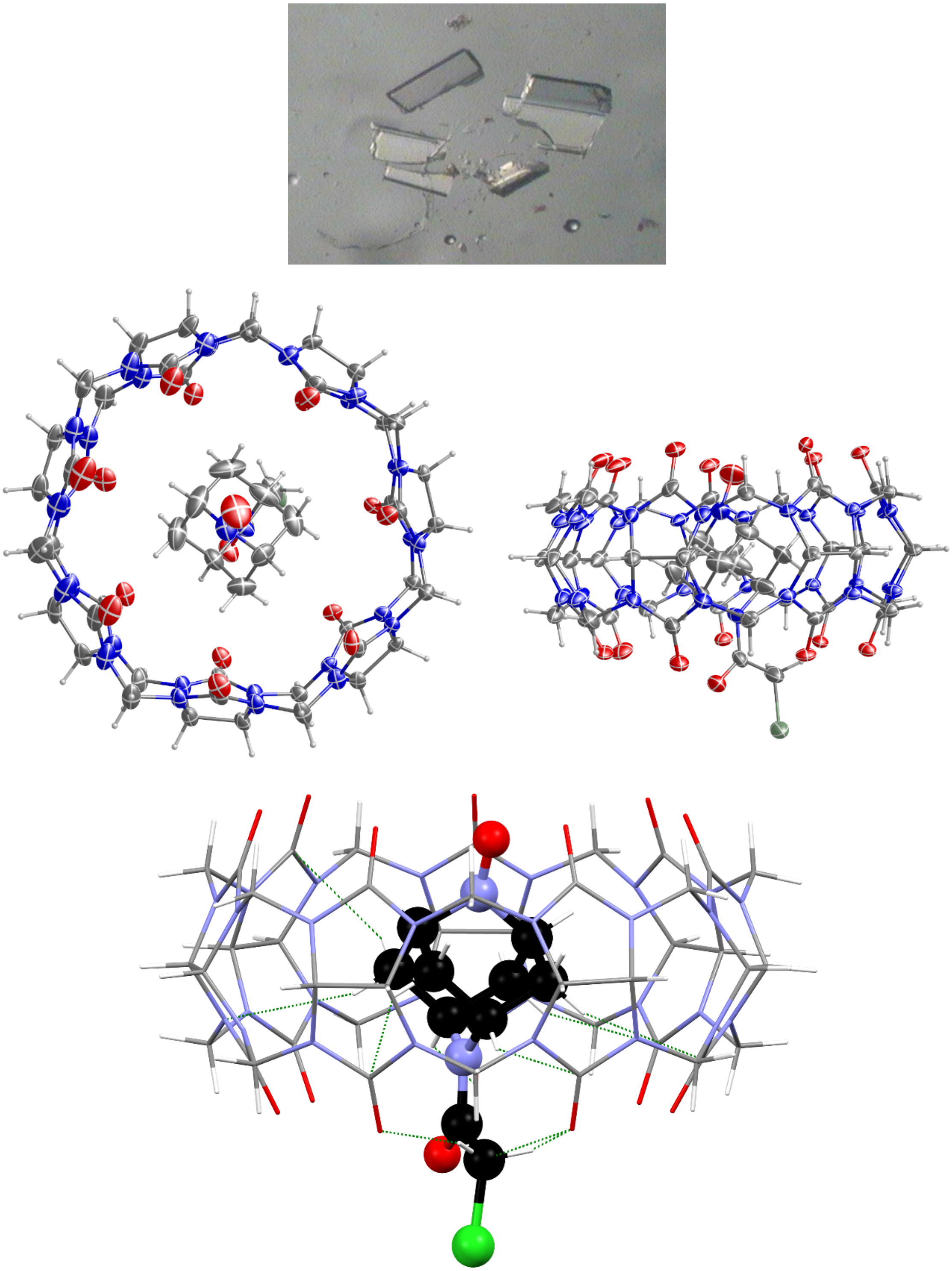

**Fig. S2.** X-ray structure of ClA-DZD@CB7 (label: 20137, sample label: ZY241). *Top panel:* photograph of bulk material. *Middle and bottom panels:* top and side view of host, guest pair A; no guest disorder is found for this pair; green dotted lines indicate 10 close contacts ≤ sum of vdW radii between the guest and the host.

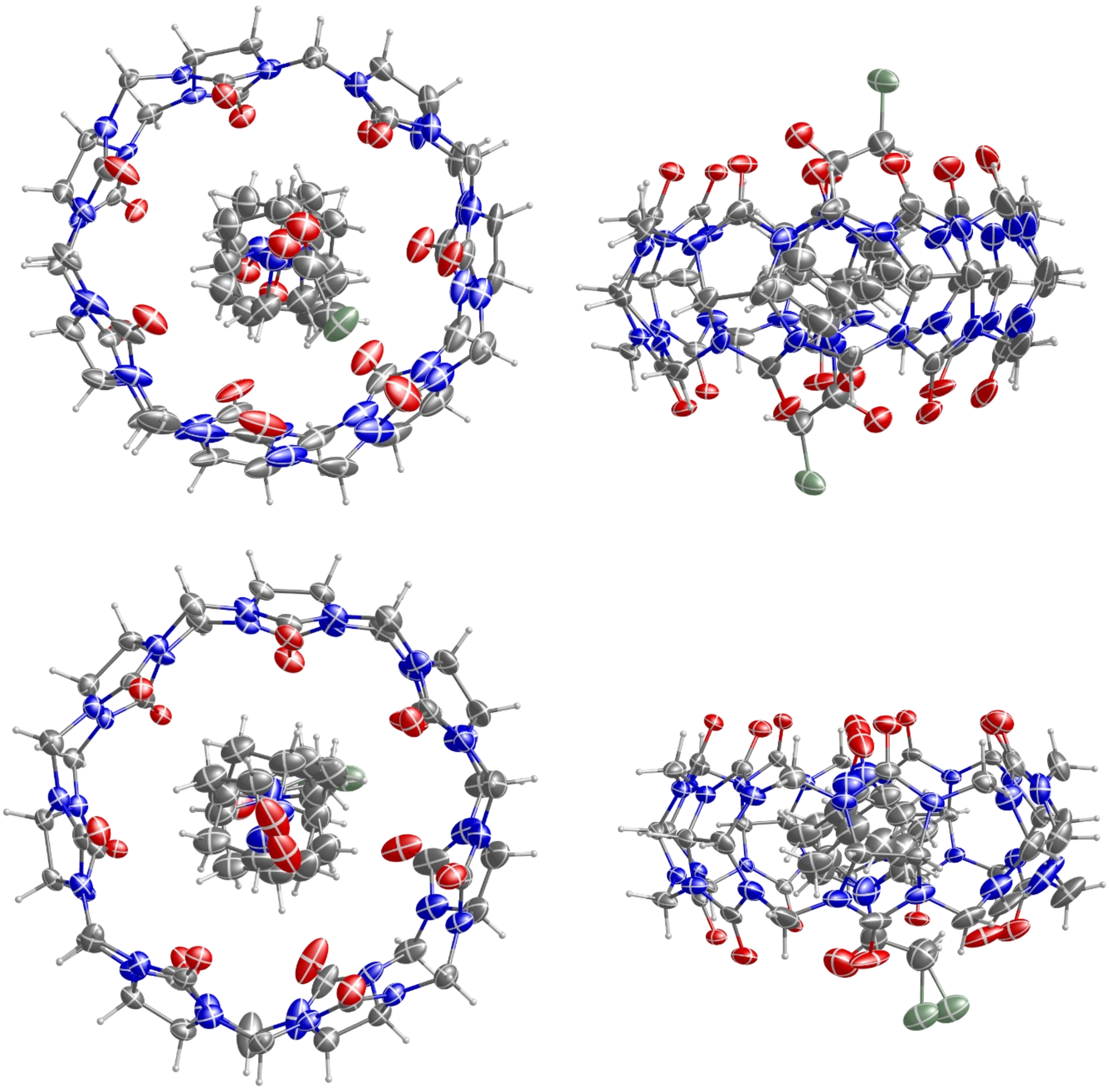

**Fig. S3.** X-ray structure of ClA-DZD@CB7 (label: 20137, sample label: ZY241): Ortep plots with carbon, nitrogen, oxygen, and chlorine atoms depicted with thermal ellipsoids set at the 50% probability level. _Top panel:_ host, guest pair B and _bottom panel:_ host, guest pair C. For each pair, top and side view of host are shown. Guest disorder over two positions is found in each pair.

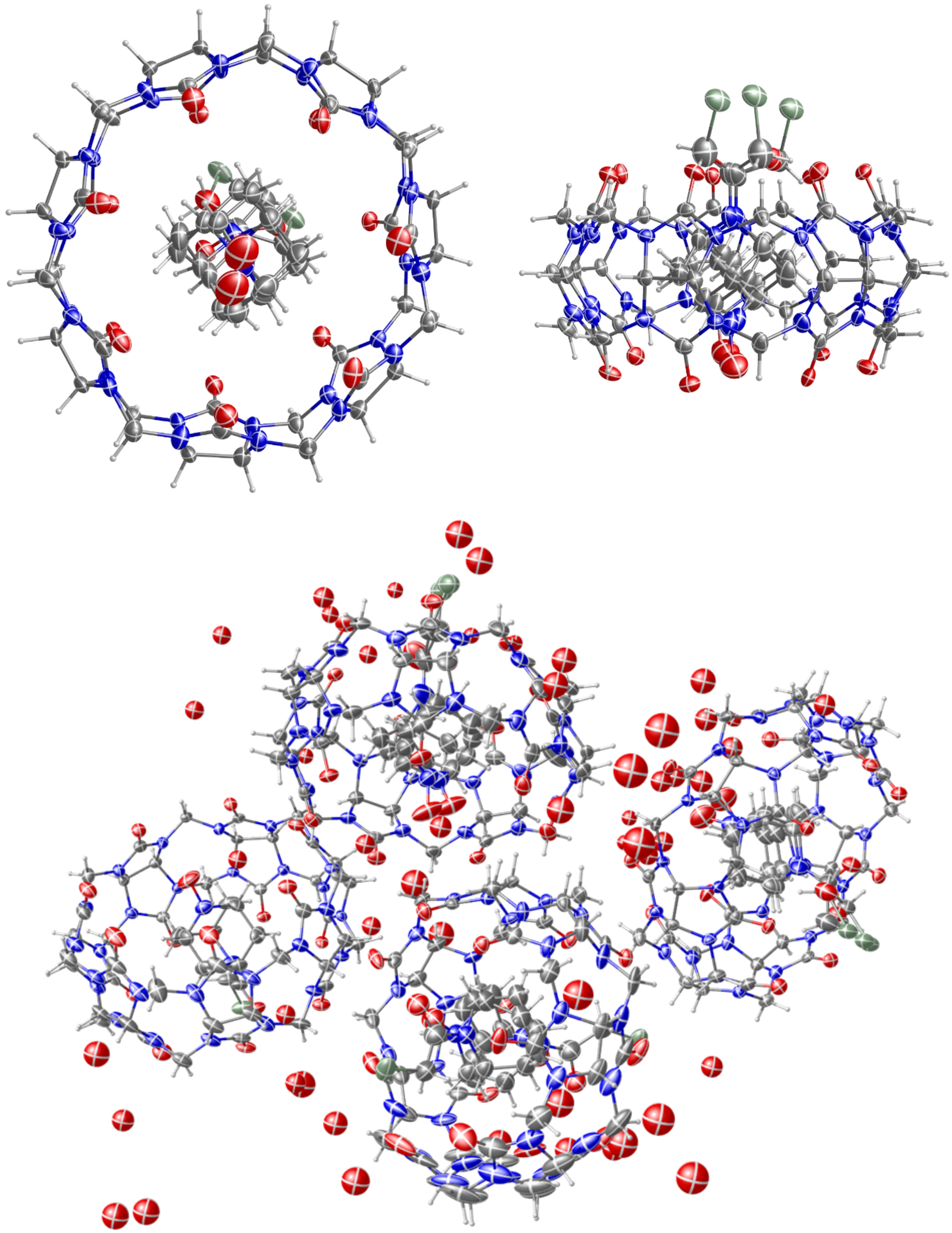

**Fig. S4.** X-ray structure of ClA-DZD@CB7 (label: 20137, sample label: ZY241): Ortep plots with carbon, nitrogen, oxygen, and chlorine atoms depicted with thermal ellipsoids set at the 50% probability level. *Top panels:* top and side view of host, guest pair D; guest disorder over three positions is found for this pair. *Bottom panel:* asymmetric unit (4 formula units of $C_{42}H_{42}N_{28}O_{14}$, $C_{10}H_{14}N_{28}ClN_2O_2$, 10.25 $H_2O$).

**Table S3**. Crystal Data and Structure Refinement for of **ClA-DZD@CB7** (label: 20137).

| | | |
|---|---|---|
| Empirical formula | C52 H76.50 Cl N30 O26.25 | |
| Formula weight | 1577.38 | |
| Crystal color, shape, size | colorless block, 0.355 × 0.187 × 0.096 $mm^3$ | |
| Temperature | 123(2) K | |
| Wavelength | 0.71073 Å | |
| Crystal system, space group | Triclinic, P-1 | |
| Unit cell dimensions | a = 13.1522(12) Å | $\alpha$ = 83.697(2)°. |
| | b = 24.929(2) Å | $\beta$ = 86.761(2)°. |
| | c = 40.858(4) Å | $\gamma$ = 87.192(2)°. |
| Volume | 13282(2) $Å^3$ | |
| Z | 8 | |
| Density (calculated) | 1.578 $Mg/m^3$ | |
| Absorption coefficient | 0.167 $mm^{-1}$ | |
| F(000) | 6604 | |
| ***Data collection*** | | |
| Diffractometer | Venture D8, Bruker | |
| Source | Iμ3.0, Incoatec | |
| Detector | Photon III | |
| Theta range for data collection | 0.502 to 26.518°. | |
| Index ranges | -16<=h<=16, -30<=k<=31, 0<=l<=51 | |
| Reflections collected | 292836 | |
| Independent reflections | 54462 [$R_{int}$ = 0.0967] | |
| Observed Reflections | 42230 | |
| Completeness to theta = 25.242° | 90.7 % | |
| ***Solution and Refinement*** | | |
| Absorption correction | Semi-empirical from equivalents | |
| Max. and min. transmission | 0.745161 and 0.575316 | |
| Solution | Intrinsic methods | |
| Refinement method | Full-matrix least-squares on $F^2$ | |
| Weighting scheme | w = [$\sigma^2Fo^2$+ $AP^2$+ BP]$^{-1}$, with | |
| | P = ($Fo^2$+ 2 $Fc^2$)/3, A = 0.1575, B = 93.5562 | |
| Data / restraints / parameters | 54462 / 6672 / 3899 | |
| Goodness-of-fit on $F^2$ | 1.043 | |
| Final R indices [I>2σ(I)] | R1 = 0.1239, wR2 = 0.3207 | |
| R indices (all data) | R1 = 0.1530, wR2 = 0.3413 | |
| Largest diff. peak and hole | 1.420 and -1.062 e.$Å^{-3}$ | |

Goodness-of-fit = $[\Sigma[w(F_o^2 - F_c^2)^2]/N_{observns} - N_{params})]^{1/2}$, all data.

$R1 = \Sigma(|F_o| - |F_c|) / \Sigma |F_o|$.   $wR2 = [\Sigma[w(F_o^2 - F_c^2)^2] / \Sigma [w(F_o^2)^2]]^{1/2}$.

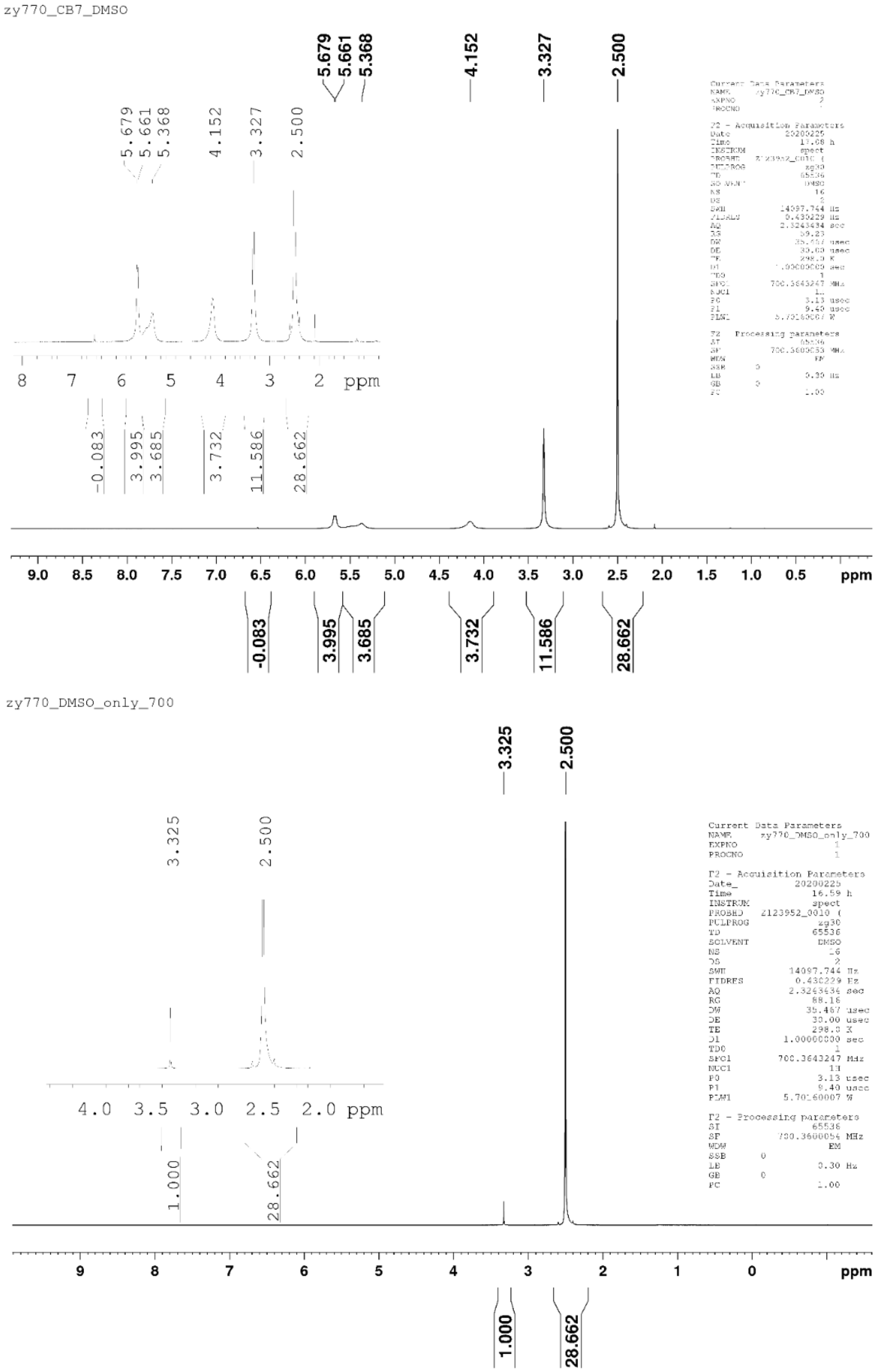

**Fig. S5.** [1]H NMR (700 MHz, DMSO-$d_6$) spectra (label: ZY770): determination of water content in CB-7. <u>Top plot</u>: CB-7 in DMSO-$d_6$. <u>Bottom plot</u>: DMSO-$d_6$ only. The data show that the integrated ratio of protons for CB-7 vs water is 11.412/(11.586 – 1.000) which corresponds to 19.5 molecules of water per CB-7 formula, $C_{42}H_{42}N_{28}O_{14}$ 19.5$H_2O$; this in turn indicates water content of 23% by weight.

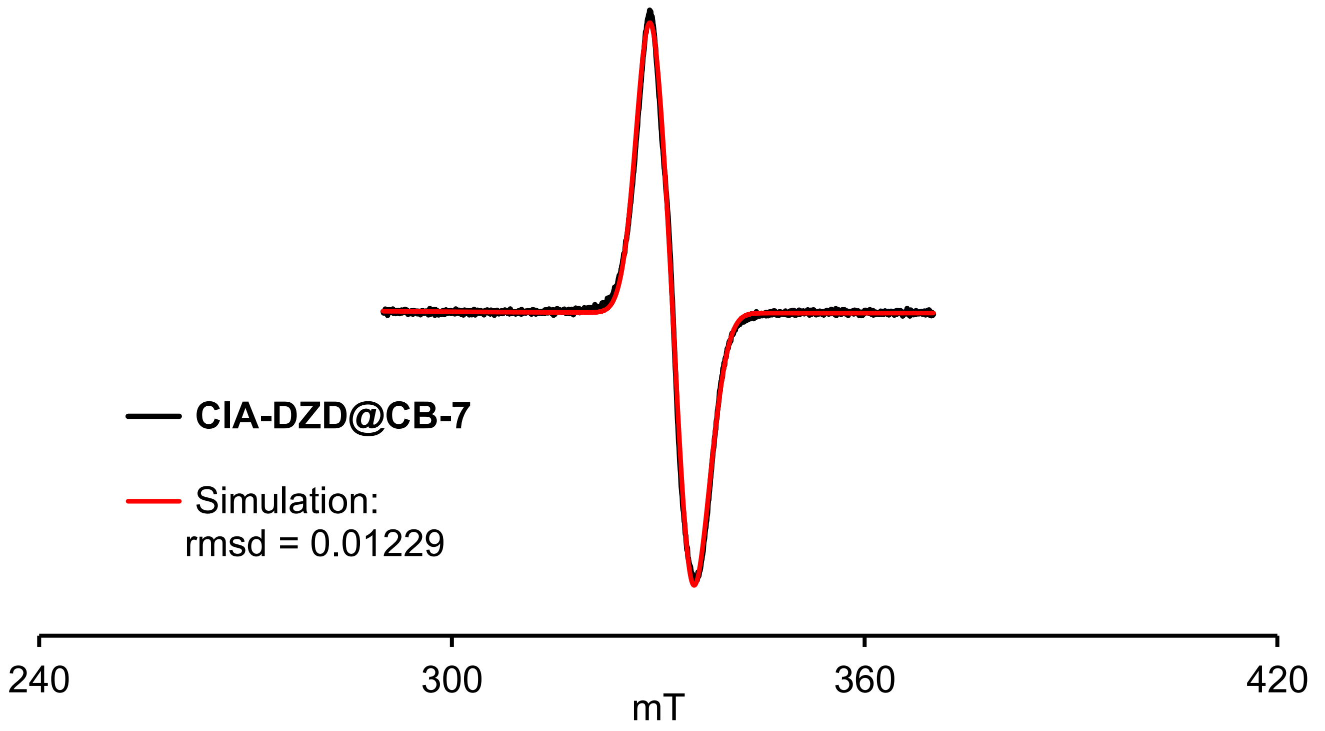

**Fig. S6.** EPR spectrum of polycrystalline **ClA-DZD@CB-7**; same data as in Figure 2C (main text). Simulation (*Pepper*)[S7] for $S$ = 1 state with the following parameters: $D$ = 120.4 MHz, $E$ = 0 MHz, $g_{xx}$ = 2.0115, $g_{yy}$ = 2.0040, $g_{zz}$ = 2.0049; strains: $H_x$ = 115.5 MHz, $H_y$ = 171.8 MHz, $H_z$ = 138.9 MHz; rmsd = 0.01229. Within the dipolar approximation, the value of $D$ = 120.4 MHz corresponds to inter-spin distance of 12.6 Å. Simulation (*Pepper*)[S7] parameters for spectrum of polycrystalline **ClA-DZD** at 294 K (Figure 3C, main text) were as follows: $S$ = 1, weight = 1.000, $D$ = 2059.7 MHz, $E$ = 79.5 MHz, $A_{xx}$ = 0.000 MHz, $A_{yy}$ = 0.009 MHz, $A_{zz}$ = 46.06, $g_{xx}$ = 2.0105, $g_{yy}$ = 2.0060, $g_{zz}$ = 2.0031; strains: $H_x$ = 90.1 MHz, $H_y$ = 130.5 MHz, $H_z$ = 37.5 MHz; $S$ = 1/2, weight = 141.4, $A_{xx}$ = 36.6 MHz, $A_{yy}$ = 0.00 MHz, $A_{zz}$ = 98.7, $g_{xx}$ = 2.0092, $g_{yy}$ = 2.0069 ($g_{yy}$ strain = 0.00592), $g_{zz}$ = 2.0029; peak-to-peak linewidths: Gaussian = 0.00833 mT and Lorentzian = 0.9052 mT; rmsd = 0.0086314.

**4. Complexation of ClA-DZD with CB-7: ITC[S13] data (Table S4, Figs. S7–S13).**

The energetics of the interactions between nitroxide ClA-DZD and CB-7 were evaluated by ITC (VP-ITC Microcal Inc., Northampton, MA). Typically, the titrant (ClA-DZD) was injected over 12 s with an interval of 300 s. After each addition, the heat was recorded. In Figs. S7 – S13 below, ITC data in purified water, phosphate buffer, and Tris buffer are shown in triplicate or duplicate.

A control experiment was carried out under identical conditions to obtain the heats of dilution and mixing involved in injection of water into the solution of CB-7 (0.2 mM) in water. The integrated heat effects of each injection were corrected by subtracting from the corresponding control experimental data. The correction was found to have negligible effect on the fitted thermodynamic parameters (Fig. S9).

**Table S4.** Isothermal Calorimetry: Association Constants ($K_a$), Enthalpy ($\Delta H$), Entropy ($\Delta S$) of ClA-DZD Complexation with CB-7 and β-CD.

| Host | Medium | $Na^+$ (mM) | pH | $N$ (sites) | $K_a$ ($M^{-1}$) | $\Delta H$ (kcal $mol^{-1}$) | $\Delta S$ (cal $mol^{-1}$ $K^{-1}$) |
|---|---|---|---|---|---|---|---|
| CB-7 | water | 0 | ~7 | 0.79±0.001 | (1.93±0.08)×$10^6$ | –13.1±0.03 | –15.3 |
| | phosphate[a] | 69 | 7.48 | 1.08±0.001 | (3.37±0.11)×$10^6$ | –13.1±0.02 | –14.2 |
| | **Tris-MOPS-1** | **53** | **7.24** | **1. 01±0.001** | **(3.74±0.07)×$10^5$** | **–10.6±0.02** | **–10.0** |
| | Tris-MOPS-2 | 3 | ~7 | 0.95±0.01 | (4.32±0.18)×$10^4$ | –9.39±0.10 | –10.3 |
| | Tris[b,c] | 0 | 7.26 | 1.02±0.002 | (1.03±0.14)×$10^5$ | –9.84±0.02 | –10.1 |
| β-CD | water | 0 | ~7 | 1.14 ± 0.02 | (7.55±0.07)×$10^2$ | –5.88±0.11 | –6.56 |

[a] 50 mM Phosphate. [b] 6 mM Tris, contains 0.01 mM EDTA (and some 0.02% (by weight) $NaN_3$). Tris-MOPS-1 and -2: 6 mM Tris/9 mM MOPS, 3-($N$-morpholino)propane-1-sulfonic acid. [c] pH is adjusted using 2.73 mM Tris and 3.27 mM Tris hydrochloride.

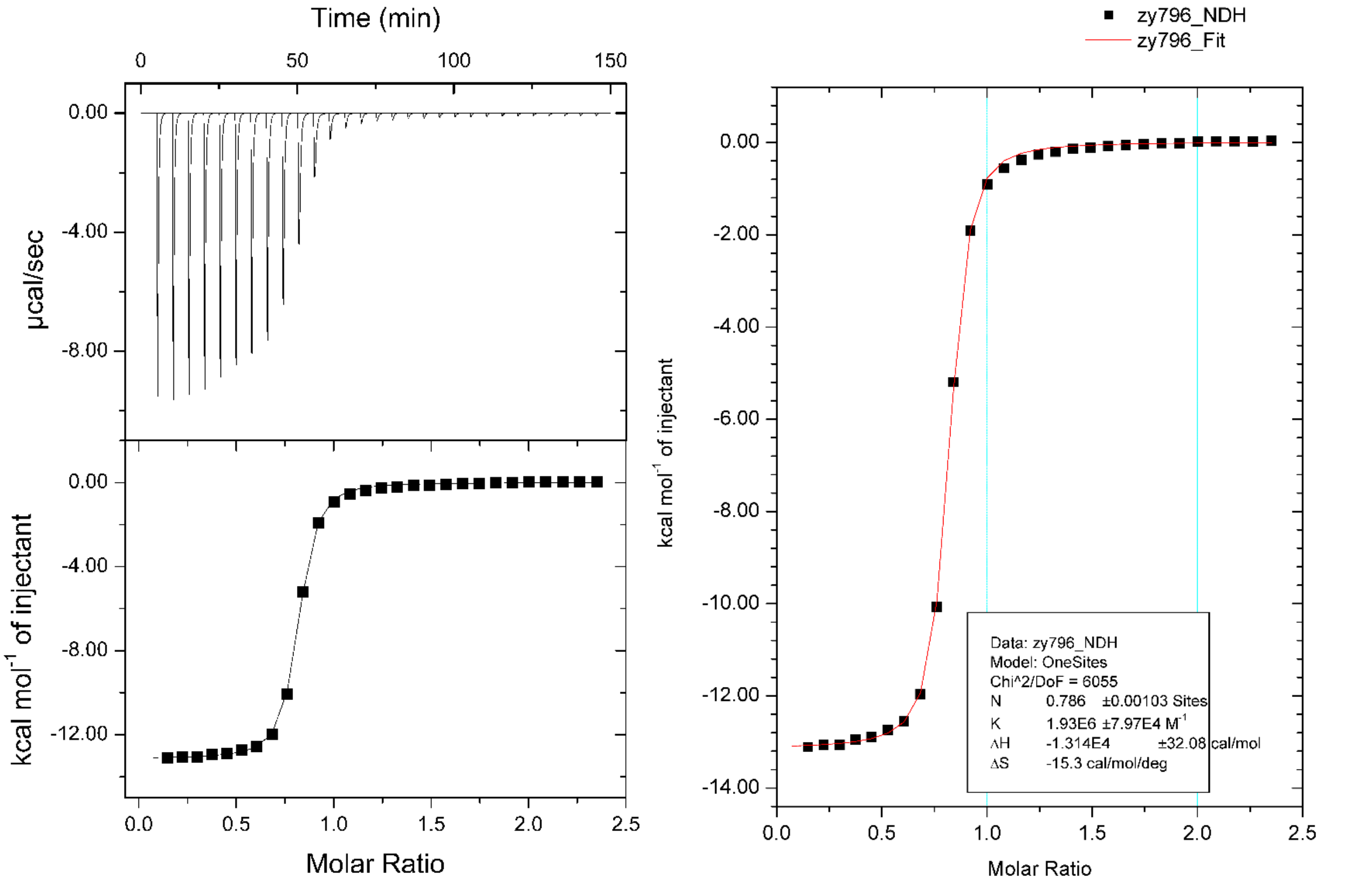

**Fig. S7.** Calorimetric titration enthalpogram (Table 1, main text; label: zy796), injection of 10 µL aliquots of 2.1 mM of nitroxide ClA-DZD (sample label: zy429_al_col, spin conc. ~100%) into the sample cell containing 0.2 mM CB-7 in purified water (pH ~7).

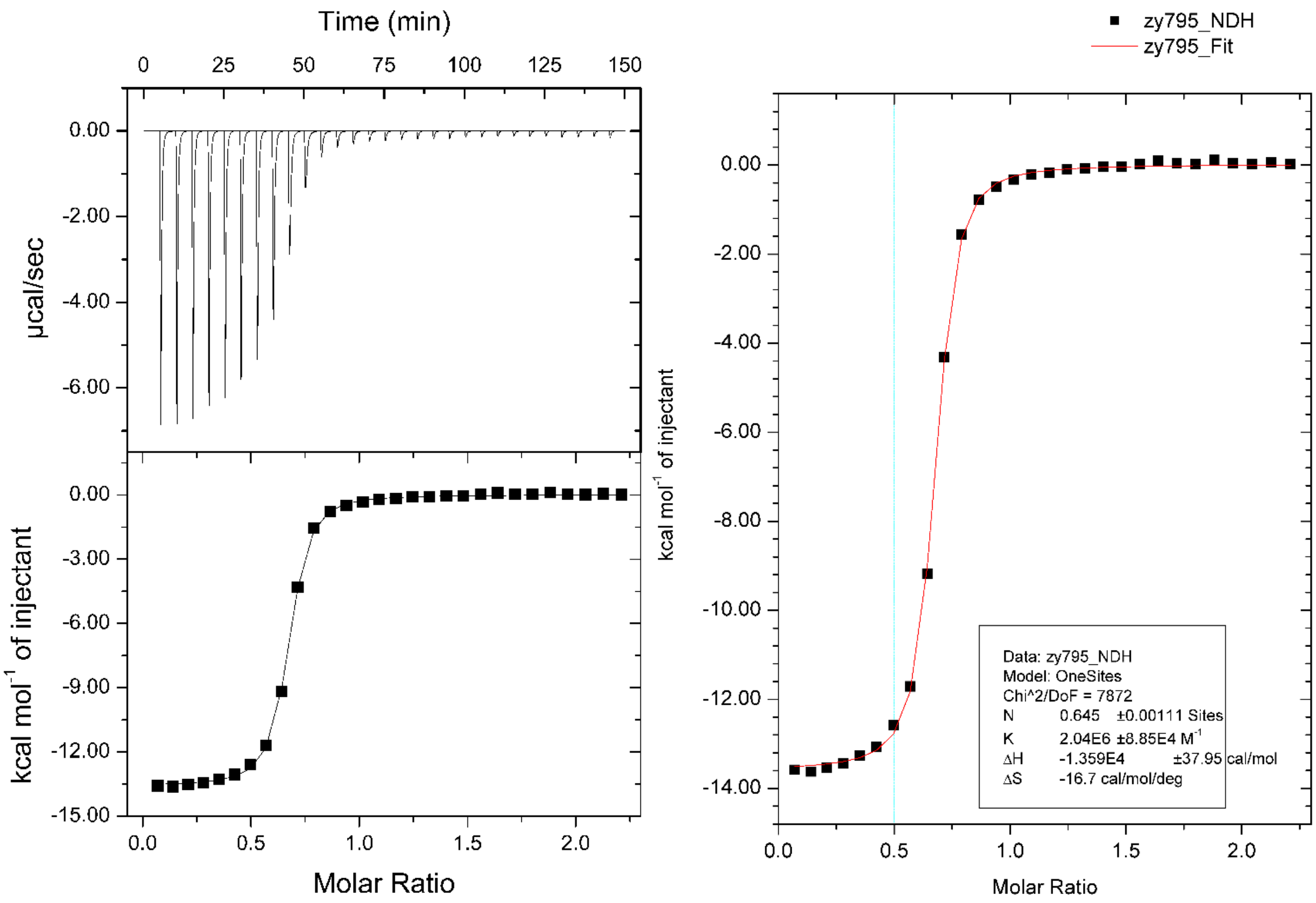

**Fig. S8.** Repeat titration in purified water. Calorimetric titration enthalpogram (label: zy795), injection of 10 µL aliquots of 2.1 mM of nitroxide ClA-DZD (sample label: zy429_al_col, spin conc. ~100%) into the sample cell containing 0.2 mM CB-7 in purified water (pH ~7).

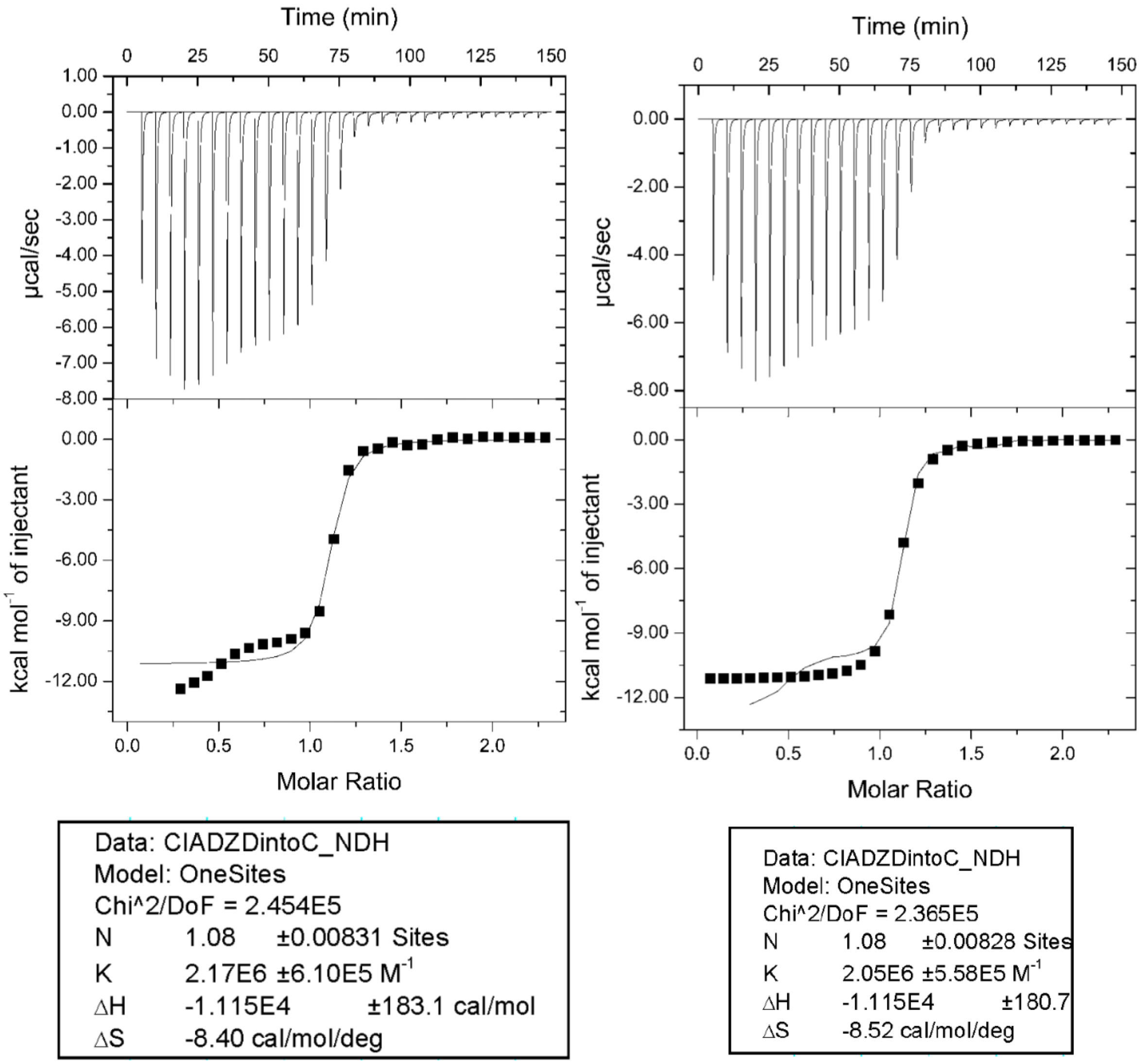

**Fig. S9.** Second repeat titration in purified water without background substraction (left plots) and with background subtraction (right plots). Calorimetric titration enthalpogram, injection of 10 μL aliquots of 2.0 mM of nitroxide ClA-DZD (sample label: zy429_al_col, spin conc. ~100%) into the sample cell containing 0.2 mM CB-7 in purified water (pH ~7).

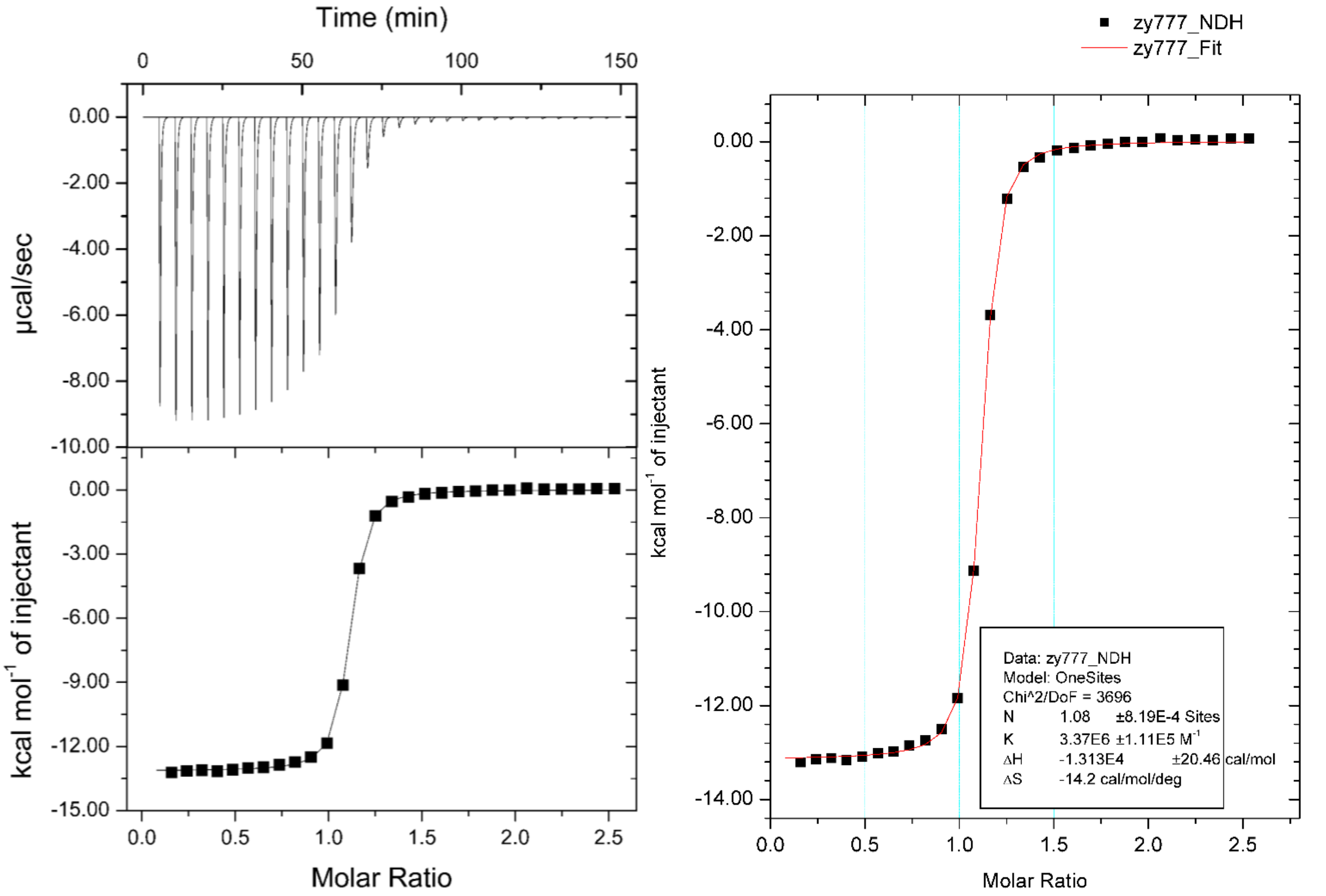

**Fig. S10.** Calorimetric titration enthalpogram (Table 1, main text; label: zy777), injection of 10 µL aliquots of 2.1 mM of nitroxide ClA-DZD (sample label: zy429_al_col, spin conc. ~100%) into the sample cell containing 0.2 mM CB-7 in 50 mM phosphate buffer (pH 7.46 – 7.48).

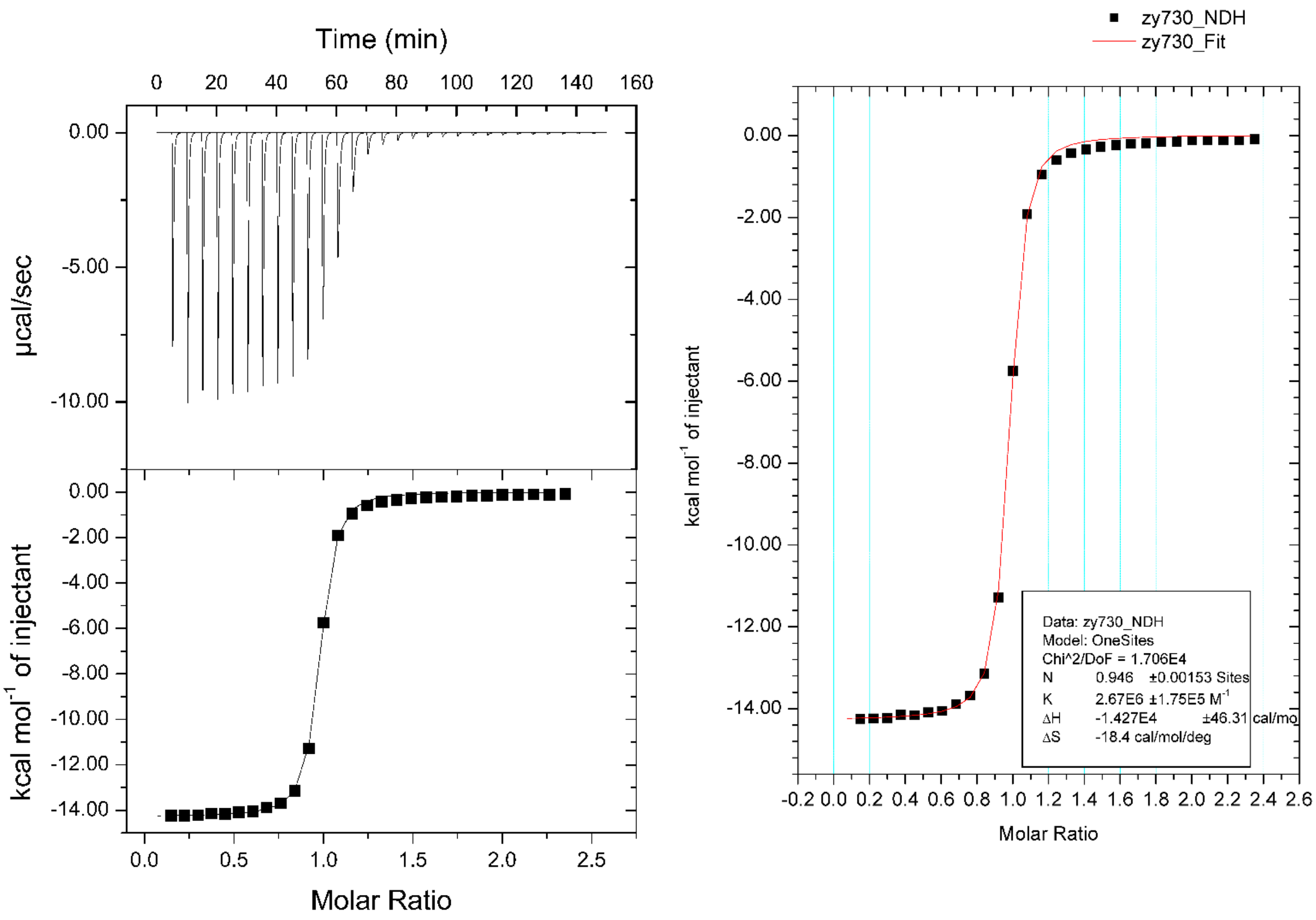

**Fig. S11.** Repeat titration in PBS buffer. Calorimetric titration enthalpogram (label: zy730), injection of 10 µL aliquots of 2.1 mM of nitroxide ClA-DZD (sample label: zy429_al_col, spin conc. ~100%) into the sample cell containing 0.2 mM CB-7 in 50 mM phosphate buffer (pH 7.46 – 7.48).

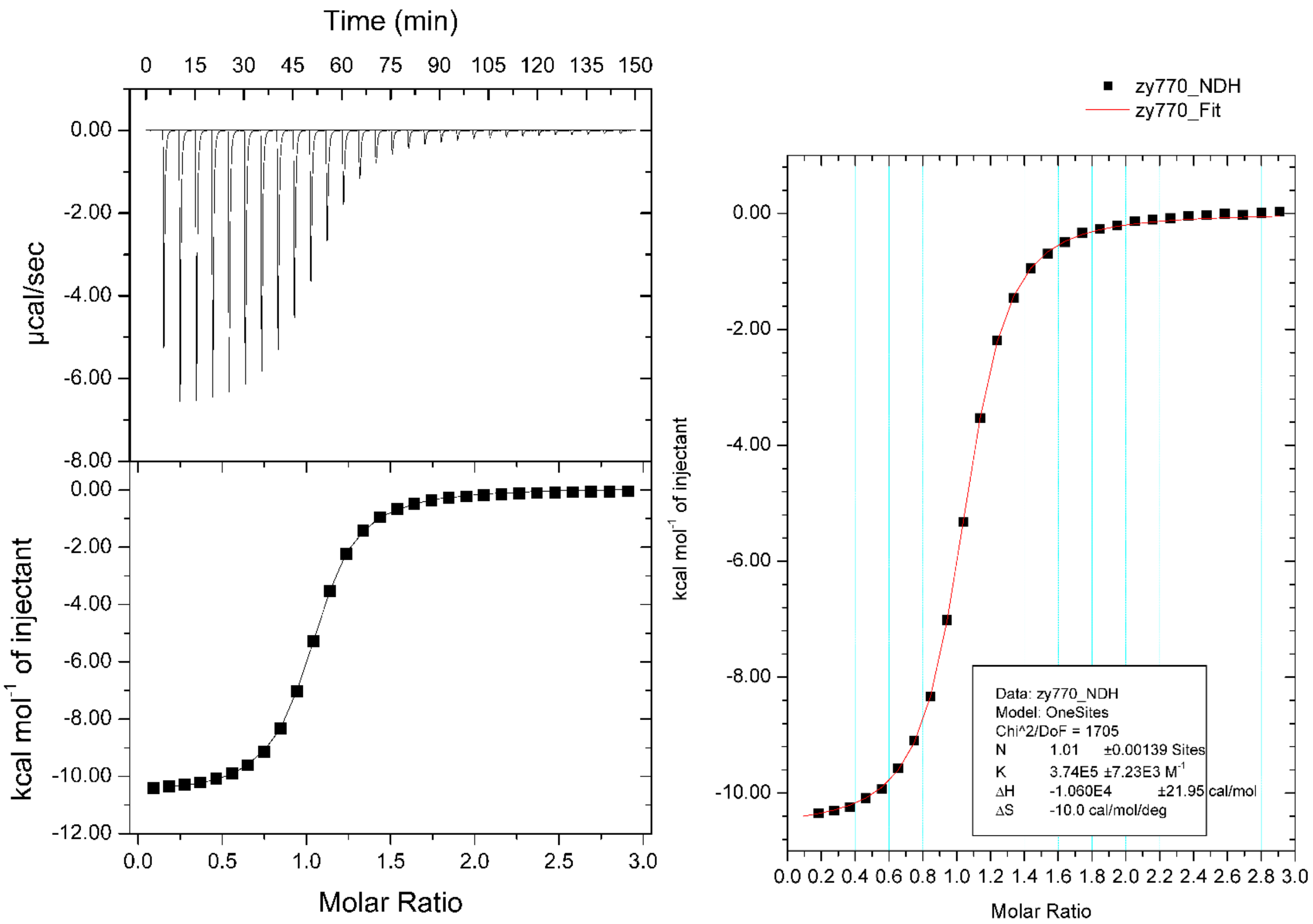

**Fig. S12.** Calorimetric titration enthalpogram (Table 1, main text; label: zy770), injection of 10 μL aliquots of 2.1 mM of nitroxide ClA-DZD (sample label: zy429_al_col, spin conc. ~100%) into the sample cell containing 0.2 mM CB-7 in 6 mM Tris buffer (9 mM MOPS and 53 mM $Na^+$, pH 7.24). This buffer is identical to that used in all EPR experiments, including DEER distance measurements.

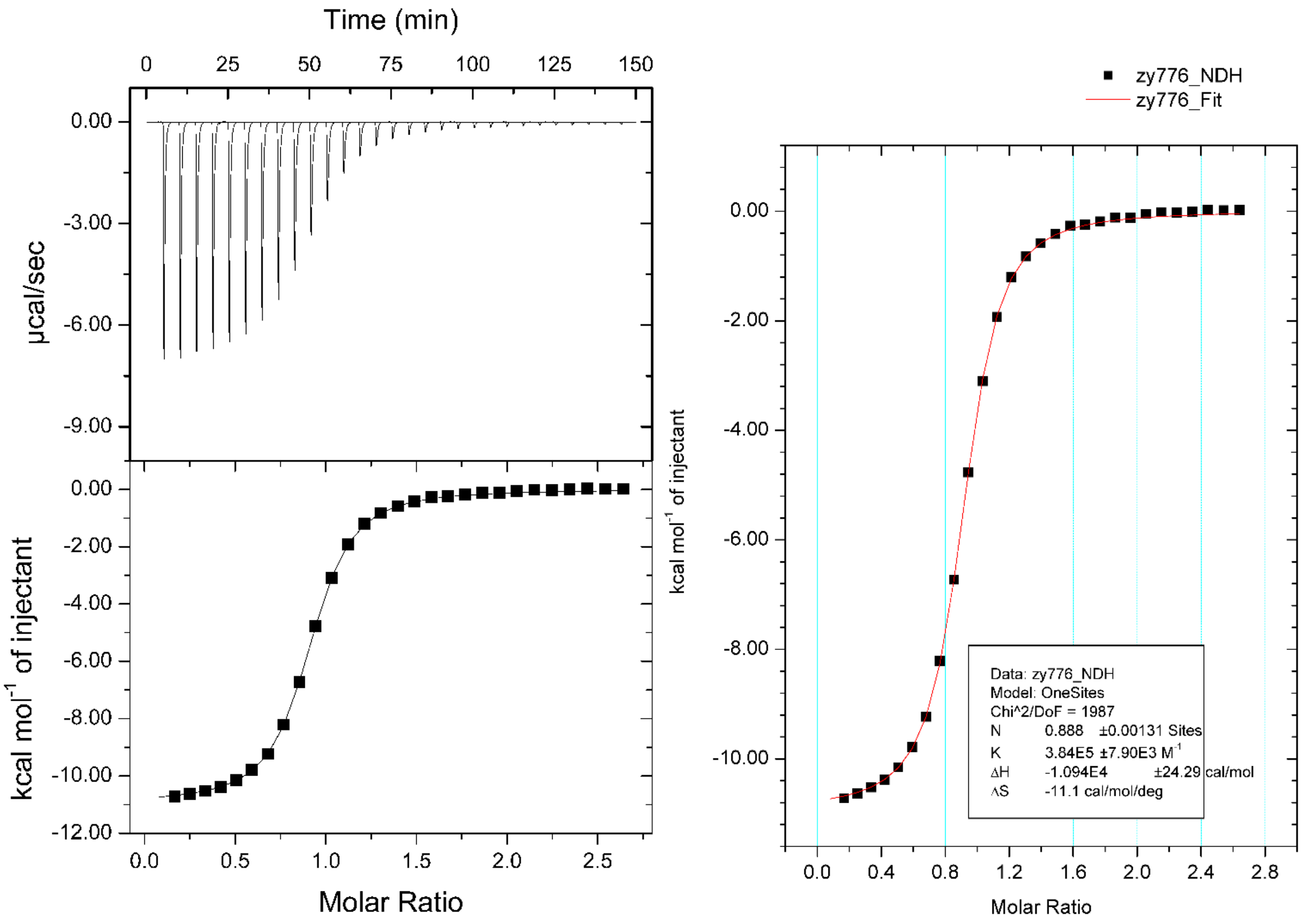

**Fig. S13.** Repeat titration in Tris buffer. Calorimetric titration enthalpogram (label: zy776), injection of 10 µL aliquots of 2.1 mM of nitroxide ClA-DZD (sample label: zy429_al_col, spin conc. ~100%) into the sample cell containing 0.2 mM CB-7 in 6 mM Tris buffer (9 mM MOPS and 53 mM $Na^+$, pH 7.24). This buffer is identical to that used in all EPR experiments, including DEER distance measurements.

**5. Complexation of ClA-DZD and DZD-T4L mutants with CB-7: CW EPR spectroscopy at Nebraska (UNL), University of Denver (DU), and Vanderbilt.** CW X-band EPR spectra for radicals were acquired on Bruker EMX-plus instrument at UNL, equipped with a frequency counter and nitrogen flow temperature control (100–400 K). The spectra were obtained in a high-sensitivity cavity. Temperatures were verified using an independently calibrated thin wire thermocouple inserted into the EPR sample tube (containing the solvent). The spectra for polycrystalline **ClA-DZD**, **ClA-DZD@CB-7** and DZD-T4L 65 were obtained at room temperature (294 K), as illustrated in Figures 2 and 3 (main text) and Figs. S1 and S6 (SI). The spectra of polycrystalline DZD-T4L 65 suspended in buffer (Fig. S1) were obtained by using two 1-mm OD quartz capillaries, sealed with beeswax at both ends, which were loaded to 5-mm OD quartz tube. The buffer was 1.6 M Na/K phosphate pH 7.4, 100 mM NaCl, 40 mM beta-mercaptoethanol.

The spectra for nitroxides **ClA-DZD** and **1** were obtained at room temperature in 6 mM Tris buffer with 10% glycerol (v/v). The buffer contained the following: 6 mM tris(hydroxymethyl)-aminomethane (Tris), 9 mM 3-(*N*-morpholino) propanesulfonic acid (MOPS), 50 mM NaCl, 0.02% sodium azide, and 0.1 mM ethylenediaminetetraacetic acid (EDTA). The spectra were recorded either without or with 2 equiv of CB-7 (Figure 4, main text). The spectra of DZD-T4L 65 and DZD-T4L 135 (both samples with spin conc. of ca. 50%) were obtained under identical conditions as described above, except 4 equiv of CB-7 were used. Analogously, the spectra for various T4L double mutants were obtained in 6 mM Tris buffer without any additional components or with 30% glycerol (i.e., w/g, 2:1), and/or with CB-7. Experimental values of $I_-/I_0$ (peak-to-peak), related to $\tau_{rot}$ are listed in Table S5.

EPR spectral simulations: simulations were carried out at DU and UNL, using the *Pepper* or *Chili* toolbox of EasySpin.[S13] Values of fitting parameters including rotational correlation times, $\tau_{rot}$, and *z*-component of $^{14}$N hyperfine *A*-tensor, $A_{zz}$, are summarized in Figs. S1 and S6, and in Table S5. The starting points for the simulations of fluid solution spectra were based on values of *g* and *A*, obtained by fitting the field-swept echo detected spectra at Q-band that are shown in section 6. Small changes, primarily $A_{zz}$, were made to match the fluid solution spectra.

**Table S5.** Summary of simulations of fluid solution EPR spectra using *Chili* toolbox of EasySpin.

| Sample | $A_{xx}$ [MHz] | $A_{yy}$ [MHz] | $A_{zz}$ [MHz] | $g_{xx}$ | $g_{yy}$ | $g_{zz}$ | Lwpp [mT] | $\tau_{rot}$ [ns] | $I_-/I_0$ | $A_{iso}$ [MHz] | Fig. |
|---|---|---|---|---|---|---|---|---|---|---|---|
| **1** | 15.4 | 18 | 103 | 2.0099 | 2.0060 | 2.0022 | 0.10 | 0.0102 | 0.91 | 45.5 | 3 (main text) |
| **1**; CB-7 (2 equiv) | 15.4 | 18 | 103 | 2.0099 | 2.0060 | 2.0022 | 0.10 | 0.0102 | 0.93 | 45.5 | |
| **ClA-DZD** | 20 | 34 | 108 | 2.0094 | 2.0062 | 2.0022 | 0.063 | 0.0032 | 0.97 | 54.0 | |
| **ClA-DZD**; CB-7 (2 equiv) | 20 | 35 | 105 | 2.0102 | 2.0070 | 2.0027 | 0.044 | 0.1150 | 0.41 | 53.3 | |
| DZD-T4L 65/80 | 22 | 35 | 112 | 2.0094 | 2.0060 | 2.0022 | 0.51 | 1.76 | 0.38 | 56.3 | S14 (SI) |
| DZD-T4L 65/80; 1 mM CB-7 | 21 | 38 | 108 | 2.0095 | 2.0060 | 2.0022 | 0.45 | 2.00 | 0.36 | 55.7 | |
| DZD-T4L 65 | 23 | 27 | 116 | 2.0094 | 2.0062 | 2.0026 | 0.54 | 1.41 | 0.41 | 55.3 | S15 (SI) |
| DZD-T4L 65; 4 equiv of CB-7 | 26 | 30 | 107 | 2.0108 | 2.0062 | 2.0020 | 0.52 | 1.62 | 0.37 | 54.6 | |
| DZD-T4L 135 | 26 | 27 | 113 | 2.0093 | 2.0057 | 2.0027 | 0.56 | 1.33 | 0.48 | 55.0 | S16 (SI) |
| DZD-T4L 135; 4 equiv of CB-7 | 28 | 29 | 106 | 2.0101 | 2.0061 | 2.0022 | 0.56 | 1.62 | 0.39 | 54.5 | |

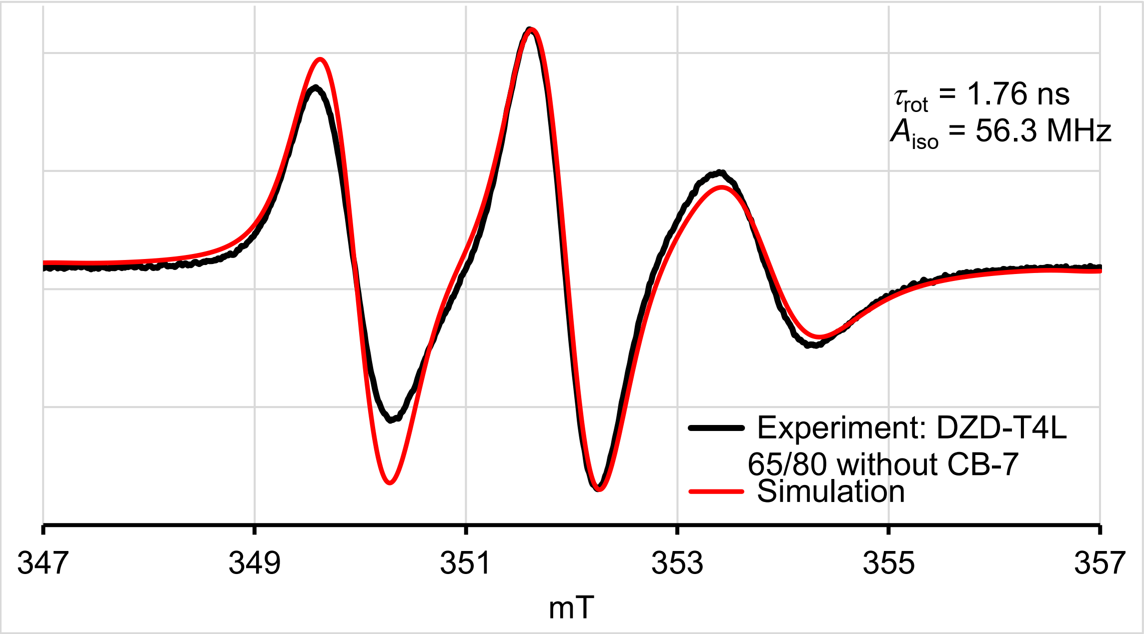

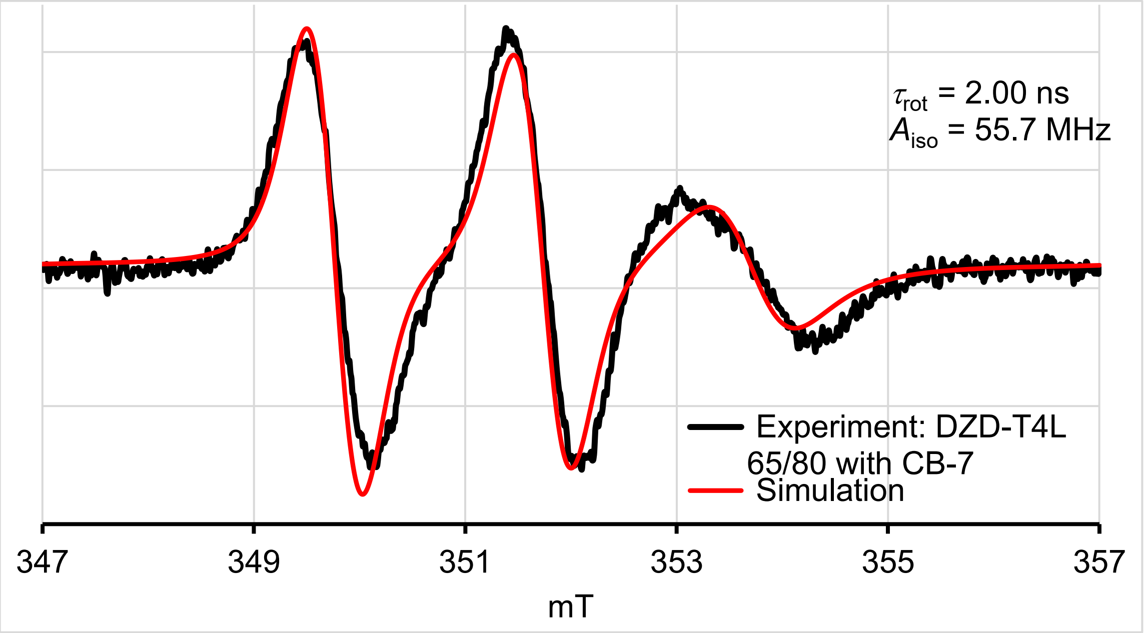

**Fig. S14.** EPR (X-Band, ν = 9.437 – 9.438 GHz) spectra at 295 K of doubly spin labeled DZD-T4L 65/80 in 30% glycerol in water v/v (6 mM Tris buffer). Black and red lines correspond to the experimental data and spectral simulation, respectively. Top plot: prior to addition of CB-7, rotational correlation time, $\tau_{rot}$ = 1.76 ns, $A_{zz}$ = 112 MHz, line width peak-to-peak, lwpp = 0.51 mT. Bottom plot: after addition of CB-7 to obtain 1 mM solution in CB-7, $\tau_{rot}$ = 2.00 ns, $A_{zz}$ = 108 MHz, lwpp = 0.45 mT. (Parameters used in the simulations are listed in Table S4.)

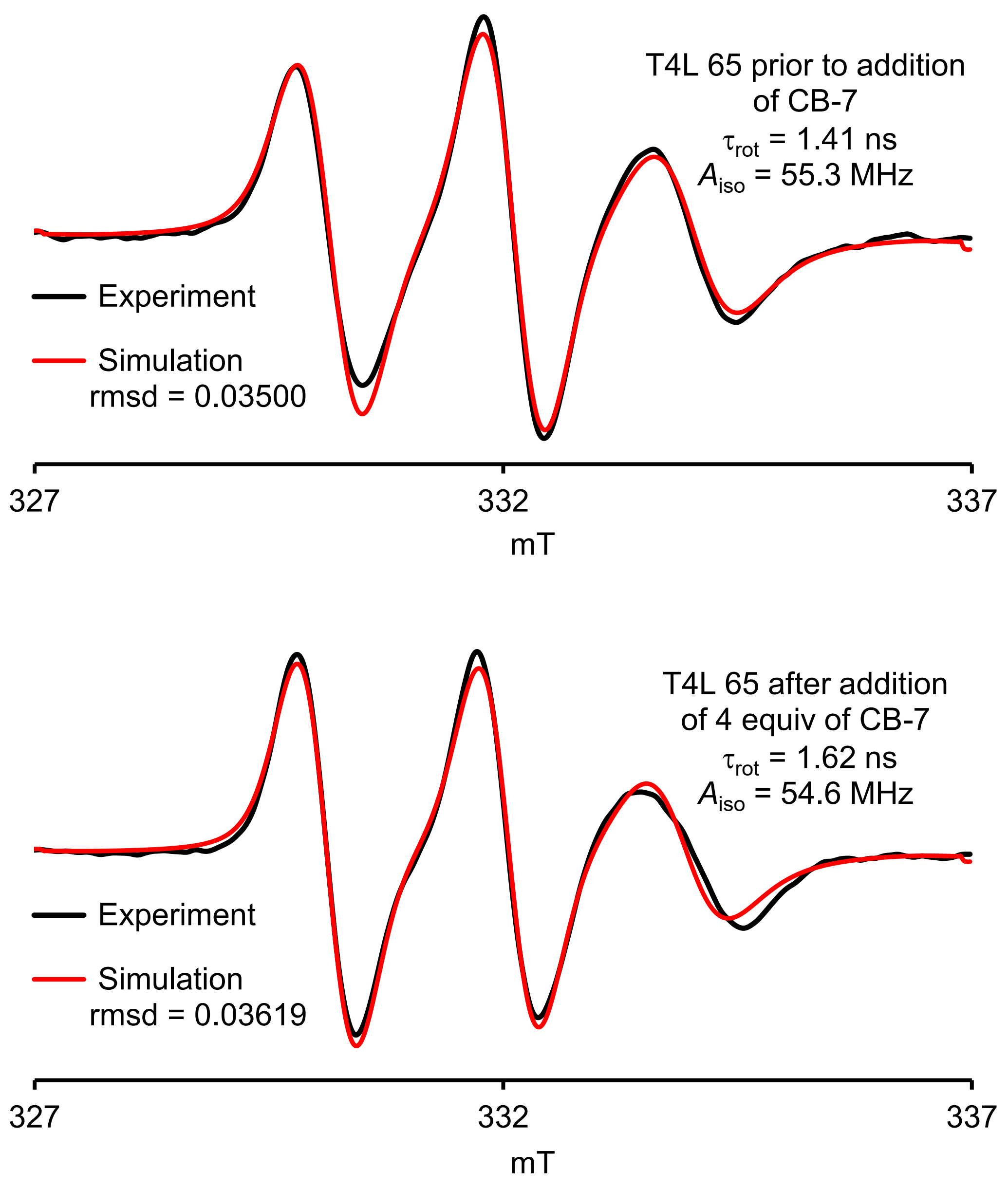

**Fig. S15.** EPR (X-Band, ν = 9.3259 and 9.3255 GHz) spectra at 295 K of mono spin labeled DZD-T4L 65 in 10% glycerol in water v/v (6 mM Tris buffer). Black and red lines correspond to the experimental data and spectral simulation, respectively. Top plot: prior to addition of CB-7, rotational correlation time, $\tau_{rot}$ = 1.41 ns, $A_{zz}$ = 116 MHz, line width peak-to-peak, lwpp = 0.54 mT.; spin conc. ca. 50%. Bottom plot: after addition of 4 equiv of CB-7, $\tau_{rot}$ = 1.62 ns, $A_{zz}$ = 107 MHz, lwpp = 0.52 mT. (Parameters used in the simulations are listed in Table S5.)

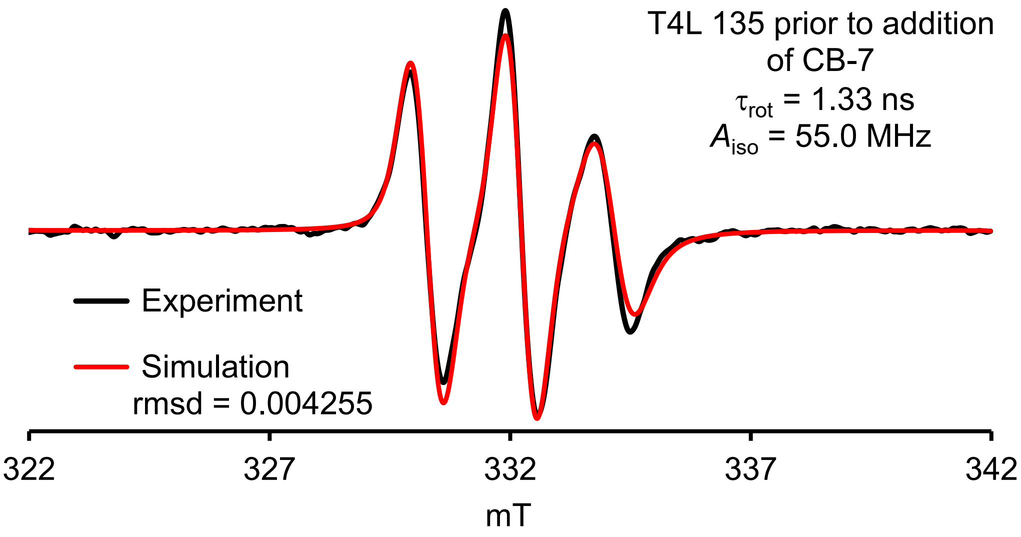

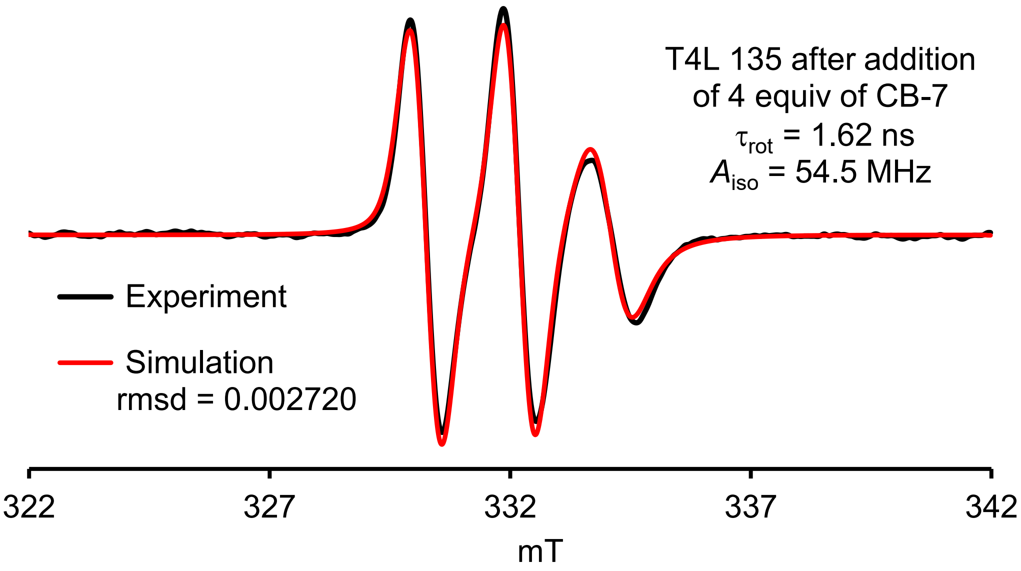

**Fig. S16.** EPR (X-Band, ν = 9.3281 and 9.3285 GHz) spectra at 295 K of mono spin labeled DZD-T4L 135 in 10% glycerol in water v/v (6 mM Tris buffer). Black and red lines correspond to the experimental data and spectral simulation, respectively. Top plot: prior to addition of CB-7, rotational correlation time, $\tau_{rot}$ = 1.33 ns, $A_{zz}$ = 113 MHz, line width peak-to-peak, lwpp = 0.56 mT.; spin conc. ca. 50%. Bottom plot: after addition of 4 equiv of CB-7, $\tau_{rot}$ = 1.62 ns, $A_{zz}$ = 106 MHz, lwpp = 0.56 mT. (Parameters used in the simulations are listed in Table S4.)

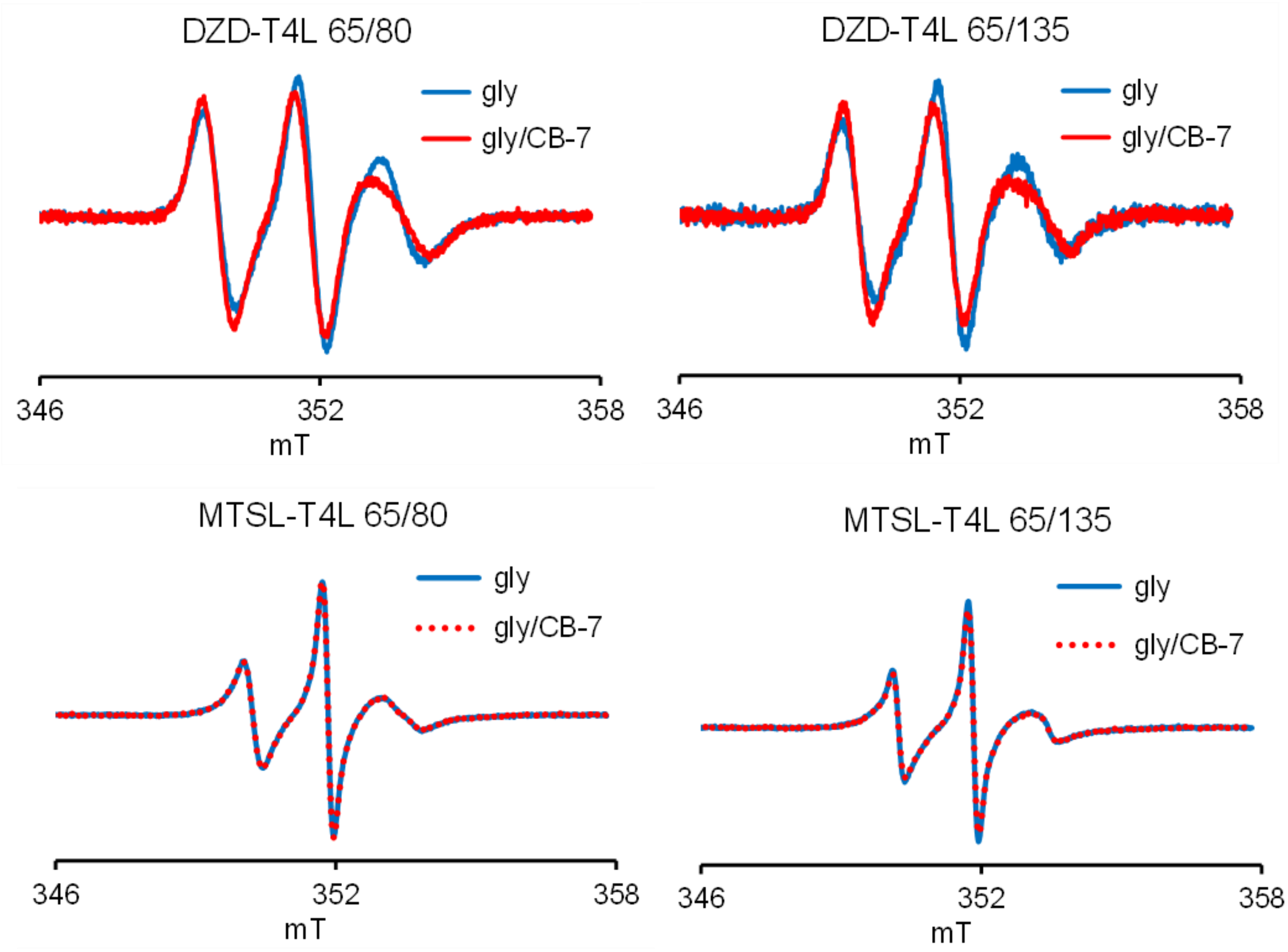

**Fig. S17.** CW EPR (X-band) spectra for T4L mutants prior to the DEER measurements shown in Figure 5 (main text). The spectra are consistent with increased rotational correlation times ($\tau_{rot}$) after addition of 1 mM CB-7 to DZD-T4L mutants and the absence of change in $\tau_{rot}$ for MTSL-T4L mutants.

### 6. Field-swept echo-detected spectra.

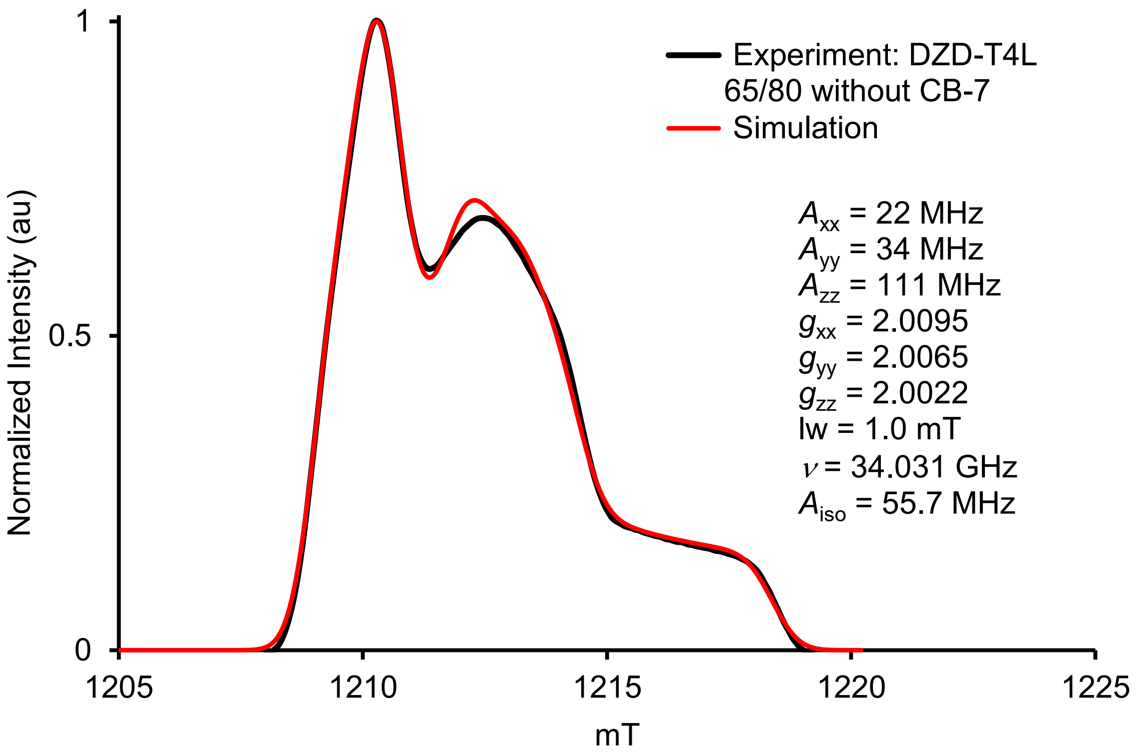

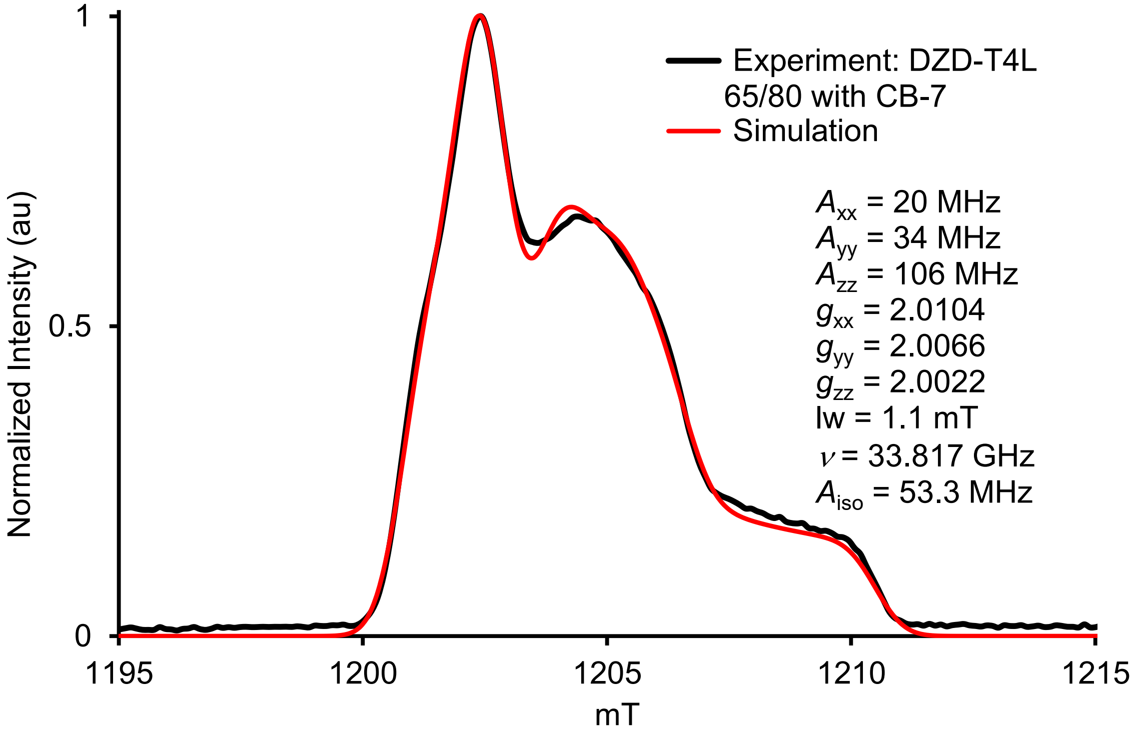

**Fig. S18.** Field-swept echo-detected spectra of DZD-T4L 65/80 in 30% glycerol in water (6 mM Tris buffer) without CB-7 (bottom plot) and with 1 mM CB-7 (top plot) at 80 K at Q-band.

**7. Electron spin relaxation studies on spin labels and doubly spin labeled DZD-T4L.**

**Stretch exponential fits for relaxation data.** The stretched exponential gives better fits to the echo dephasing and the stretch parameter is a useful indicator of mobility. The disadvantage of the stretch exponential fits is that by having two adjustable parameters there is more scatter in the resulting values. For the variable temperature experiments we typically do not record multiple data sets at each temperature so we do not have replicate measurements to use in calculating uncertainties. For single exponential fits the reproducibility in $T_m$ values typically is about 5%. Uncertainties are larger for the values obtained by stretched exponential fits. Typically, stretched exponential fits will provide larger values of $T_m$, compared to single exponential fits, as illustrated in Fig. S28.

**Electron spin relaxation studies of spin labels**. Variable temperature $T_m$ and $T_1$ relaxation measurements of 0.95 mM IA-DZD in 1:1 water:glycerol (v/v), 0.95 mM IA-DZD + 1.4 mM CB-7 in 1:1 water:glycerol and 0.5 mM MTSL in 1:1 water:glycerol were performed at X band (9.702 GHz). Each sample was transferred into a 4 mm OD quartz tube. Several freeze-pump-thaw cycles to remove oxygen were performed and each sample was flame-sealed under a partial pressure of helium gas (~100 mtorr) to facilitate thermal equilibrium. Since no orientation dependence was observed, 2-pulse echo decays to measure $T_m$ and 3-pulse inversion recoveries to measure $T_1$ were collected at the center line (maximum peak) of the EDFS. The possibility of contributions to the echo decays from instantaneous diffusion was checked by measuring echo decays with different pulse lengths, and no change in $T_m$ was observed. Unless otherwise noted, the parameters for the pulse experiment were $\pi/2$ = 40 ns with 15 dB attenuation of the output of the TWT, and initial $\tau$ = 200 ns. Other parameters such as time increment, shots per point, number of scans, short repetition time and gain are adjusted at every temperature depending on the level of signal to noise and anticipated length of $T_m$ and $T_1$.

Two representative echo detected field swept spectra (EDFS) are shown in Fig. S19, which are representative of IA-DZD and IA-DZD+CB7 at 80 K. The spectra are very similar, indicating no differences in $g$ value. The value of $A_{zz}$ is slightly smaller in the presence of CB-7, as has been observed in other experiments. The spectra are consistent with immobilized nitroxide.

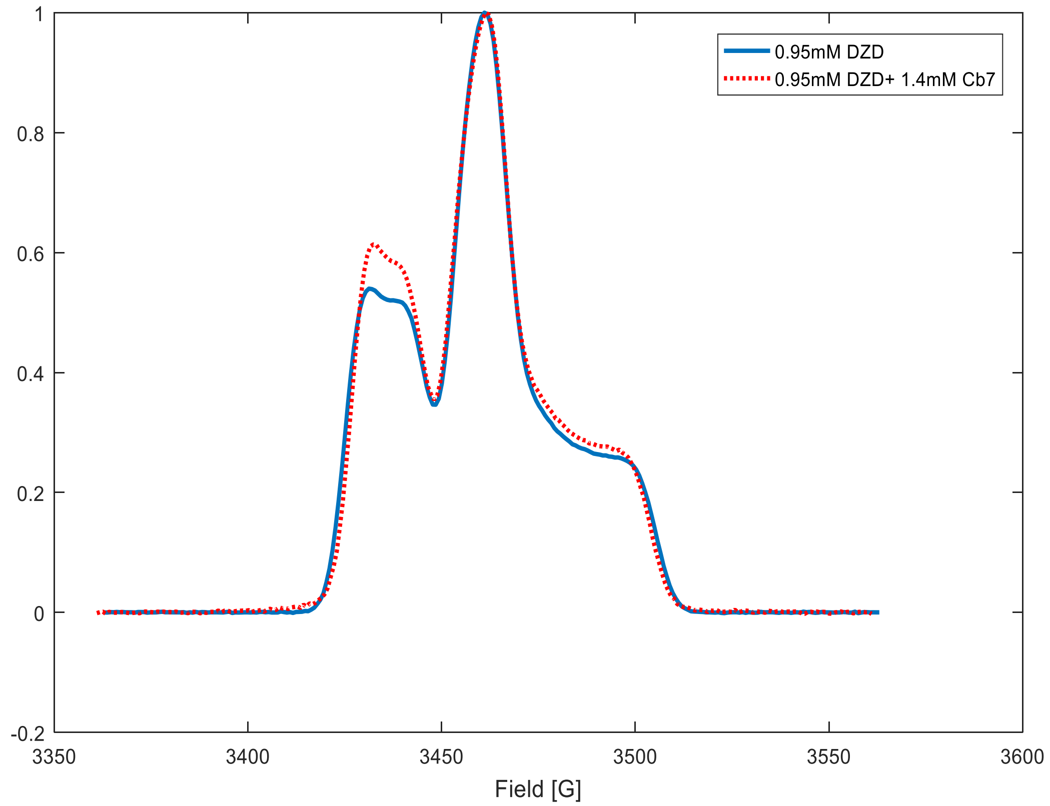

**Fig. S19.** Echo detected field swept spectra for 0.95 mM IA-DZD and 0.95 mM IA-DZD+1.4 mM CB-7.

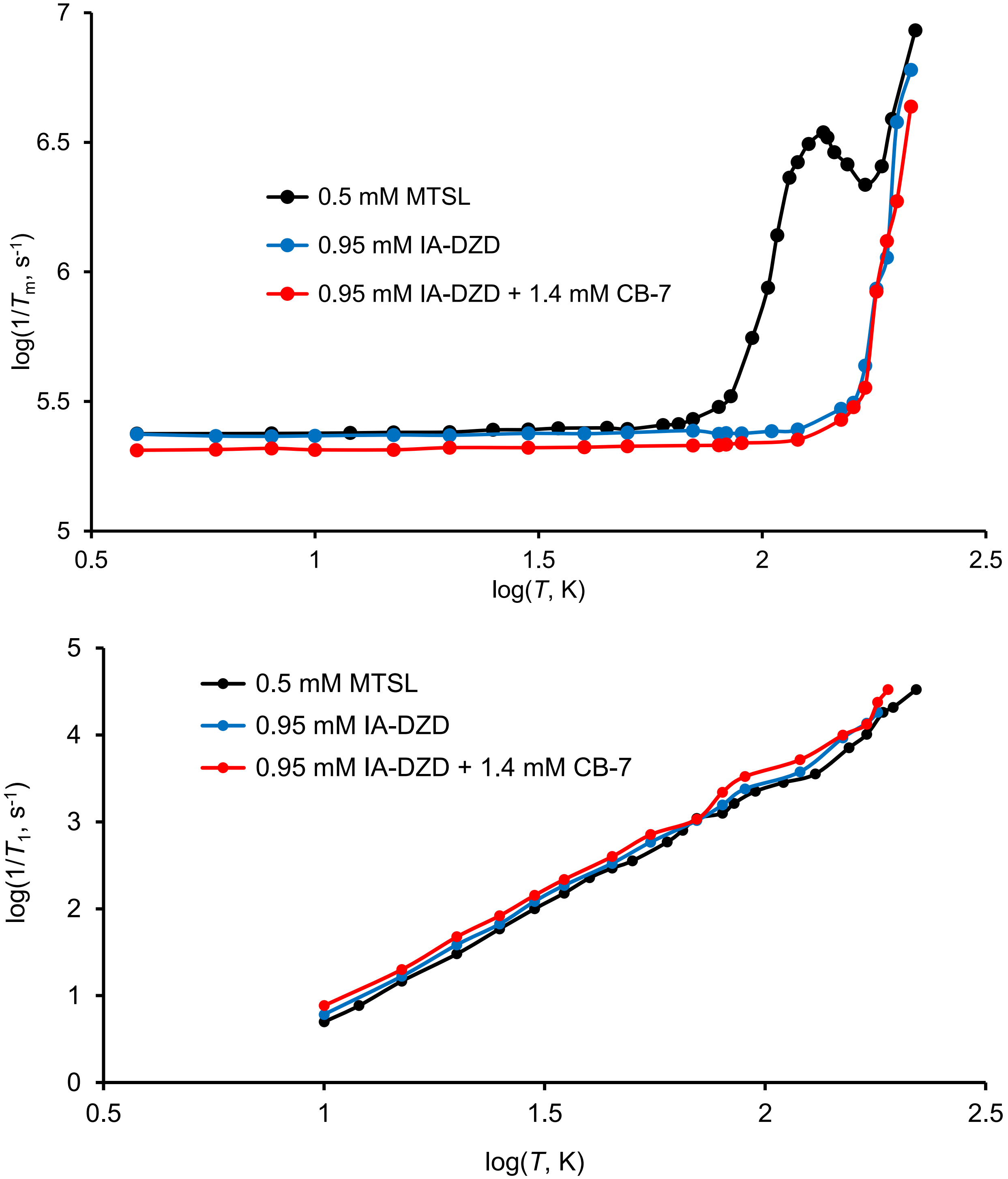

**Fig. S20.** Temperature dependence of $1/T_m$ (top plots) and $1/T_1$ (bottom plots) for 0.5 mM MTSL, 0.95 mM IA-DZD, and 0.95 mM IA-DZD +1.4 mM CB-7 1:1 glycerol/water at X-Band ($\nu = 9.702$ GHz) recorded at the maximum of the EDFS shown in the preceding Fig. For $T_1$, data below 10 K are not included because nitroxides have very long $T_1$ at cryogenic temperatures and it becomes very difficult to obtain a window long enough to accurately measure the recovery below about 10 K.

**Electron spin relaxation studies of double spin labeled DZD-T4L.** Orientation dependence of $T_m$ and $T_1$ was measured on frozen glassy 65-135 DZD, 65-135 DZD+CB-7, 65-80 DZD and 65-80 DZD+CB-7 at 80 K and 150 K at relevant positions in the spectrum. Data for 65-135 DZD were acquired at 130 K because the sample was small and the S/N was poor at higher temperature because there was some loss when the sample was transferred from a broken tube. A representative Q-band 2 pulse echo detected spectrum is presented in Fig. S21.

$T_m$ was measured by two pulse echo decay and $T_1$ was measured by inversion recovery with $\pi/2$ = 20 ns pulses. $T_m$ was calculated by fitting the data to a stretched exponential. In fitting these data with two different programs we realized that the Bruker software defines the first time in the stored data array as t = 0. This does not take account of the fact that the initial time is actually d1, where d1= time delay after application of the first pulse. This is a source of systematic error, but is unlikely to impact trends if d1 is constant.

In Fig. S24, it is shown that $1/T_m$ is slightly slower for 65-135 DZD+CB-7 than 65-135 DZD at 80 K and is nearly constant across the orientations. $1/T_m$ for 65-135 DZD+CB-7 at 150 K is slower than for 65-135 DZD at 130 K, despite the higher temperature. At 150 K, $1/T_m$ for 65-135 DZD+CB-7, becomes more dependent on position in the spectrum at intermediate positions of the field with respect to the molecular *z* axis, which is the region of the spectrum that is most strongly motion dependent. The shapes of the echo decays are significantly different at 80 K (Fig. S25), which is reflected in the differences in the stretch parameter, $\beta$, and $T_m$.

**Table S6.** Summary of $T_1$ and $T_m$ for selected CB-7 complexes at Q band.

| T4L in buffer:gly | $T_m$ (μs) and [β] | | $T_1$ (μs)[a] | | $T_1$ (μs)[b] | | $\beta$ (for $T_1$) | |
|---|---|---|---|---|---|---|---|---|
| | 80 K | 150 K | 80 K | 150 K | 80 K | 150 K | 80 K | 150 K |
| 65-80DZD | 4.16 [2.12] | 3.20 [1.59] | 460 | 115 | 261 | 64 | 0.72 | 0.74 |
| [c]65-80 DZD+CB-7 | 4.97 [2.33] | 4.62 [2.08] | 815 | 195 | 481 | 124 | 0.74 | 0.77 |
| 65-135 DZD | 4.20 [2.03] | 3.63 [1.63] | 438 | 160 | 218 | 85 | 0.70 | 0.72 |
| 65-135DZD+CB-7 | 4.94 [2.38] | 4.27 [1.89] | 836 | 206 | 510 | 365 | 0.70 | 0.73 |

[a] Long component in a fit to two exponentials. [b] $T_1$ from stretched exponential and corresponding $\beta$.

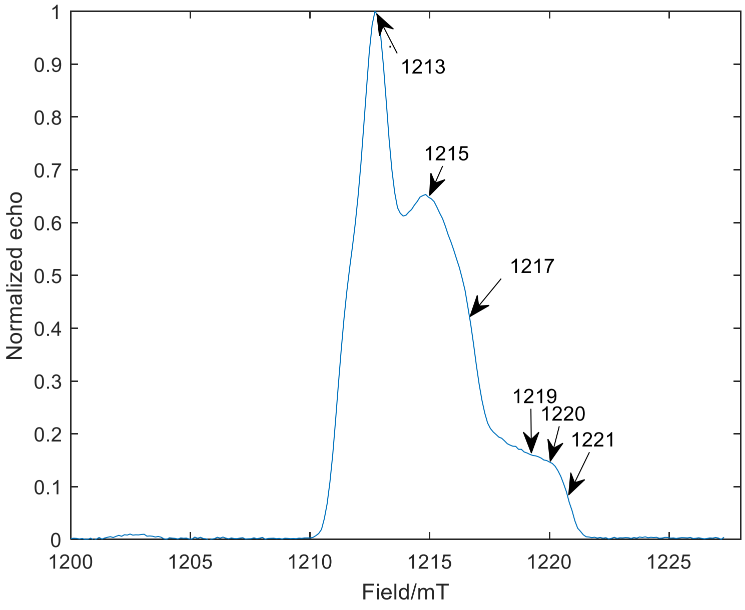

**Fig. S21.** 2-Pulse Q band echo detected spectrum for DZD-T4L 65/80: the Q band mw frequency: 33.965 GHz. The π/2 pulse was 20 ns. $T_m$ was measured at 1213 mT, which corresponds to *x* axis, 1215 mT (*y* axis), 1221 mT (*z* axis) and at intermediate positions shown in the figure.

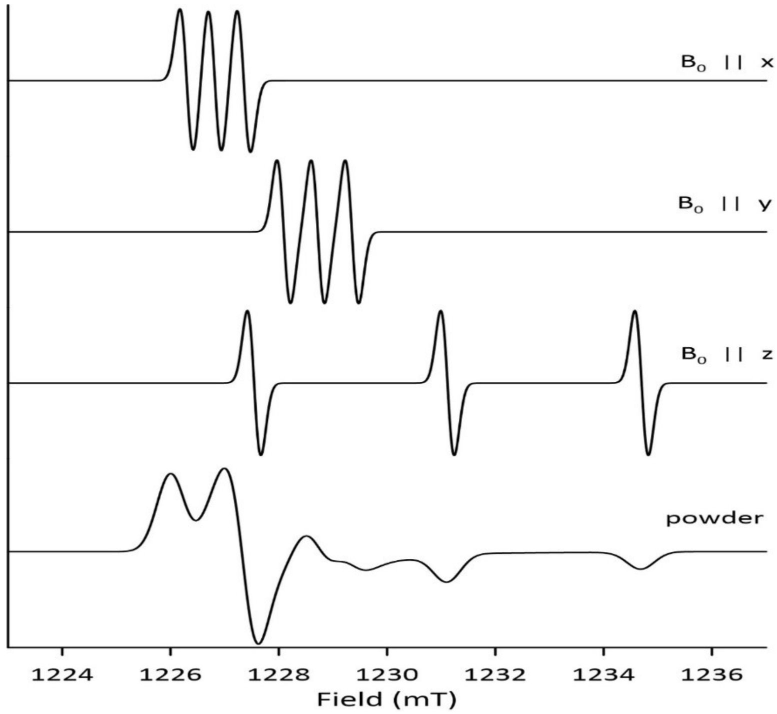

**Fig. S22.** Simulated spectra for Q Band single crystal and powder spectra that show the relationship between spectra for the magnetic field along the principal axes and the powder spectrum. The calculation was done for a higher microwave frequency than in the preceding figure, so the field axis is offset in that figure.

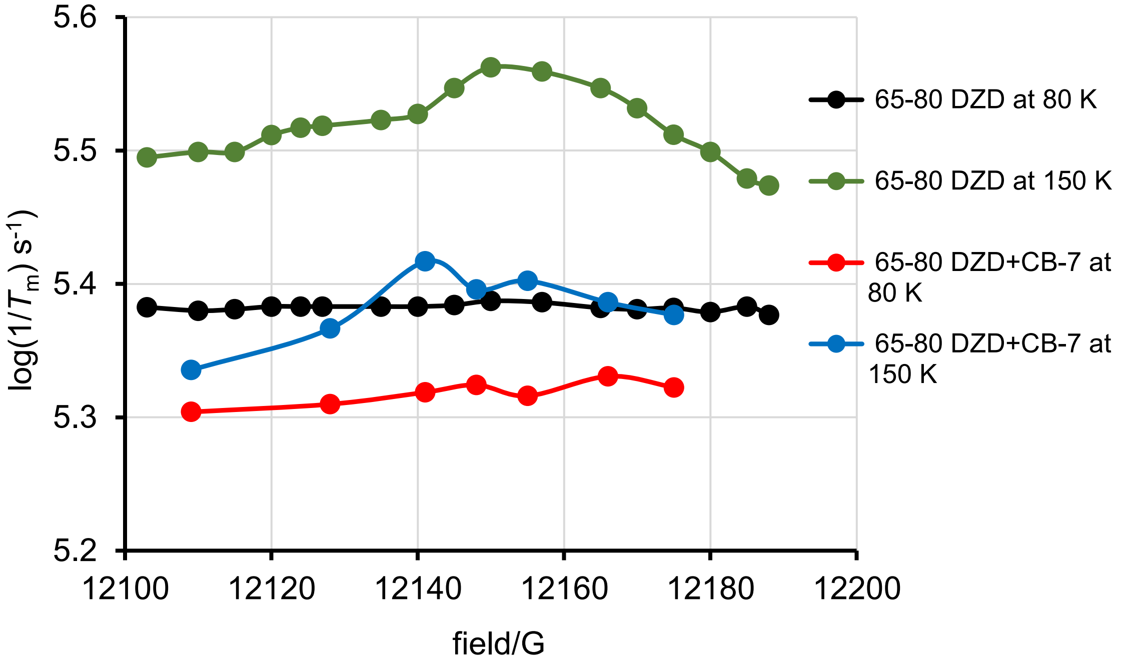

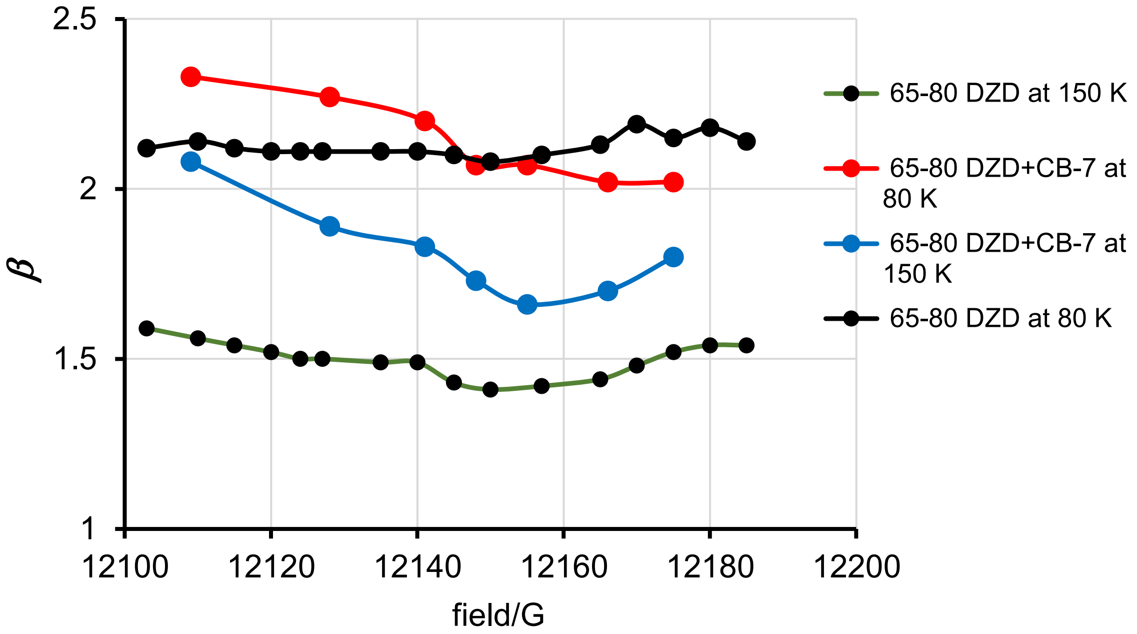

**Fig. S23.** Field and temperature dependence for doubly spin labeled DZD-T4L 65/80 in 30% glycerol in water without and with 1 mM CB-7 at Q-band. _Top panel:_ dependence of $1/T_m$. _Bottom panel:_ dependence of stretch parameter $\beta$.

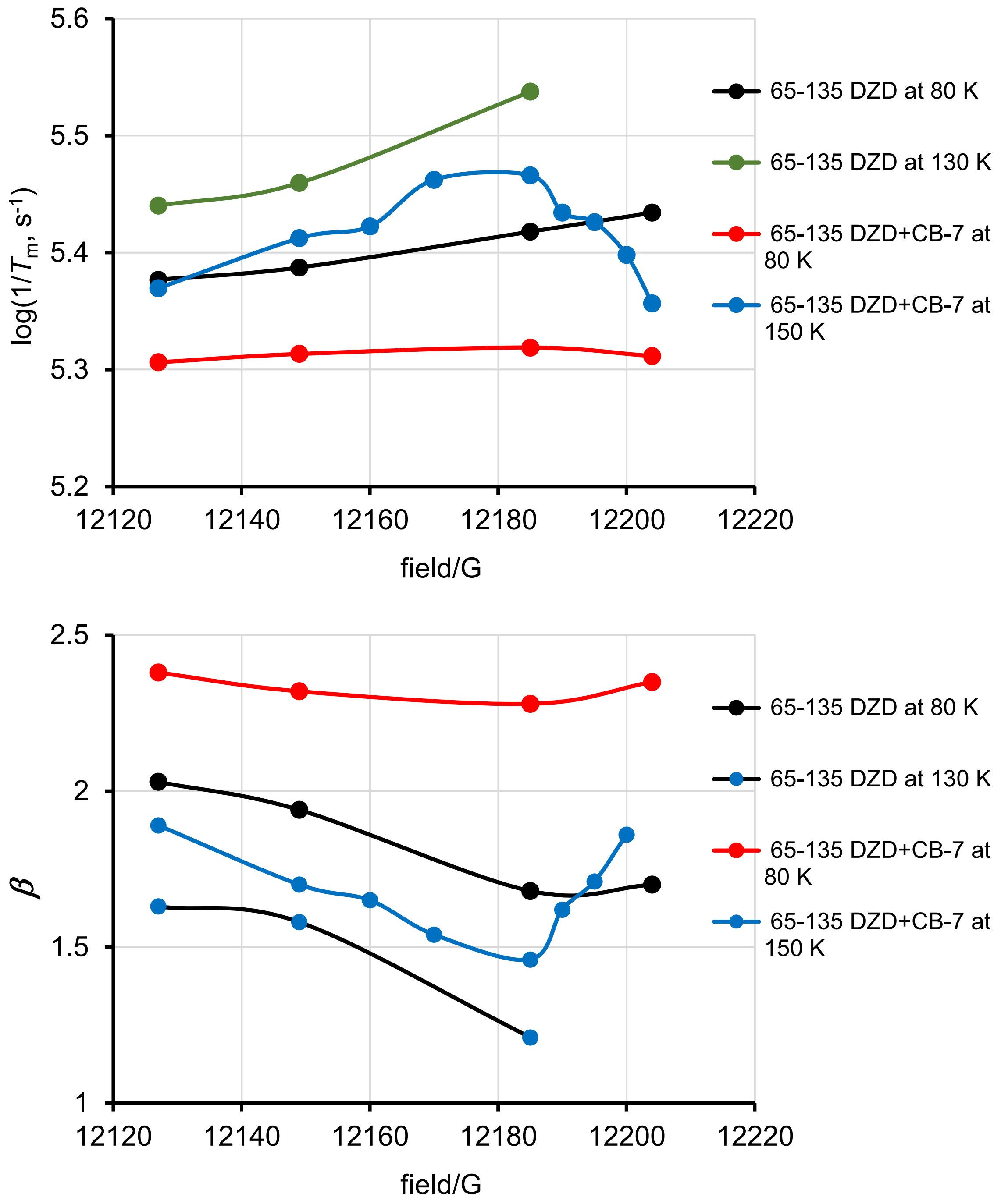

**Fig. S24.** Field and temperature dependence for doubly spin labeled DZD-T4L 65/135 in 30% glycerol in water without and with 1 mM CB-7 at Q-band. *Top panel:* dependence of $1/T_m$. *Bottom panel:* dependence of stretch parameter $\beta$.

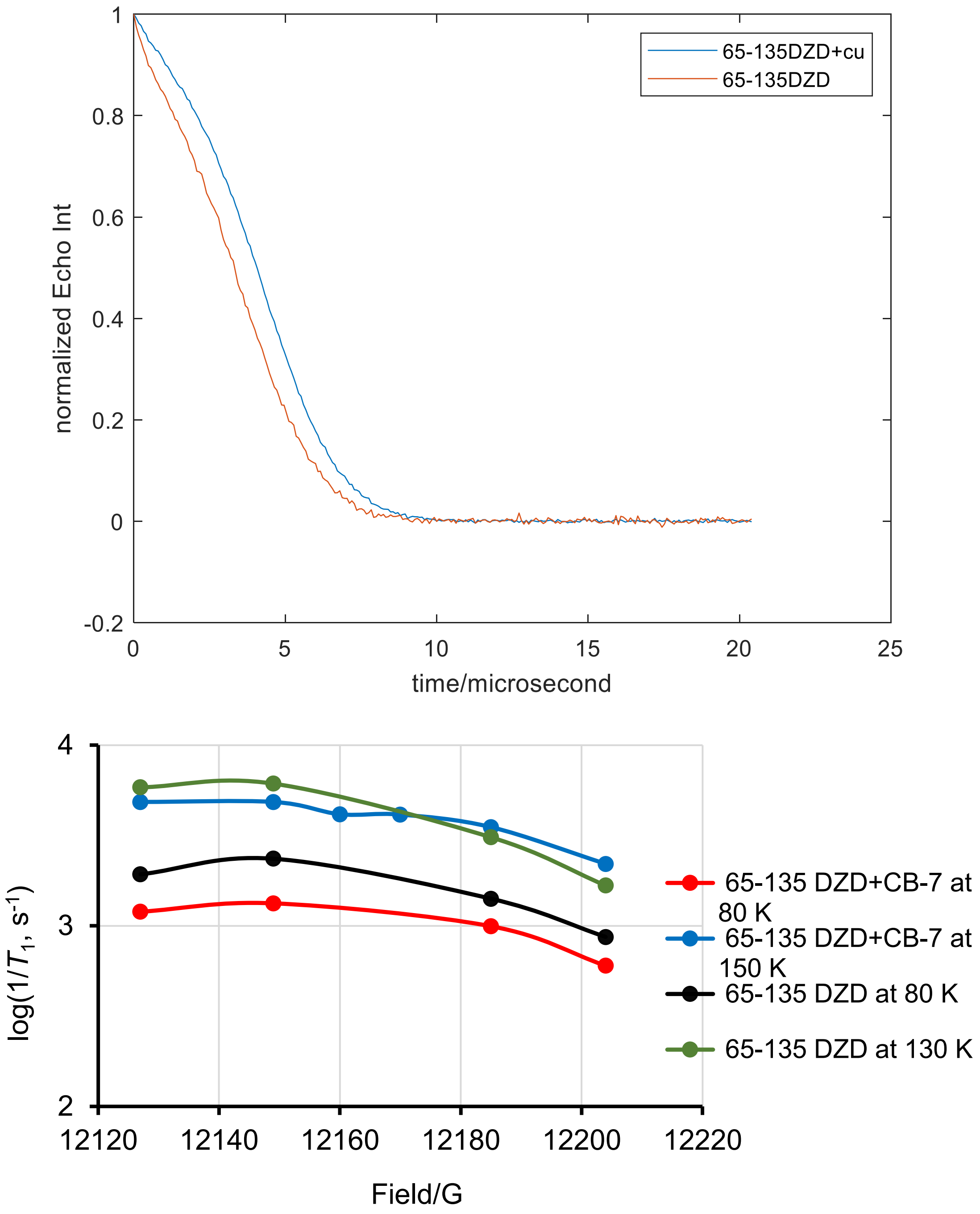

**Fig. S25.** *Top panel:* Normalized echo intensity for DZD T4L 65/135 in 30% glycerol in water without CB-7 (brown line, = cu) and with 1 mM CB-7 (blue line) at 80 K. The lines reflect the difference in $T_m$ = 4.20 μs and $T_m$ = 4.94 μs without CB-7 and with 1 mM CB-7. *Bottom panel:* Field and temperature dependence of $1/T_1$ for doubly spin labeled DZD-T4L 65/135 in 30% glycerol in water without and with 1 mM CB-7 at Q-band.

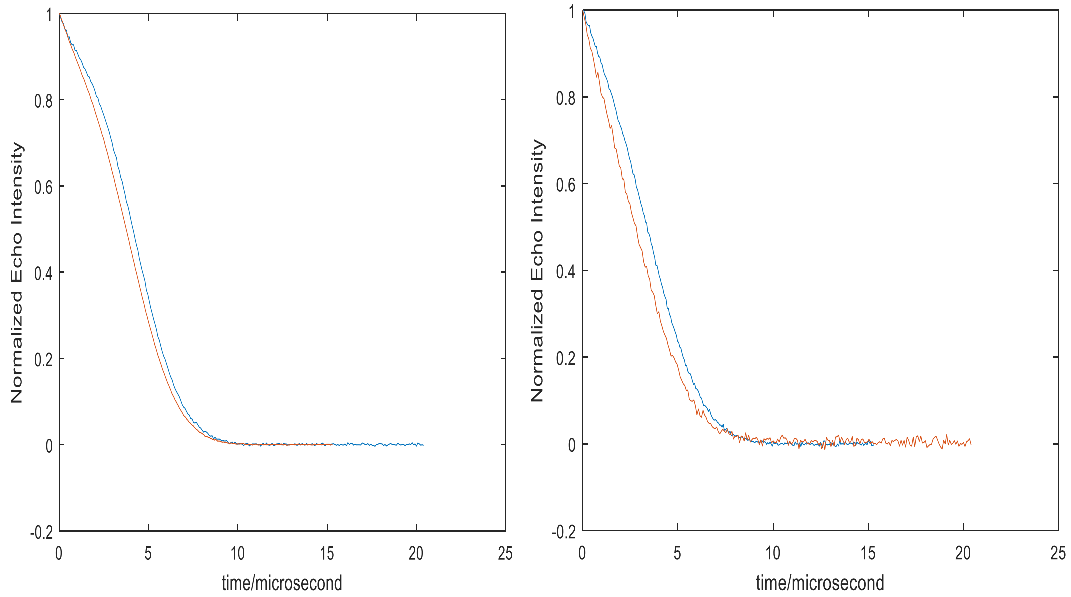

**Fig. S26.** Normalized echo intensity for DZD T4L 65/135 in 30% glycerol in water for β-CD complex (brown line) and for CB-7 complex (blue line) at 80 K (left) and at 150 K (right). The curves show the difference in echo amplitude at a particular time for samples with different values of $T_m$, which is particularly significant at longer times where DEER data are acquired .

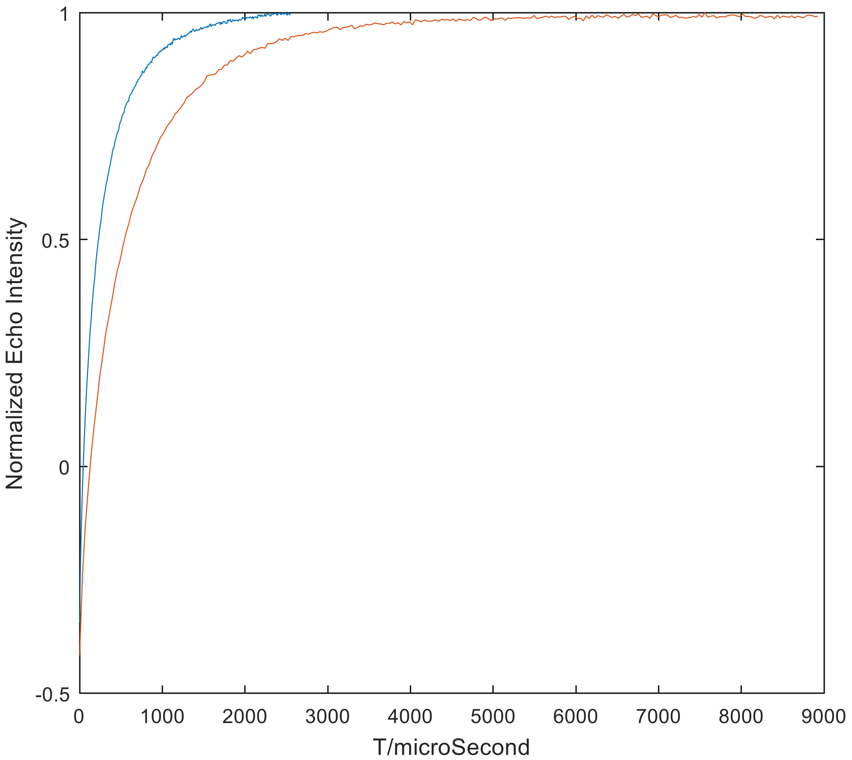

**Fig. S27.** Comparison of inversion recovery curves showing that, for DZD-T4L 65/135, $T_1$ is longer for the CB7 complex (brown line) than for the β-CD complex (blue line) at 80 K.

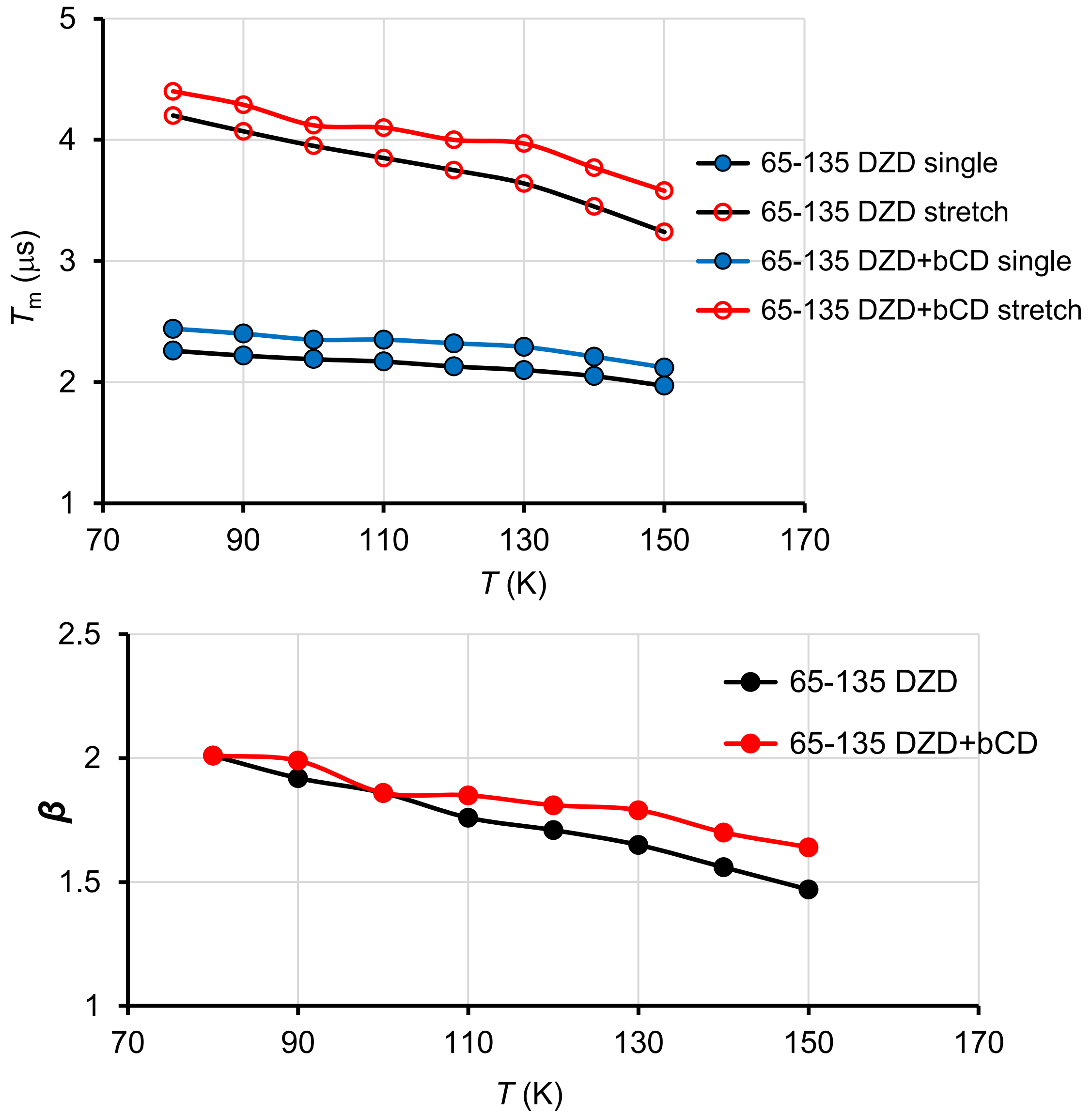

**Fig. S28.** *Top panel:* Comparison of $T_m$ temperature dependence for the stretched exponential vs. single exponential fits to two pulse echo decay for DZD-T4L 65/135 without and with β-CD. *Bottom panel:* As above for temperature dependence for the stretch parameter $\beta$.

**8. Spin labeling and DEER measurements.**

**Spin labeling of T4L.** Sample labeling was carried out as previously described (Table S7).[S1,S15,S16] Briefly, T4L mutants were expressed in K38 cells in Luria Broth (LB) and purified using cation exchange chromatography. Labeling of the mutants was initiated by adding an excess of spin-label and incubating for 2 h at room temperature, a second addition of spin-label was made, and the samples incubated overnight at 4 °C. The samples were then desalted and concentrated. Prior to DEER measurements (e.g., Figure 5, main text), CW EPR spectra for T4L mutants were obtained (Fig. S17). The spectra are consistent with increased rotational correlation times ($\tau_{rot}$) after addition of 1 mM CB-7 to DZD-T4L mutants and the absence of change in $\tau_{rot}$ for MTSL-T4L mutants.

**Table S7.** Summary of spin labeling of T4L mutants with IA-DZD.

| T4L mutant | Protein conc. [μM] | Nitroxide conc. [μM] | Labeling efficiency per Cys [%] | Notes |
|---|---|---|---|---|
| 65/80 | 47.8 | 35.4 | 37 | Same preparation, used for several experiments |
| 65/135 | 419 | 276 | 33 | Single preparation |
| 65 | 909 | 366 | 40 | Samples for X-ray and CW EPR studies |
| 135 | 812 | 310 | 38 | |

**DEER measurements.** The dipolar time evolution data was obtained at 83 or 150 K using a standard DEER four-pulse protocol, (π/2)mw1–τ1–(π)mw1–τ1–(π)mw2–τ2–(π)mw1–τ2–echo,[S17] on a Bruker 580 pulsed EPR spectrometer operating at Q-band frequency (33.9 GHz).

**Sensitivity of DEER measurements.** We estimate relative sensitivity of DEER measurements for T4L 65/80 and 65/135, doubly labeled with either MTSL or DZD with and without CB-7 at 83 – 200 K, using limiting concentration, LC, of spin label (Table 3, main text and Tables S8 and S9). Parameter LC is defined as minimum concentration of spin label in the sample providing signal-to-noise ratio, SNR = 10 in one scan, with SNR defined as maximum signal amplitude divided by the standard deviation of the residual noise. <u>The lower LC corresponds to higher sensitivity of DEER measurement.</u> Also, for T4L 65/80 at 83 – 200 K, we provide relative measurement time by partially optimizing shot repetition time (SRT), due to lowered electron spin relation time $T_1$ at higher temperature. At higher temperatures, the sensitivity of DEER measurement is getting progressively lower, mainly because decreased $T_m$.

**Table S8.** Summary of DEER data: distance distributions and relative sensitivity as limiting concentration of protein [LC].[a]

| $d_2$[b] (μs) | T4L | Spin label | Gly matrix | Temp (K) | *r* (σ)[c] [nm] | *r* (σ)[d] [nm] | [SL][e] (μM) | #scans | SNR | [LC] (μM) |
|---|---|---|---|---|---|---|---|---|---|---|
| 3.5 | 65/80 | DZD | | 83 | 2.44 (0.29) | 2.44 (0.29) | 27.4 | 150 | 518 | 6.48 |
| | | | + CB-7 | | 2.56 (0.19) | 2.62 (0.33) | 20.1 | | 751 | 3.28 |
| | | MTSL | | | 2.60 (0.34) | 2.60 (0.34) | 27.3 | | 231 | 14.5 |
| | | | + CB-7 | | 2.60 (0.35) | 2.60 (0.35) | 24.6 | | 233 | 12.9 |
| | 65/135 | DZD | | | 4.56 (0.47) | 4.56 (0.47) | 22.6 | | 718 | 3.86 |
| | | | + CB-7 | | 4.90 (0.19) | 4.76 (0.95) | 21.5 | | 1527 | 1.72 |
| | | MTSL | | | 4.45 (0.27) | 4.45 (0.27) | 19.8 | | 187 | 13.0 |
| | | | + CB-7 | | 4.51 (0.26) | 4.51 (0.26) | 19.9 | | 150 | 16.2 |
| 3.5 | 65/135[f] | DZD | | | 4.54 (0.53) | | 80 | 103 | 1560 | 5.2 |
| 3.2 | | | + β-CD[f] | | 4.90 (0.49) | | 55 | 173 | 2110 | 3.43 |
| 3.5 | | MTSL | | | 4.49 (0.23) | | 50 | 530 | 664 | 17.3 |
| 3.5 | | | + β-CD[f] | | 4.82 (0.25) | | | 600 | 1180 | 10.4 |
| 2.0 | 65/80 | DZD | + CB-7 | 83 | 2.62 (0.18) | 2.83 (0.76) | 20 | 9 | 1052 | 0.57 |
| | | | | 150 | 2.63 (0.26) | 2.72 (0.49) | | 18 | 734 | 1.15 |
| | | | | 170 | 2.58 (0.23) | 2.85 (0.90) | | 27 | 582 | 1.78 |
| | | | | 180 | 2.68 (0.69) | 2.68 (0.69) | | 45 | 394 | 3.40 |
| | | | | 190 | 2.61 (0.23) | 2.79 (0.59) | | 89 | 154 | 12.2 |
| 1.2 | 65/80 | DZD | + CB-7 | 200 | 2.73 (0.49) | 2.73 (0.49) | | 2250 | 249 | 38.1 |

[a] $[LC] = \{10*[SL]*(\#scans)^{1/2}\}/SNR$. [b] $d_2$ = echo acquisition time. [c] *r* (σ), average distance and breadth for the main components of the distance distribution. [d] *r* (σ), average distance and breadth for the overall distance distribution, including small components. [e] [SL] = concentration of spin label. [f] ref S1, gly+β-CD = with 10 mM β-CD.

**Table S9.** Relative sensitivity (LC) and relative measurement time for DZD-T4L 65/80 gly/CB-7 sample at 83 – 200 K. (LC data are the same as in Table 3, main text).

| Temp (K) | $d_2$ (ns) | SRT | no. scans | SNR | LC | Relative measurement time SRT*scans/9000 |
|---|---|---|---|---|---|---|
| 83 | 2000 | 1000 | 9 | 1052 | 0.57 | 1 |
| 150 | 2000 | 500 | 18 | 734 | 1.15 | 1 |
| 170 | 2000 | 300 | 27 | 582 | 1.78 | 0.9 |
| 180 | 2000 | 200 | 45 | 394 | 3.40 | 1 |
| 190 | 2000 | 200 | 89 | 154 | 12.2 | 1.98 |
| 200 | 1200 | 150 | 2250 | 249 | 38.1 | 37.5 |
| LC = (10*[SL]*√scans)/SNR ([SL] = 20 μM)<br><br>SRT = shot repetition time (μs) | | | | | | |

**DEER distance analysis by Gaussian fit.** Distance distributions were obtained from the time evolution data by assuming that the distance distribution can be approximated by a sum of Gaussians.[S18,S19] For all studied mutants, we are showing distance distribution with 95%-confidence bands using Gaussian fits in Figs. S29 and S31.[S20]

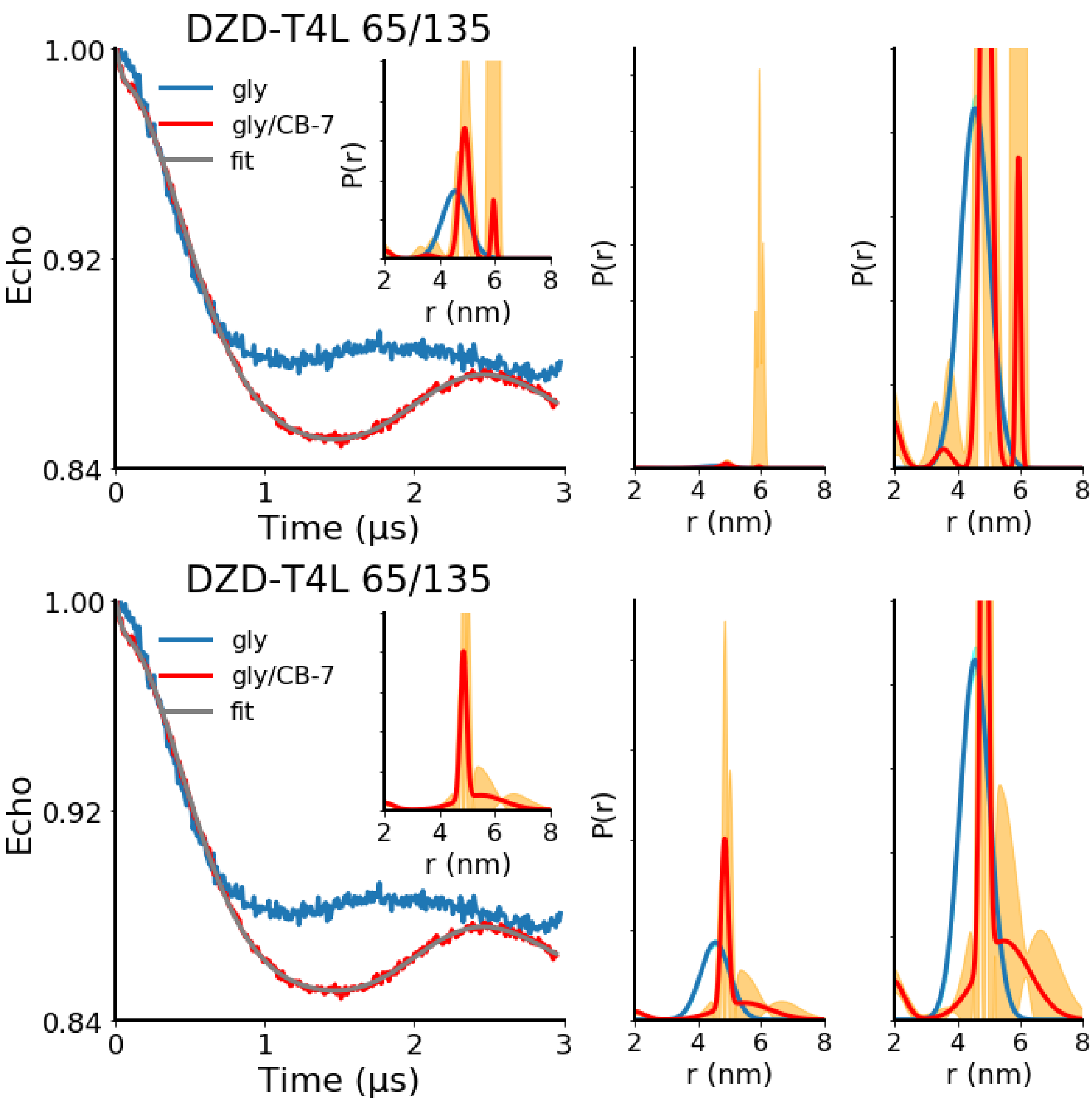

**Fig. S29.** DEER distance measurements on doubly spin labeled DZD-T4L 65/135 in 2:1 w/g matrix without CB-7 (gly) and with 1 mM CB-7 (gly/CB-7) at 83 K. The left side main plots correspond to normalized echo vs. time with fits, using a sum of Gaussians as the distance distribution. Inset plots and the right-side expansion plots correspond to Gaussian distance distributions with 95% confidence bands (navy blue and orange lines). Upper panels: data and fits as shown in Figure 5 (main text). Bottom panels: an alternative fit for gly/CB7, with the increased minimum width of the Gaussians in the distance distribution, is shown. This provides a fit with a broader long component but increases the 95% confidence bands for the main component.

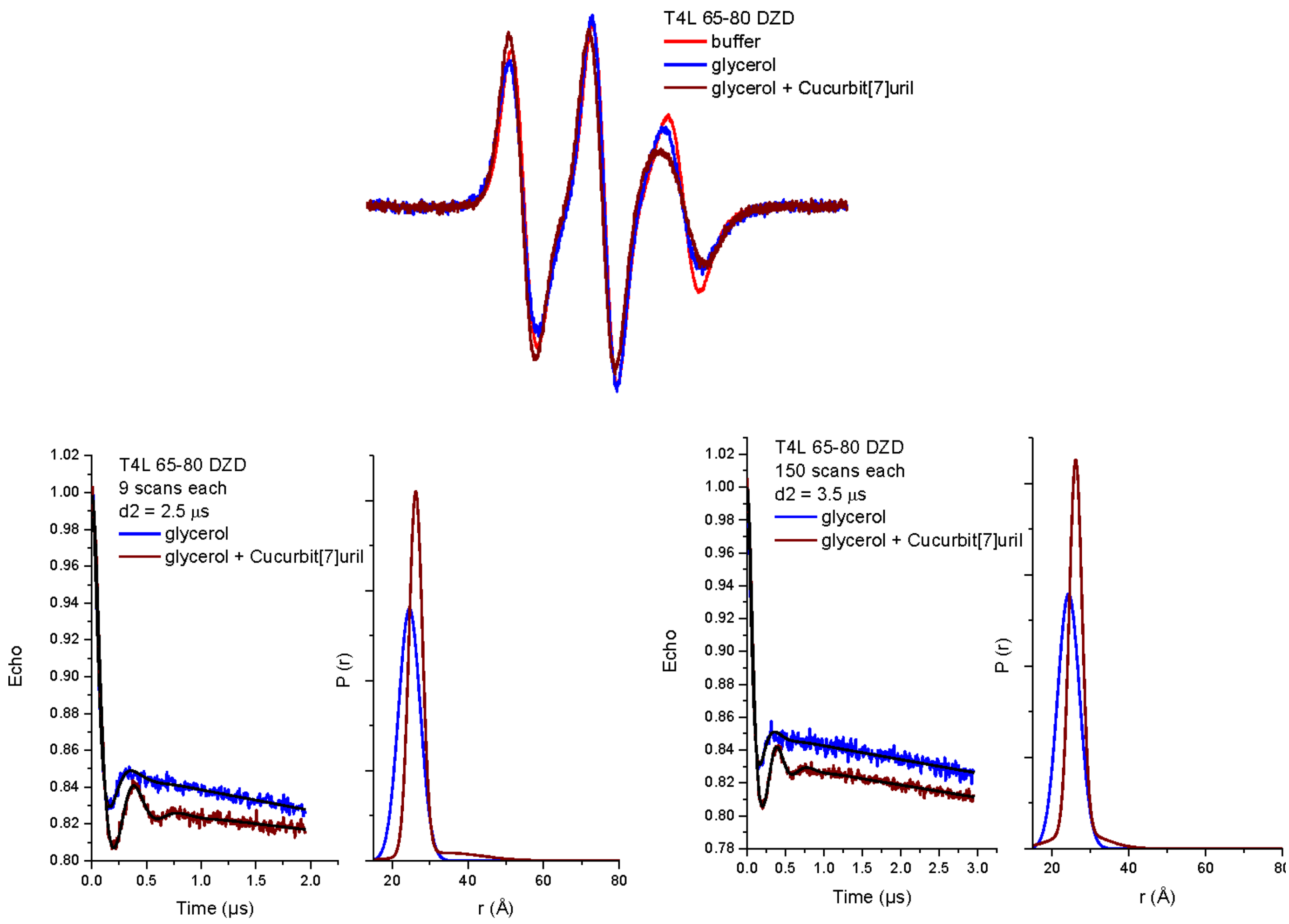

**Fig. S30.** Repeat sample of DZD-T4L 65/80: red line = 6 mM Tris buffer (pH = 7.2, 9 mM MOPS, and 50 mM NaCl), blue line = 2:1 w/g matrix, brown line = after addition of 1 mM CB-7. Top plots: CW EPR spectra at room temperature. The spectra are consistent with increased rotational correlation times ($\tau_{rot}$) after addition of glycerol and, in particular, CB-7. Bottom plots: DEER measurements at 83 K: echo delay $d_2$ = 2.5 (left plots) and $d_2$ = 3.5 (right plots) μs; normalized echo vs. time with fits, using a sum of Gaussians, and distance distributions, $P(r)$ vs $r$. The noise level is comparable for the data collected at 2.5 μs. Interestingly, the noise level is significantly better at the longer data collection time. This would suggest a significant increase in $T_m$, especially upon addition of CB-7.

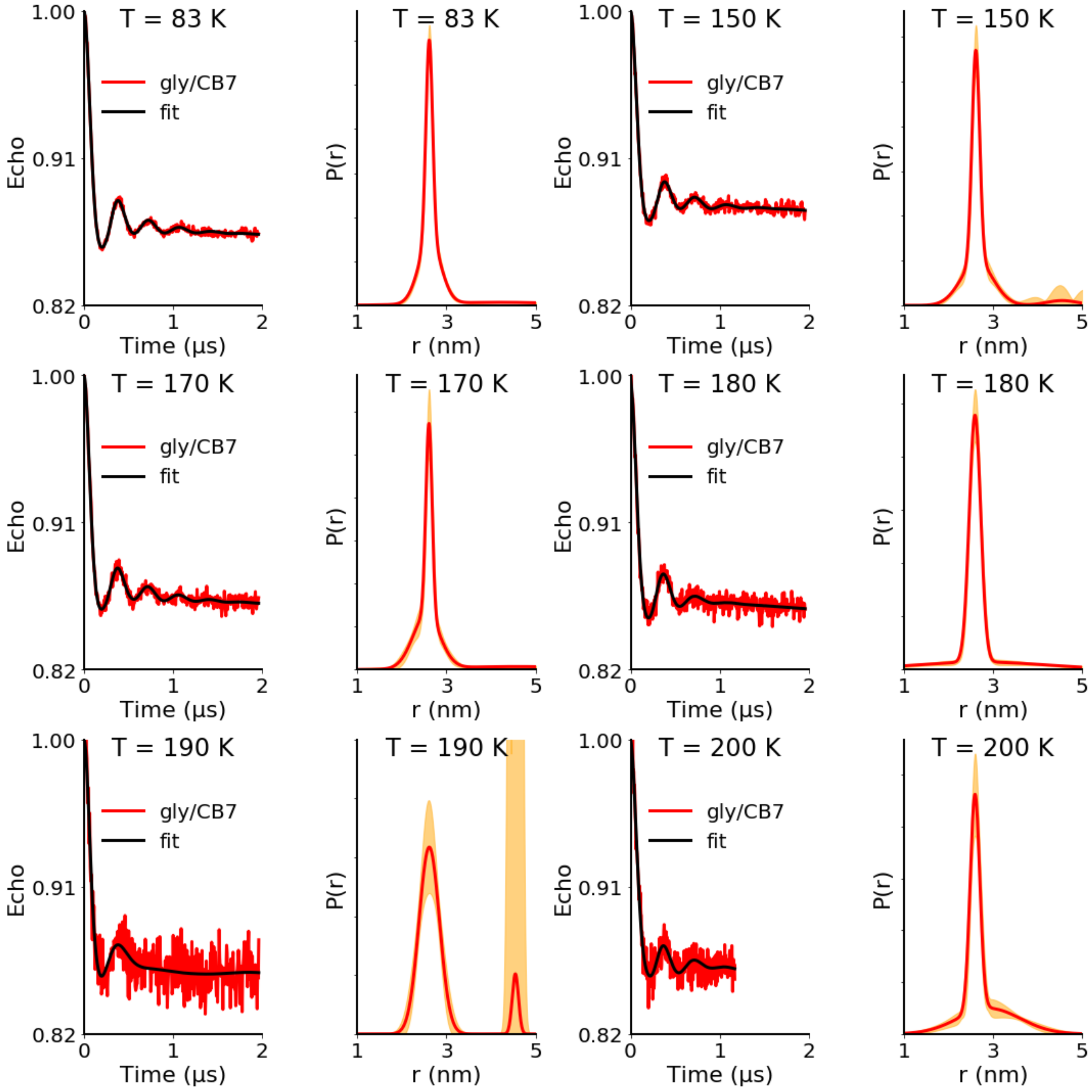

**Fig. S31.** Enlarged version of Figure 6 (main text). DEER distance measurements in the $T = 83 - 200$ K on doubly spin labeled DZD-T4L 65/80 in 2:1 water/glycerol with 1 mM CB-7. For each $T$, the left panels correspond to plots of normalized echo vs. time with fits where the distance distribution is a sum of Gaussians and the right panels display the corresponding distance distributions, including 95% confidence bands (range lines). All data are presented in Table 3 (main text) and Table S9. The data at $T$ = 190 K may not be adequately fit because of the relatively low SNR.

**9. Addition of 1 mM CB-7 does not affect the T4L structure.** It is well known that CB-7 forms strong guest-host complexes with *N*-terminal amino acids such as Phe ($K_a$ ~ $10^6$–$10^7$ $M^{-1}$).[S21,S22] The CB-7 recognition of N-terminal Phe was confirmed by X-ray crystallography of insulin@CB-7 complex.[S21] When N-terminal Phe was replaced with Glu no interaction between insulin and CB-7 was found.[S21] CB-7 selectively binds the epigenetic mark $N^\varepsilon$,$N^\varepsilon$,$N^\varepsilon$-trimethyllysine (LysMe3, $K_a = (1.8 \pm 0.6)\times 10^6$ $M^{-1}$) by 3500-fold over lysine ($(5.3 \pm 0.7) \times 10^2$ $M^{-1}$) in aqueous solution; the trend in $K_a$ is $LysMe_3 > LysMe_2 > LysMe > Lys$.[S23] Using *Ralstonia solanacearum lectin* (RSL), an extensively characterized and highly stable ~29 kDa trimer, possessing lysines in positions 25, 34, and 83, it was found by NMR spectroscopy that the native protein does not interact with CB-7.[S24] In particular K34Me2 mutant, with $N^\varepsilon$,$N^\varepsilon$-dimethyllysine at position 34, was found to possess some interaction with CB-7, which was also confirmed by crystallography; however, ITC studies failed to provide association constant,[S24] i.e., most likely $K_a \ll 10^3$ $M^{-1}$. Association constant, $K_a \sim 10^2$ $M^{-1}$ was estimated for complex of HSA with CB-7.[S25] Based on these results it is not likely that the addition of 1 mM CB-7 will affect the structure of T4L or its Cys mutants.
